# Magnetic organelle gene clusters enable bacterial growth and ATP homeostasis under oxidizing conditions

**DOI:** 10.64898/2026.11.06.764584

**Authors:** Mila Kojadinovic-Sirinelli, Mélissa Garry, Juan Wan, Agnès Dumont, Ramon Egli, Julie Carlut, Laurent Modolo, Daniel M. Chevrier, Lourdes Marcano, Fadi Choueikani, Christophe Gatel, Ingrid Marie Andersen, Ernesto Scoppola, Chenghao Li, Catherine Brutesco, Mathilde Nonnon, Emilie Gachon, Gaël Yvert, Aurèle Piazza, Damien Faivre, Arash Komeili, Matthieu Amor

## Abstract

Biological organisms can produce magnetic nanoparticles made of magnetite to detect the Earth’s magnetic field for orientation. The evolutionary mechanisms that led to the acquisition of this capacity remain unknown. The most ancient organisms known to perform magnetic orientation are magnetotactic bacteria. Under tight genetic control, they produce chains of magnetite nanoparticles in dedicated organelles to allow efficient navigation in aquatic environments containing molecular oxygen. Yet, these bacteria have been proposed to emerge before molecular oxygen became abundant on Earth, at a time when this navigation system could not have served its current function. Here, we show that gene clusters responsible of magnetite formation and organization enable resistance against hydrogen peroxide, a toxic oxygen-derived compound that was produced during the early stages of Earth’s oxygenation when iron oxidation consumed molecular oxygen. Characterization of the nanoparticle’s oxidation state supported the involvement of magnetite in resistance to hydrogen peroxide-induced oxidative stress. These findings are compatible with an early emergence of magnetotactic bacteria, before natural waters became oxygenated. We propose a conceptual framework to explain the origin of magnetic orientation in bacteria, and offer tools to identify the initiation of Earth’s oxygenation in a time-resolved manner.

---

Biological organisms have evolved the remarkable capacity to actively or passively detect the magnetic field of the Earth. This process, known as magnetoreception, often relies on magnetic nanoparticles made of magnetite (Fe^II^Fe^III^_2_O_4_) (1). Yet, when and how magnetite-based magnetoreception appeared remains unknown. The most ancient organisms known to possess this capability are magnetotactic bacteria (MTB), a diverse group of widespread aquatic microorganisms that synthesize chains of magnetite nanoparticles in a genetically controlled manner (2). In MTB, magnetite is contained within specialized organelles called magnetosomes that enable bacteria to align with the magnetic field of the Earth (2). This capacity optimizes navigation along oxygen (O_2_) gradients that develop in stratified waters between an oxidized surface and a more reduced deeper layer, and help MTB to efficiently escape highly oxygenated conditions (2, 3). MTB were proposed to have emerged at least 3 Ga, well before the so-called Great Oxidation Event that led to the oxygenation of Earth’s surface, dating back 2.5 Ga (4). Geochemical proxies also reported an oxygen-stratified ocean starting no earlier than 2.4 Ga (5, 6). MTB and their magnetosomes may thus have emerged long before O_2_ became abundant on Earth, at a time when such a magnetite-based navigation system would have no adaptive value. One possible explanation for this apparent contradiction is that magnetosomes originally served a distinct and potentially non-magnetic function, and were later adapted for magnetoreception (7). Consistent with this hypothesis, some modern MTB show inefficient magnetosome structures and magnetic properties that prevent them from proficiently detecting Earth’s magnetic field (8, 9), suggesting that magnetosomes had – or still have – an entirely different role. Several hypotheses have been proposed regarding additional functions of magnetosomes including resistance to hydrogen peroxide (H_2_O_2_) and other toxic elements (10–13), or more indirectly energy storage (14), but none could have been experimentally demonstrated. Here, we demonstrate that genetically induced fragmentation of magnetosome chains impairs the magnetoreception capacity of MTB. Yet, bacterial strains possessing these genes showed stronger resistance against H_2_O_2_, an oxidizing and toxic O_2_-derived compound generated on the early Earth when initially produced O_2_ was consumed almost immediately by oxidation of ferrous iron (15). The more oxidized state of nanoparticles in bacteria resisting H_2_O_2_ supported the involvement of magnetite to oxidative stress resistance. These findings legitimate an early origin of MTB prior to the Great Oxygenation Event.

## Genetically encoded modulation of magnetic properties

To explore alternative function(s), we began by comparing *Paramagnetospirillum magneticum* strain AMB-1 with mutants altered in magnetosome formation and organization. The genetic factors responsible for magnetosome formation are organized in gene clusters. In several MTB, these gene clusters are contained in a specific genomic region called the magnetosome island (MAI) (16). In addition to the MAI, some MTB such as AMB-1 also contain an additional gene cluster termed the magnetotaxis islet (MIS) (17). MIS genes, in conjunction with the MAI, place newly born magnetosome membranes between pre-existing magnetite-containing magnetosomes (18, 19). The result is chain fragmentation, with magnetite-containing magnetosomes spaced by nanoparticle-free vesicles, which causes a decrease of the capacity of cells to align with external magnetic fields (Figs. 1A-F, Fig. S1) (18). It also increases the distance between two magnetosome vesicles and elongates nanoparticles (Figs. S2,3) (18). We investigated the contribution of MIS genes to the magnetic behavior of AMB-1 by measuring the coefficient of magnetism (C_mag_) in wild-type bacteria (fragmented chains) and a mutant strain from which all MIS genes were deleted (ΔMIS, continuous chains) using a differential spectrophotometric assay when a magnet is placed either vertically or horizontally close to a bacterial suspension (see Methods for details). Consistent with our previous findings (18), C_mag_ values were 35 % higher in AMB-1 populations lacking MIS genes (*P* = 0.001) (Fig. 1H). This shows that MIS-induced fragmentation of magnetosome chains reduces the ability of AMB-1 to align with external magnetic fields. In contrast, deletion of MAI (ΔMAI strain) completely abolishes magnetosome formation (Fig. 1G, Fig. S1) (16). To resolve the mechanisms enhancing the magnetic response of ΔMIS bacteria, we characterized the magnetosome chains of wild-type and ΔMIS AMB-1. X-ray diffraction identified magnetite as the sole contributor to the magnetization of both strains (Fig. 1I). They produced comparable magnetite content, although slightly lower in ΔMIS AMB-1, which exhibited an 8% reduction in magnetite mass normalized to bacterial dry weight (Figs. 1L,M) that is also compatible with the decrease in magnetite size in this strain (8.5 %) (Table S1). FORC (first-order reversal curves) diagrams were then acquired to further elucidate their magnetic properties. They are interpretable as the joint distributions of hysterons (elemental squared hysteresis loops) with half-width *B*_c_ (coercivity) and bias field *B*_u_ for the fraction of nanoparticles that are sufficiently large for their magnetization to be stable within the measurement time (20). FORC diagrams of wild-type and ΔMIS AMB-1 are similar, with typical signatures of magnetosome chains evidenced by the central ridge along *B*_u_ = 0, consistent with switching between two-state magnetic configurations where all magnetic moments are parallel or antiparallel to the chain axis (Figs. 1L,M). This was also confirmed at the nanoparticle scale with mapping of the magnetic flux in magnetosome chains, showing that magnetic moments of magnetite were nearly parallel to the chain or chain fragment axis (Figs. 1A-F, Fig. S2) (9). However, FORC diagram details revealed important departures from idealized gap-free magnetosome chains, which require a finite minimum field to be switched owing to the chain-axis anisotropy and the strong inter-crystal magnetic coupling. The coercivity distributions were not unimodal and included a low-coercivity (LC) component truncated at *B*_c_ = 0 (Fig. S4, Table S2). The other high-coercivity (HC) component, spanning ~10 mT to ~70 mT, closely mathced numerical simulations of intact, gap-free chains of equidimensional magnetosomes (9). Its lower median fields in ΔMIS AMB-1 (~29 mT) than in wild-type cells (~35 mT) is consistent with the lower particle elongation in the mutant (Figs. S3,4, Tables S1,2) (21). The LC component was attributed to chains with minimum gaps of ~4 missing magnetite long enough to decouple the magnetic polarity of individual fragments (Supporting Notes), which are more frequent in wild-type cells (Fig. S1), as shown by the larger LC contribution (~44%), compared to ΔMIS cells (~24%). Smaller gaps (1-2 missing magnetite) were also detectable, but contributed equally small signals in both strains. We thus hypothesized that chain fragments with opposite native polarities develop within a single wild-type cells (22). The magnetic moment of the same cell suspension was measured before and after exposure to large magnetic fields (200 mT) above magnetite coercivity. Application of a strong external field imparted the same polarity to all fragments, increasing the total cellular magnetic moment of wild-type AMB-1 by 40 %, which is consistent with the increase of C_mag_ (35 %) in the ΔMIS strain (Fig. 1J). A drastically different behavior was observed for ΔMIS AMB-1, where the application of a 200-mT field generated a 0 – 10 % increase of the cellular magnetization (Fig. 1K, Fig. S5). Thus, most particles in continuous chains have their magnetic moments aligned to maximize the magnetic contribution of each nanoparticle. Impaired AMB-1 magnetization induced by MIS may be evolutionarily neutral if fragmented chains maintain magnetic moments sufficiently strong to sustain magnetoreception in natural habitats, or even be beneficial to avoid obstacles more efficiently (23). Alternatively, the fragmented chain phenotype may represent an evolutionary trade-off between magnetoreception and an unidentified function. We previously proposed that MIS-induced chain fragmentation facilitates segregation of magnetosomes to daughter cells during cell division (18). Here, we investigated additional, potentially non-magnetic advantages provided by MIS, but also by MAI.

**Fig 1.**
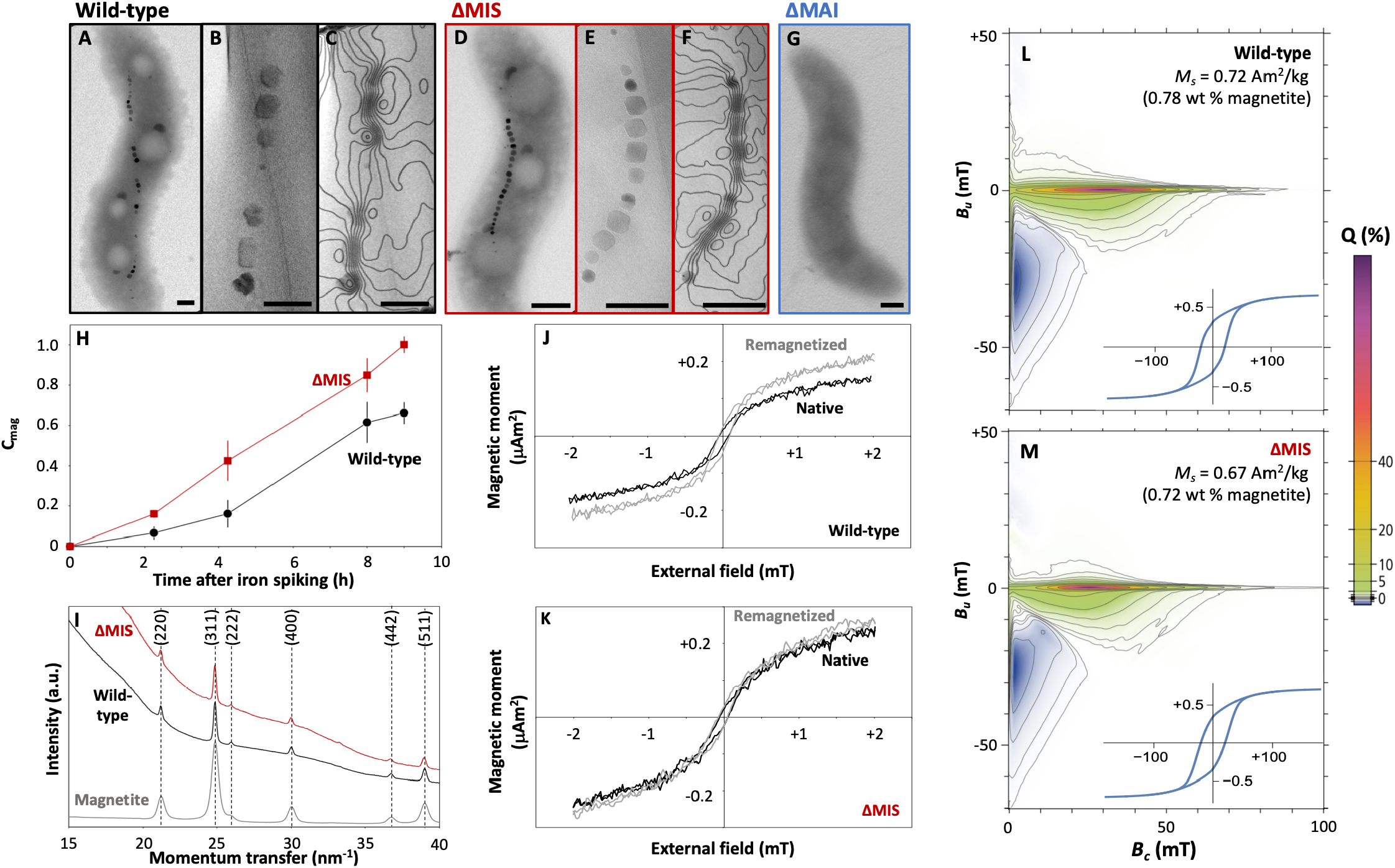
(A, D, G) Electron microscopy observations of (A) wild-type, (D) ΔMIS and (G) ΔMAI AMB-1. (B, C, E, F) Magnetic phase contours of magnetosome chains determined by off-axis electron holography in (B, C) wild-type and (E, F) ΔMIS AMB-1. Scale bars = 200 nm. (H) C_mag_ of wild-type, ΔMIS and ΔMAI cultures (mean values of three replicates for each strain). Bacteria were cultivated without iron to prevent magnetite formation until a bacterial concentration corresponding to an optical density (600 nm) of 0.01 was reached. Iron was then spiked to the growth media to monitor C_mag_. (I) X-ray diffraction analyses of magnetite reference, wild-type and ΔMIS AMB-1 cells. The scattered intensity was calculated as a function of the momentum transfer. (J, K) Magnetic moment of (J) wild-type and (K) ΔMIS cultures in their native states (black curves) and after remagnetization (exposure to an external field of 200 mT – grey curves). (L, M) FORC diagrams of wild-type (L) and ΔMIS (M) AMB-1 cultures with quantile contours (Q). The mass of magnetite (wt %) was calculated from saturation magnetization (*M*_*s*_) normalized to the mass of dried cells. Insets show the major hysteresis loop, which is the envelope of all measurements.

### Altered bacterial physiology in ΔMIS and ΔMAI strains

We first tested two parameters: growth and intracellular levels of adenosine triphosphate (ATP). Wild-type, ΔMIS and ΔMAI strains were cultivated in 1.5-mL Eppendorf tubes under standard conditions, in which bacteria are inoculated at ambient O_2_ conditions. At a given time point, three tubes corresponding to the three strains were sacrificed for simultaneous measurement of bacterial concentrations, cell morphology, and ATP content. For each strain, three independent biological replicates initiated from separate inocula were prepared and cultivated either in a zero-magnetic field environment or exposed to a constant, unidirectional 2-mT field (i.e., 40 times higher than the Earth’s magnetic field) (Fig. S6). Results obtained in the two field conditions were identical within analytical uncertainty (Figs. S7-9), showing that exposure to external magnetic fields in our experimental conditions had no observable effects. They are thus treated as replicates (n = 6 replicates from three independent inocula for each strain) hereafter. Wild-type AMB-1 yielded the highest bacterial densities in cultures ending exponential phase (44 h) at 2.08 ± 0.40 × 10^8^ cells per mL, while mutant bacteria showed lower densities at 0.79 ± 0.01× 10^8^ (ΔMIS, *P* = 1.03× 10^-4^) and 0.97 ± 0.07× 10^8^ (ΔMAI, *P* = 3.16× 10^-4^) cells per mL (Fig. 2A, Figs. S7-9). We observed elongated cells in the two populations of mutant bacteria initiating the stationary phase (46 h, Fig. 2B-E). Intracellular ATP levels were also elevated in the two mutant strains, up to 2.91 ± 0.37 (ΔMIS, *P* = 6.00× 10^-7^) and 2.77 ± 0.52 times (ΔMAI, *P* = 1.60× 10^-6^) higher than in wild-type bacteria (Fig. 2F). To better elucidate the differential roles of MIS and MAI, AMB-1 growth was characterized at high temporal resolution in 96-well plates. Continuous monitoring using this experimental setup revealed a sharp and transient two-fold drop of optical density (between 55 and 80 h of culture) in all ΔMAI cultures (Fig. 2G, Fig. S10). Wild-type and ΔMIS bacteria did not exhibit such a drop with only some ΔMIS cultures (5 out of 12 replicates) reaching a maximum drop of 30 % in optical density (Fig. 2G, Fig. S10). Finally, higher doubling times were observed in ΔMAI AMB-1 relative to wild-type, increasing by 28 % (*P* = 7.54× 10^-4^) (Fig. 1H, Fig. S11). Overall, these data demonstrate that deletion of MIS, and more pronouncedly of MAI, affected both growth and ATP levels in AMB-1.

**Fig 2.**
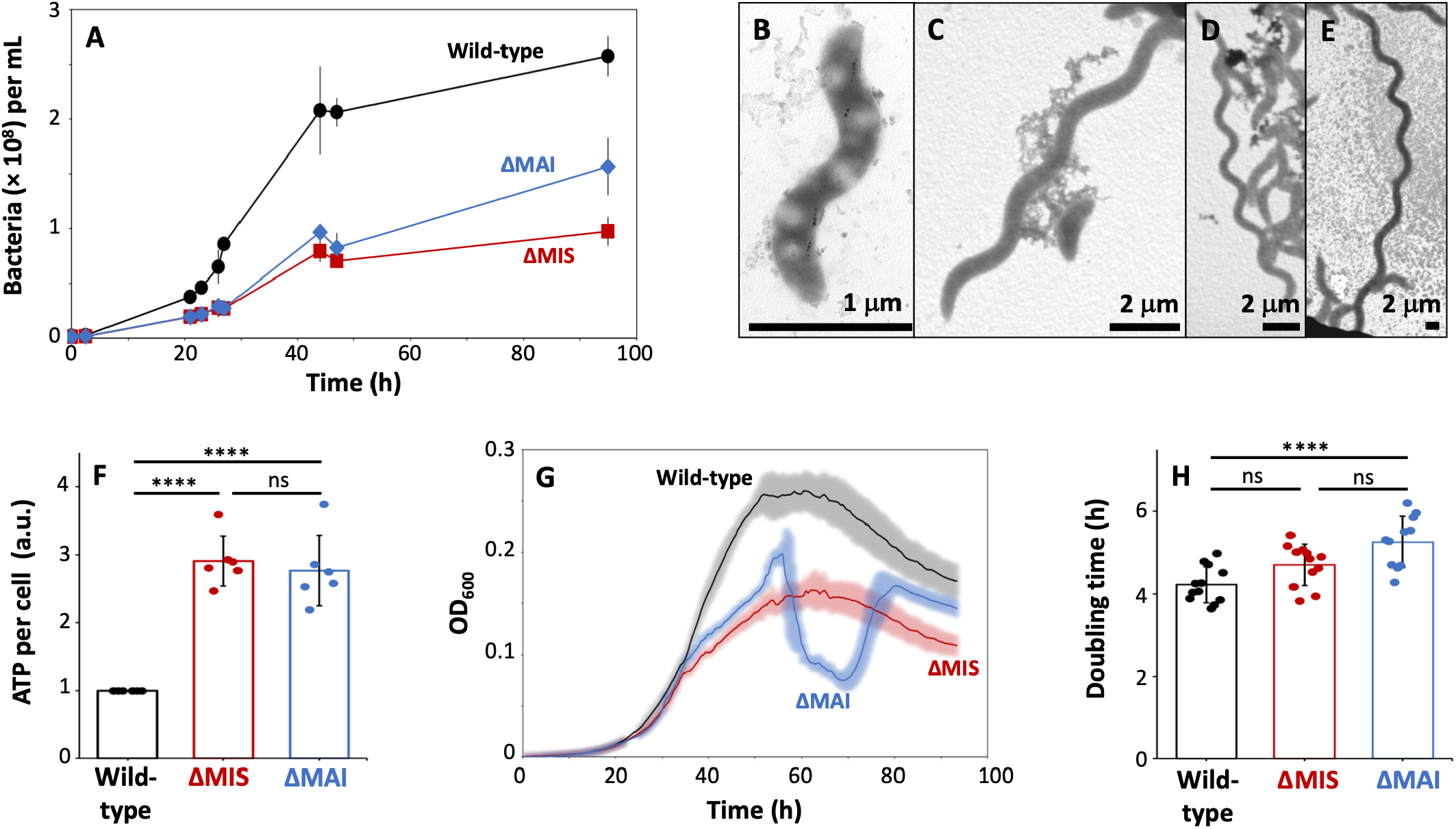
(A) Growth of wild-type (black curves - circles), ΔMIS (red curves – squares) and ΔMAI (blue curves – diamonds) AMB-1 in Eppendorf tubes measured using flow cytometry. (B) Electron micrograph of a wild-type cell. (C, D, E) Electron micrographs of filamentous cells observed in ΔMIS and ΔMAI cultures. (F) ATP content per cell in the three AMB-1 strains during the exponential growth phase, normalized to wild-type. (G) Growth of the three AMB-1 strains cultivated in 96-well plates and monitored with optical density at 600 nm (OD_600_). (H) Doubling time of AMB-1 populations calculated during the exponential phase from growth in 96-well plates. All data in (A) and (F) correspond to the mean value of six replicates (three exposed to a 2-mT field and three cultivated in a zero-field environment – see main text – ±1SD). Data in (G) and (H) correspond to the mean values of six replicates, with shaded areas illustrating standard deviation. ****: *P* < 0.0001, ns: not significant. See Supporting Information for detailed data.

### Deletion of magnetosome gene clusters alters global gene expression

To better elucidate how MIS and MAI deletion affected bacterial physiology, we characterized gene transcription profiles (whole-genome RNA-sequencing) of the three AMB-1 strains (n = 3 replicates for each strain) harvested during exponential growth phase (40 h of culture). Principal component analyses (PCA) revealed that deletion of MIS and MAI resulted in distinct transcriptomic profiles, with wild-type, ΔMIS and ΔMAI forming three separate clusters (Fig. S12). PC1, which accounted for 71% of the total variance, separated ΔMAI from the other two strains, whereas PC2 (17 % of the total variance) separated ΔMIS from the wild-type. We identified 278 (ΔMIS) and 728 (ΔMAI) differentially expressed genes (DEG) with a fold change (FC) of at least |log_2_FC| > 1, relative to wild-type (false-discovery rate < 0.05) (Fig. S13). Genes involved in the flagella assembly (*flg*) and the synthesis of ribosomal proteins (*rpl*) were among the most up- and down-regulated ones, respectively, in both mutant strains (Fig. S14, Table S3). Down-regulation of genes involved in anabolic pathways could explain the lower bacterial densities and higher ATP levels reached in the two mutant cultures, as well as the higher doubling time of ΔMAI AMB-1. Consistent with this interpretation, we also identified down-regulation of peptidoglycan synthesis genes in both mutants, particularly in ΔMAI AMB-1, and those of cell cycle and fatty acids synthesis in ΔMAI only (Fig. 3, Table S4). Genes related to nucleotide synthesis were also down-regulated in ΔMIS AMB-1, but up-regulated in the ΔMAI strain (Fig. 3). In sum, gene expression profiles were compatible with altered growth and ATP levels observed in the two mutant strains. Although several factors could contribute to these phenotypes, oxidative stress induced by hydrogen peroxide (H_2_O_2_), a naturally occurring compound, is consistent with all of them and may provide a common explanation. In non-magnetotactic bacteria, it was shown to reduce cell proliferation and increase ATP levels (24, 25), trigger cell elongation in populations initiating the stationary phase (26), increase the expression of flagellar genes (27), and down-regulate genes involved in anabolic pathways including the synthesis of ribosomal proteins (28). Similar to ΔMAI cultures (Fig. 2G), a sharp and transient drop in bacterial densities corresponding to H_2_O_2_-induced cell lysis was reported in populations of *Escherichia coli* lacking sigma factors (RpoE and RpoS), which are key regulators of the stress response that maintain membrane homeostasis and cell viability under prolonged oxidative stress (29, 30). Consistently, gene expression of many pathways involved in oxidative stress response was up-regulated in the two mutant AMB-1 strains (Fig. 3). They included glutathione peroxidases (*gpx*) and glutathione reductases (*gor*), superoxide dismutases (*sod*), and proteases (*ftsH*) that degrade damaged proteins in the context of oxidative stress (31). Notably, expression of the redox-sensitive transcriptional regulator involved in oxidative stress response (*oxyR*-like) was down-regulated in ΔMIS and ΔMAI bacteria. Finally, the housekeeping (*rpoD*) and membrane homeostasis (*rpoE*) sigma factors were down-regulated in ΔMAI but not in ΔMIS. These results suggest that ΔMIS and ΔMAI actively experienced H_2_O_2_-induced oxidative stress. We thus asked whether MIS and MAI provided resistance against H_2_O_2_.

**Fig 3.**
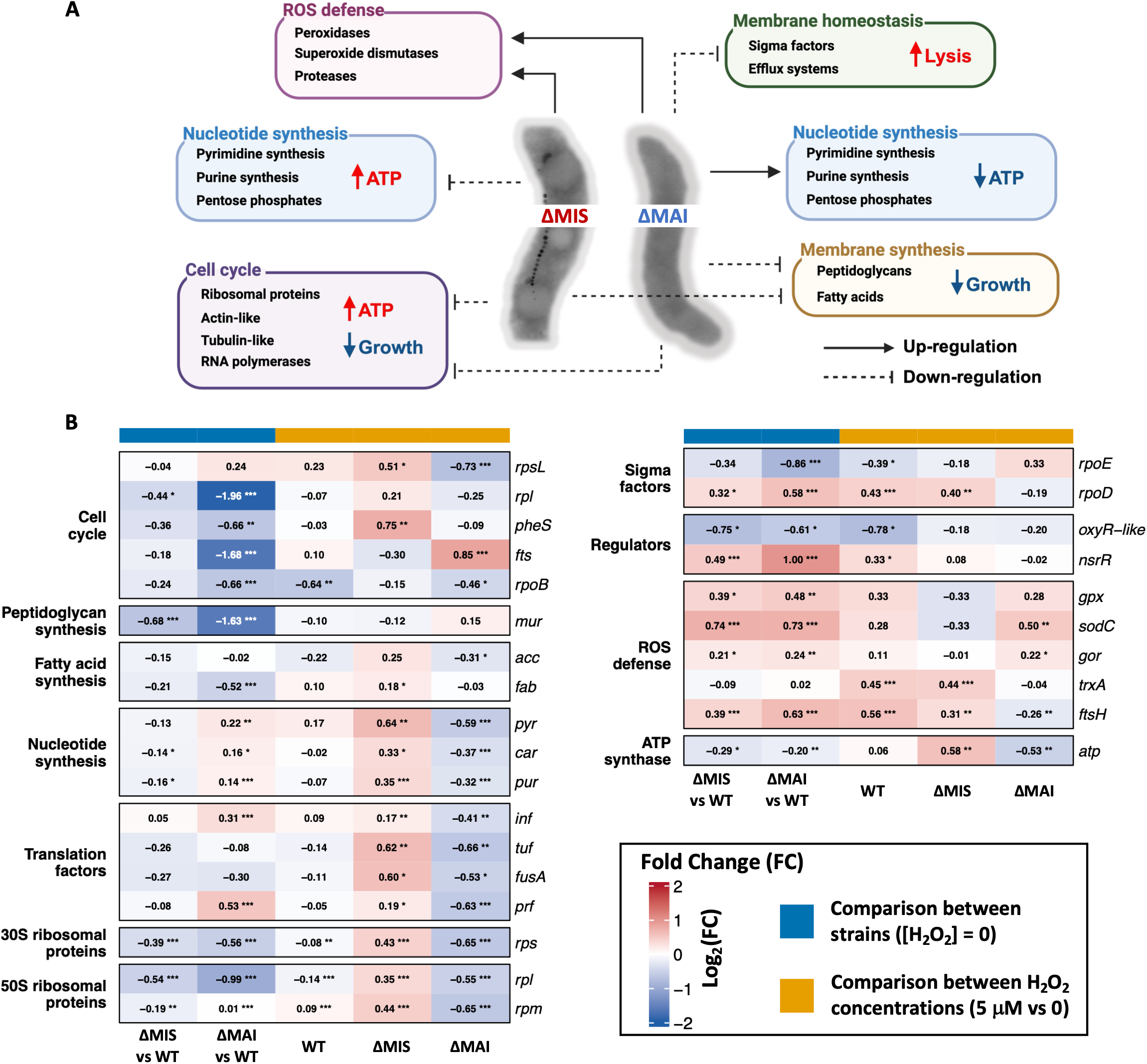
(A) Summary of metabolic pathways differentially regulated in ΔMIS and ΔMAI AMB-1 (relative to wild-type) cultivated without added H_2_O_2_, and their impact on growth, ATP levels and cell lysis. (B) Heatmaps showing differential gene expression involved in anabolic pathways, the oxidative stress response, and ATP production. WT: wild-type AMB-1. Numbers correspond to Log_2_(FC) values. *: P < 0.05, **, *P* < 0.01, ***: *P* < 0.001, ****: *P* < 0.0001. [H_2_O_2_] = 0 refers to standard conditions with no added H_2_O_2_.

### Enhanced H_**2**_**O**_**2**_ tolerance in wild-type AMB-1

To demonstrate that MIS and MAI provided protection against H_2_O_2_, we cultivated AMB-1 under increasing conditions of added H_2_O_2_ (0, 2.5, 5,× 10 µM) using a microplate reader.

Hydrogen peroxide concentrations in natural aquatic environments are typically in the range of 0 – 5 µM, with rare episodic peaks above 5 µM (32). Experimental conditions were thus selected to mimic environments that harbor MTB, and to simulate episodes of elevated H_2_O_2_. Results demonstrated that only wild-type bacteria were able to grow under H_2_O_2_ concentrations above 5 µM (Figs. 4A-C). RNA sequencing of wild-type AMB-1 cultivated with 5 µM of H_2_O_2_ revealed differential transcriptional response: it led to 314 DEG with |log_2_FC| > 1, most of which corresponding to down-regulated genes (Fig. S13). The transcriptional regulator *oxyR*-like was among down-regulated genes, which demonstrated that deletion of MIS and MAI had comparable effect than exposure to H_2_O_2_ in terms of *oxyR*-like expression (Fig. 3B). In addition, H_2_O_2_ induced up-regulation of the nitrosative stress regulator *nsrR, rpoD, ftsH* and genes encoding for thioreductases (*trxA*) (Fig. 3B). Exposure to H_2_O_2_ induced a more limited transcriptional response in ΔMIS (28 DEG) and ΔMAI (90 DEG) (Fig. S13). Remarkably, the number of DEG between the untreated mutants and the H_2_O_2_-treated wild-type was 35 % (ΔMIS; 40% when excluding MIS genes) and 38 % (ΔMAI; 45% when excluding MAI genes) lower than between the untreated mutants and the untreated wild-type (Fig. S15). The positively correlated |log_2_FC| of the two comparisons pointed to the shifted transcriptome of wild-type towards the mutant strains under H_2_O_2_ exposure, with Spearman’s ρ = 0.28 and 0.33 for ΔMIS and ΔMAI, respectively (adjusted *P* < 0.002 in both cases). When excluding MIS and MAI genes, ρ values increased to 0.30 (ΔMIS, adjusted *P* < 0.002) and 0.35 (ΔMAI, adjusted *P* < 0.002). Specifically, the basal profiles (no H_2_O_2_ treatment) in both ΔMIS and ΔMAI mutants were significantly and strongly coordinated with the wild-type response to H_2_O_2_ in the oxidative stress-related genes identified above, with a stronger and more consistent signal in ΔMAI (ρ = 0.708, adjusted *P* = 0.011) than in ΔMIS (ρ = 0.644, adjusted *P* = 0.011). These findings indicated that the coordination is specific to oxidative stress-related genes rather than reflecting a genome-wide association between H_2_O_2_ response and mutant versus wild-type differences. The two mutants also showed contrasting transcriptional response under increased oxidative stress regarding anabolic pathways (cell cycle and division, nucleotide and amino acid synthesis, transcription factors) and ATP synthase-related genes: they were up-regulated in ΔMIS bacteria, but down-regulated in ΔMAI AMB-1 (Fig. 3). These findings showed that ΔMIS bacteria can maintain an active metabolic state under increased oxidative stress at 5 µM of H_2_O_2_, whereas ΔMAI bacteria adopted a survival strategy characterized by reduced metabolism. They further pointed to a more severe oxidative stress in ΔMAI AMB-1.

**Fig 4.**
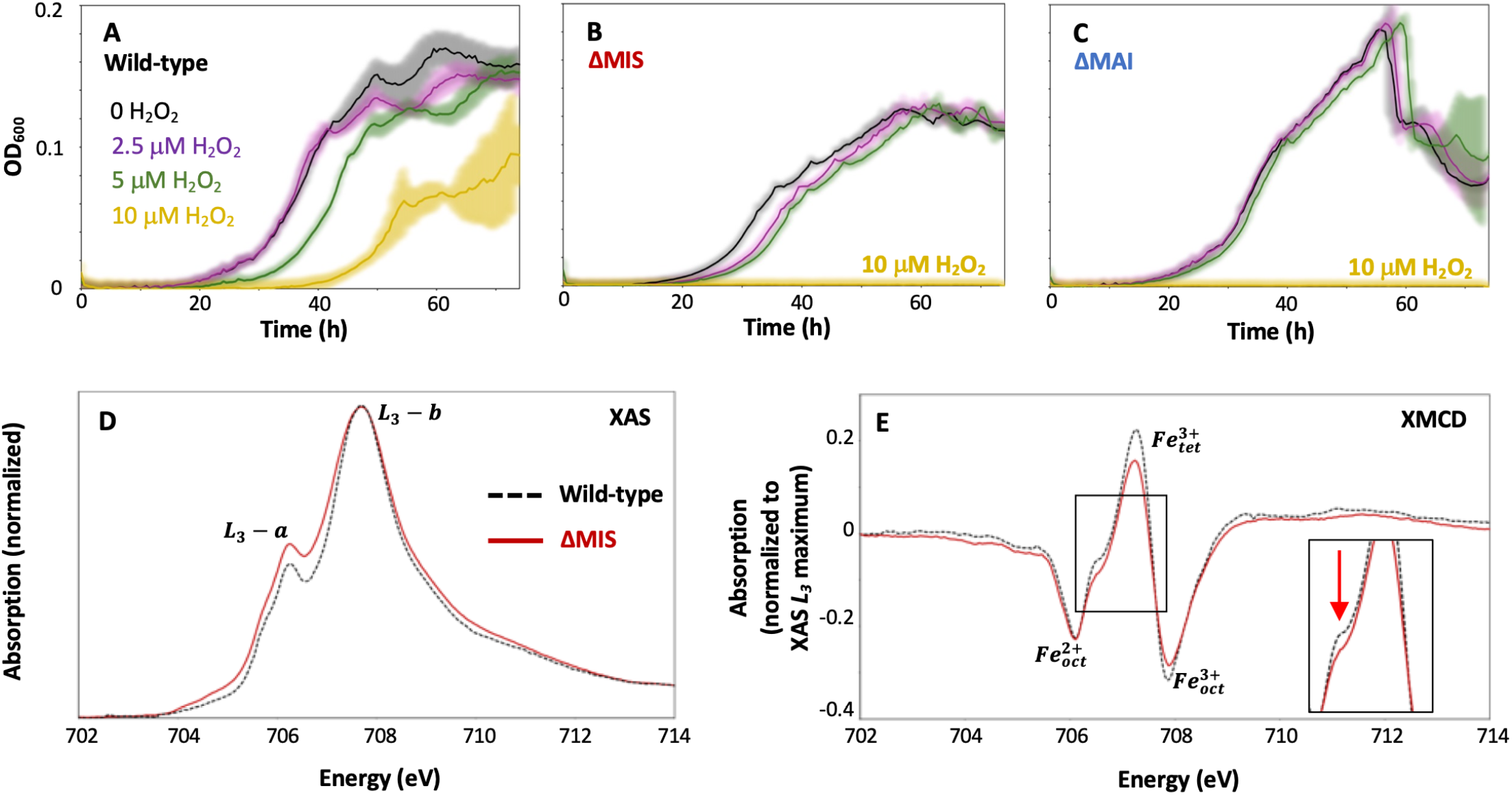
(A-C) growth of (A) wild-type, (B) ΔMIS and (C) ΔMAI AMB-1 measured from optical density at 600 nm (OD_600_) in media supplemented with H_2_O_2_. Shaded areas correspond to standard deviation obtained from three replicates. (D) Fe *L*_*3*_-edge X-ray absorption spectra (XAS) of wild-type (dotted black line) and ΔMIS (red line) AMB-1. (E) Fe *L*_*3*_-edge XMCD spectra of wild-type (dotted black line) and ΔMIS (red line) AMB-1.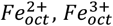 and 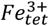 octahedral Fe^2+^, octahedral Fe^3+^ and tetrahedral Fe^3+^, respectively. Spectra at the full *L*_*2,3*_-edge range are provided in Supporting Information. Inset in (E) highlights the XMCD feature at 706.6 eV indicated by the red arrow.

### Antioxidant properties of magnetosomes

To elucidate the mechanisms enabling H_2_O_2_ resistance in wild-type bacteria, we searched for genes involved in the oxidative stress response in the MIS and MAI. They contain 25 (MIS) and 75 (MAI) genes with no known functions regarding magnetosome formation (Tables S5,6). No gene involved in the oxidative stress response could be identified from HMM (Hidden Markov Model) analysis, and all of these additional genes with an identified HMM product possessed between 1 and 60 predicted paralogs outside of these two gene clusters. We thus hypothesized that magnetosomes and their magnetite actively contributed to this protective function. Magnetite can efficiently eliminate H_2_O_2_ by reducing it into H_2_O via Fenton reactions, which leads to its oxidation (33, 34). We performed characterization of iron oxidation state in bacteria and their nanoparticles with X-ray absorption spectroscopy (XAS) at the Fe *L*_*2*_,_*3*_-edge. Given the enhanced sensitivity of ΔMIS bacteria to oxidative stress under standard conditions (no artificially added H_2_O_2_), we predicted a more limited oxidation of magnetite in ΔMIS samples. Consistent with previous findings (35), linear combination fitting identified 35 % magnetite (Fe^II^Fe_2III_O_4_), 57 % maghemite (γ-Fe^III^ O – which represents oxidized magnetite) and 8 % ferrihydrite (Fe^III^ O •0.5H O) in wild-type AMB-1 (Figs. S16-18). XAS spectra of ΔMIS bacteria pointed to a more reduced state of iron with 86 % magnetite and 14 % ferrihydrite, as evidenced by the higher *L*_*3*_*-a*/*L*_*3*_*-b* peak ratio (2.05 and 1.78 for wild-type and ΔMIS, respectively) and the shifted rising *L*_*3*_*-a* peak towards lower energy (Fig. 4D, Figs. S16-18, Table S7). Whereas XAS probes all iron phases present in the cell, X-ray magnetic circular dichroism (XMCD) is selectively sensitive to the magnetically ordered spinel phase and thus provides the iron valence and site distribution (tetrahedral – tet, or octahedral – oct) of the magnetite/maghemite nanoparticles specifically. XMCD evidenced a deficit of 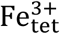 and 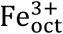 relative to 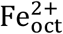 in ΔMIS (Fig. 4E, Figs. S16, 17) (36), which was confirmed by linear-combination analysis of the XMCD spectra using theoretical peak contribution (Supporting Notes, Figs. S19). In addition, wild-type samples showed a more pronounced shoulder at 706.6 eV, a feature observed in maghemite and ferrihydrite but not in magnetite (Fig. 4E, Fig. S17). Given the low relative weight of ferrihydrite evidenced from XAS, and the low magnetic contrast of ferrihydrite relative to iron oxides, this observation is also compatible with a more oxidized magnetite in wild-type bacteria. These results were quantitatively confirmed by a detailed analysis of the vertical offset of the FORC central ridge (Figs. 1L-M), given its sensitivity to the magnetic moment of individual magnetosomes and therefore to their spontaneous magnetization, which in turn depends on the degree of magnetite oxidation (Supporting Notes, Fig. S20). Consequently, we propose that H_2_O_2_ tolerance involved the magnetite nanoparticles. An antioxidant function of magnetosomes protecting AMB-1 from H_2_O_2_ provides a satisfactory explanation for several observations, such as the low Verwey temperature in MTB that is compatible with H_2_O_2_-induced magnetite oxidation (37), or the increased levels of reactive oxygen species in MTB devoid of magnetite measured with fluorescence (10, 13). Yet, an antioxidant role is difficult to reconcile with magnetite, since it is well known to catalyze the conversion of H_2_O_2_ into highly reactive OH• radicals (38), causing non-specific oxidative damage to DNA, proteins and membranes (39, 40). In addition, fluorescence-based quantifications of reactive oxygen species are also compatible with oxidative properties of magnetite in the specific case of MTB (Supporting Notes) (10, 13). Recently, it has been proposed that H_2_O_2_ conversion proceed via a non-radical pathway under the basic pH conditions of magnetosomes (34, 41). The non-radical pathway is enhanced in the presence of phosphate, a key component of the magnetosome membrane in contact with magnetite (42, 43). Our findings support the antioxidant properties of magnetosomes, and we propose a mechanistic model enabling this function (Fig. 5). According to this model, magnetite-catalyzed Fenton reactions reduce intracellular H_2_O_2_ into OH^−^ within magnetosomes. The basic pH of the magnetosome medium (41) increases magnetite stability and also favors the non-radical pathway of Fenton reactions. Sequestration of magnetite into lipid vesicles could further stimulates the non-radical pathway thanks to the phosphate groups of phospholipids comprising the magnetosome membrane. Consequently, growth, ATP, RNA-seq, FORC and XAS/XMCD results obtained from ΔMIS AMB-1 pointed to increased antioxidant properties of fragmented magnetite chains, although this benefit comes at the expense of magnetic properties (Figs. 1J,K). A likely explanation for these findings is that chain fragmentation and the associated increase in chain length, reaching the cell poles (Fig. 1A, Fig. S1), prevent H_2_O_2_ accumulation in these regions. The higher nanoparticle length in wild-type AMB-1 increased by ~20 % the mean reactive surface area per cell for reduction of H_2_O_2_ (Supporting Notes). In addition, chain fragmentation in wild-type bacteria increased the number of nanoparticles having one side facing no other reactive nanoparticle. Both reactive surface area and neighboring particles are important parameters controlling reaction kinetics (44). Therefore, we propose that magnetosome chains represent an evolutionary trade-off between magnetoreception and their antioxidant function.

**Fig 5.**
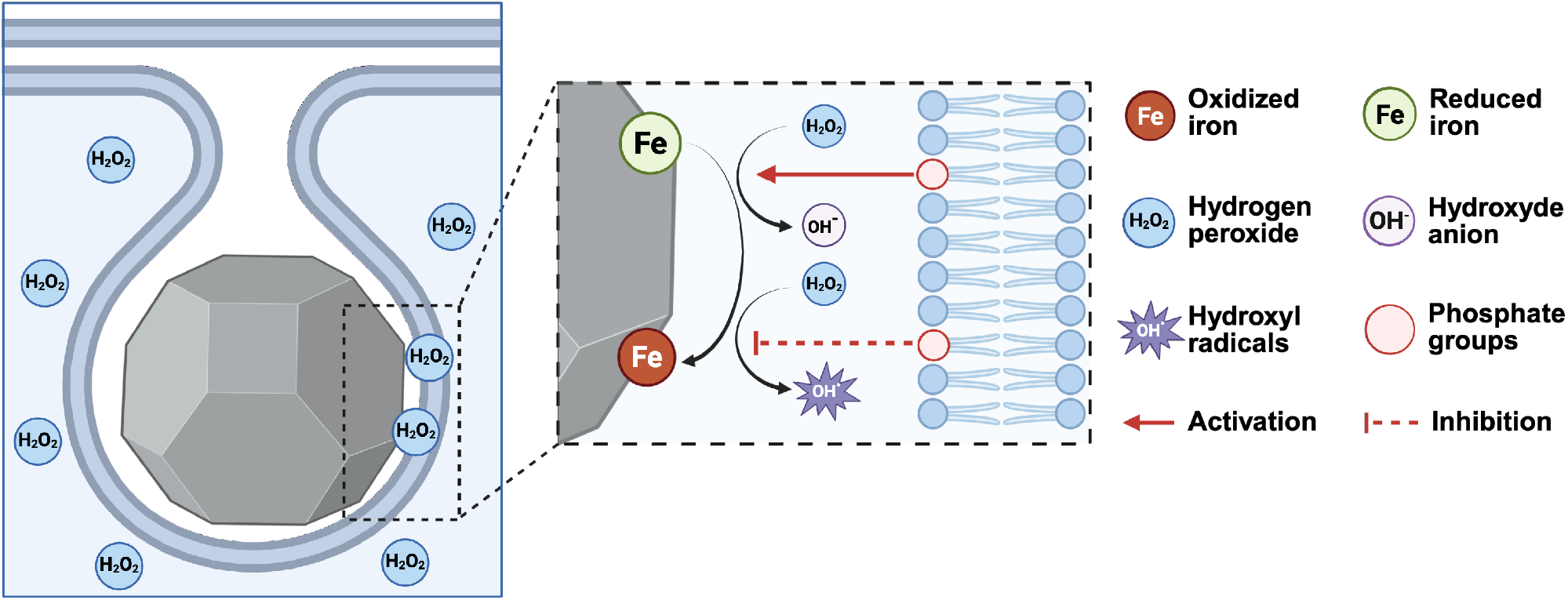
Proposed model for reduction of H_2_O_2_ by magnetite contained in magnetosomes. Magnetosomes are formed from the invagination of the cytoplasmic membrane of bacteria. Phosphate groups of the phospholipid membrane enhance the non-radical pathway of Fenton reactions.

## Discussion

During the early Archean eon (4 – 2.5 Ga), the oceans were anoxic and rich in Fe^2+^, while non-biological sources of H_2_O_2_ were insufficient to generate oxidative stress in microorganisms (15). This changed with the emergence of cyanobacteria performing oxygenic photosynthesis (15). The oxygen they produced was rapidly consumed through oxidation of Fe^2+^, preventing its accumulation in the ocean and atmosphere but secondarily produced H_2_O_2_. As a result, microorganisms became exposed to H_2_O_2_-mediated oxidative stress before the Great Oxidation Event. In this contribution, we demonstrated that magnetosome gene clusters help AMB-1 to overcome high levels of H_2_O_2_. The more oxidized state of nanoparticles in wild-type cells supported an active role of magnetite during this process. According to this proposed scenario, magnetite nanoparticles could have enabled ancestral MTB to survive H_2_O_2_-induced toxicity when cyanobacteria emerged (Fig. 6). Once Fe^2+^ in oceans was consumed by O_2_ reduction, molecular oxygen became sufficiently abundant to establish gradients in natural waters. At this point, assembly of magnetosomes into chains may have enabled bacteria to efficiently swim away from highly oxygenated surface waters using the magnetic field of the Earth. If confirmed, this scenario would imply that Earth’s oxygenation triggered magnetite-based magnetoreception in bacteria (Fig. 6). Consequently, the discovery of magnetite produced by MTB in rocks would point to ancient H_2_O_2_ produced by early oxygenic photosynthesis, which cannot be detected using current geochemical proxies (45, 46). Ultimately, this would reveal the initiation of Earth’s oxygenation caused by the emergence of cyanobacteria and oxygenic photosynthesis. Identification of ancient magnetite produced by MTB in rocks, together with the broader geochemical context, will be needed to characterize the redox conditions of paleoenvironments hosting ancient MTB and test the proposed model for their emergence.

**Fig 6.**
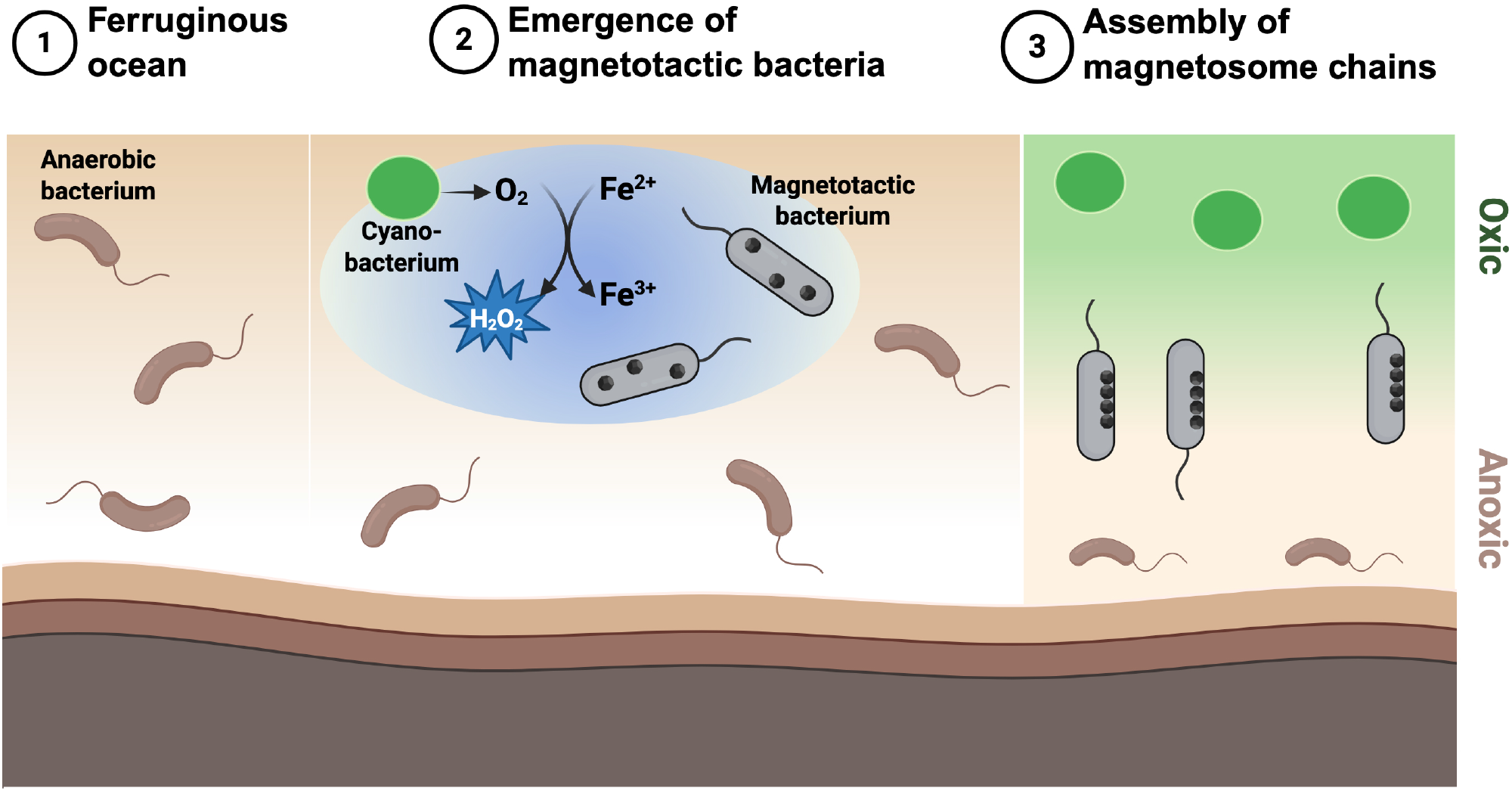
Proposed model for the emergence of magnetotactic bacteria in the Archean ocean. Ferrous iron abundant in the Archean oceans scavenged much of the oxygen produced by early cyanobacteria. This process generated hydrogen peroxide, exposing microorganisms to oxidative stress. Such selective pressure may have driven the emergence of early magnetotactic bacteria, whose magnetosomes helped them cope with these oxidizing conditions. As the oceanic ferrous iron reservoir became depleted, oxygen started to accumulate, establishing redox and oxygen gradients throughout the water column. The subsequent organization of magnetosomes into chains could have allowed magnetotactic bacteria to align with Earth’s magnetic field, enabling efficient navigation along these gradients and facilitating escape from highly oxidative environments.

## Materials and Methods

### AMB-1 strains and growth conditions

*P. magneticum* AMB-1 (ATCC 700264) was cultivated following a protocol previously published (18). The sole iron source provided to AMB-1 cultures was Fe(III)-citrate at 100 µM. ΔMAI and ΔMIS AMB-1 cells were previously produced (16, 18). All AMB-1 strains were recovered from glycerol stocks in 1.45 mL of medium introduced in 1.5-mL Eppendorf tubes. The latter were kept at 28 °C for 4 days. Pre-cultures were then diluted to a 600 nm - Optical Density (OD_600_) of 0.002 to start cultures in (i) Eppendorf tubes containing 1.45 mL of medium or (ii) in 96-well microplates (reference 655180, Greiner Bio-One GmbH, Germany). Cultures conducted in Eppendorf tubes were used to investigate the effect of the magnetic field conditions on bacterial growth and physiology, which were monitored at regular time intervals through OD_600_, flow cytometry and ATP measurements. Cultures under either 2-mT or zero-field conditions were conducted simultaneously in the same incubator at 28 °C to ensure identical temperature conditions (Fig. S6). In the 2-mT field condition, a constant and unidirectional field was applied with Helmholtz coils purchased from 3B scientific. To enable zero-field condition, bacterial cultures were placed in a zero-gauss chamber coated with µ-metal (Soudupin, France). The intensity of the external fields was checked with a GM08 gaussmeter (Hirst Magnetic Instrument), and was 2.060 and 0.016 mT in the coil and the gauss chamber, respectively (i.e., the intensity of the magnetic field of the Earth is about 0.050 mT). Cultures in 96 well-microplates were conducted at 28 °C using an Infinite M200 plate-reader (Tecan, Switzerland) to monitor growth via OD_600_ measurements at short time intervals (30 min). The volume of cultures under these conditions was 350 µL. The microplate reader shakes cultures at each time point. Microplates were sealed with a plastic film (ViewSeal, sealer, Greiner Bio-One GmbH, Germany). Experiments aiming at testing the effect of H_2_O_2_ on growth were performed in the same culture conditions with addition of increasing H_2_O_2_ concentrations obtained via dilution of a 30 % (w/w) commercial solution (Sigma-Aldrich, Merck, Germany) in the growth medium.

### C_**mag**_ measurements

The magnetic behavior of bacteria was assessed by adapting the calculation of the coefficient of magnetism (C_mag_) (47). This coefficient relies on the differential measurement of a culture’s OD_600_ when a magnet is oriented either vertically or horizontally close to the cell suspension. Several calculation methods have been previously used, such as 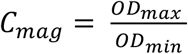, or 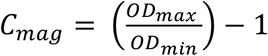. Here, C_mag_ is calculated as 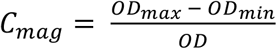, where OD_max_ and OD_min_ are the maximal and minimal optical densities, respectively, and OD is the optical density when no magnet is used. Normalizing to the unperturbed signal rather than to OD_min_ (16) expresses the magnetically induced change in light scattering as a fraction of the baseline, reduces sensitivity to noise at low cell densities, and minimizes the influence of differences in cell concentration between cultures. Raw data of OD, OD_max_ and OD_min_ are reported in Table S8. Cell elongation of ΔMIS and ΔMAI strains were observed during stationary phase only (see main text), meaning that different cell morphology did not impact C_mag_ measurements.

### Transmission electron microscopy

Bacterial samples were deposited on carbon-coated copper-grids (Delta Microscopies, France) adapting a protocol described elsewhere (16). Bacteria deposited on grids were rinsed with Milli-Q water to remove salts precipitated from growth media components. Samples were observed with a FEI Tecnai G2 Biotwin (Institut de Radioprotection et de Sûreté Nucléaire in Cadarache, France) and a FEI Tecnai 12 (University of California, Berkeley, USA) microscopes operating at 100 and 120 kV, respectively.

### X-ray diffraction

XRD measurements were performed at the µSpot beamline (BESSY II synchrotron facility, Herlmotz Zentrum für Materialen und Energie, Berlin, Germany). The procedure for data acquisition and processing is provided in a previously published contribution (48). The scattered intensity was calculated as a function of the momentum transfer Q, defined as 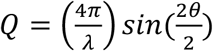 with λ and θ being the photon wavelength and the scattering angle, respectively.

### X-ray absorption spectroscopy (XAS and XMCD)

XAS and XMCD measurements were performed at Soleil synchrotron facility (DEIMOS beamline equipped with a total electron yield detection method (1 meV energy resolution), which probes the first ~5 nm of the sample), at 3 K and under a magnetic field of 6 T. Detailed methodology for measurements and preparation of standard materials is provided in Supporting Information. Bacterial samples were cultivated under standard conditions, centrifuged (8,000 rpm for 10 min), rinsed with Milli-Q water to remove excess water and salts, and immediately dried. All samples were stored under anoxic conditions for transportation to Soleil synchrotron. The DEIMOS beamline possesses a glove box for sample manipulation and mounting on sample holder in an oxygen-free environment. Samples were spread onto carbon tape on copper sample holders, which were then transferred to the superconducting magnet for loading in and acquisition of XAS and XMCD spectra.

### First-order reversal curves

Magnetic properties of bacterial samples were characterized with high-resolution FORC analyses using a vibrating-sample magnetometer (Lakeshore 8600 VSM). Samples were transferred in 1.5 mL-Eppendorf tubes and dried at room temperature in a dessicator. Measurements were performed in steps of 0.2 mT for all samples, using a stepwise approach to the reversal field to avoid overshooting artifacts (49), and a pause of 3 s at reversal. Measurements have been processed with VARIFORC (50) using a variable smoothing approach that grants maximum resolution along *B*_*c*_ = 0, for a correct characterization of the central ridge. Procedure for coercivity analysis is detailed in Supporting Information.

### Off-axis electron holography

Magnetic flux in the magnetosome chains was mapped at the single-crystal level with off-axis electron holography at the Centre d’Elaboration de Matériaux et d’Etudes Structurales (CEMES – Toulouse, France), using a Hitachi HF3300 C microscope operating at 300 kV and following the procedure detailed elsewhere (9).

### Measurements of native magnetic moments

We investigated the native magnetic state of wild-type and ΔMIS AMB-1 strains at the population level by comparing the magnetization of two different preparations of aqueous cell suspensions in a maximum external field of 2 mT, using a vibrating-sample magnetometer (VSM – Lakeshore Micro-Mag 3900). The external field is sufficiently large to partially align the cells, but not large enough to alter their native magnetic configuration. In this case, the magnetization of the suspension is expected to be proportional to the mean strength of the magnetic moments of individual cells. The first preparation contains cells directly taken from the culture and is used to assess the native magnetic moments. In the second preparation, the same suspension is previously exposed to a 200-mT field. This field is strong enough to reset the native magnetic states, replacing them with a saturation remanent state that corresponds to the maximum magnetic moment that can be maintained in a null field. Cell suspensions for these experiments were prepared by centrifugation of AMB-1 cultures (8,000 rpm, 10 min), and subsequent suspension in 10 ml of phosphate buffer (PBS). 100 μl of cell suspension were transferred in plastic vials (diameter of 4 mm) and placed in the VSM for acquisition of magnetization at room temperature.

### Flow cytometry

Bacterial counts were determined by flow cytometry at different time points during growth using a Guava® easyCyte 5HT flow cytometer (Merck Millipore, Germany). Bacterial samples were diluted between 4 to 1200 times – depending on their growth stage – in sterile and filtered growth medium to prevent saturation of the cytometer and ensure that all bacteria can be counted (between 100 and 800 counts/µL). To maximize cell detection, diluted bacterial suspensions were incubated with SYBR® Green II Nucleic Acid Gel Stain (Lonza, Switzerland) for 1 hour (1/10,000 vol:vol dilution of the commercial solution). Filtered medium and unstained bacteria were used to establish the signal background. Bacterial fluorescence induced by SYBR® Green II was recorded on the Green B channel (525/30 nm) after excitation by the Guava® easyCyte 5HT blue laser (488 nm). Analyses were performed using a Green-B (GRN-B) threshold > 10. Bacteria were optically resolved based on their side scatter and Green (GRN-B) fluorescence intensities. Counts obtained in the stained samples were considered to determine bacterial abundances.

### ATP quantification

Adenosine triphosphate (ATP) content of the different strains was quantified using the BacTiter-Glo kit (Promega, UK), which quantifies ATP via a luminescence-based method. Assays were performed following the manufacturer instructions in 96 well-microplates (reference 655094, Greiner Bio-One GmbH, Germany) on non-diluted or 10-times diluted samples - depending on their growth stage. Luminescence was measured after 5 to 10 minutes of incubation at 30 °C using an Infinite M200 plate-reader (Tecan, Switzerland). Preliminary tests were performed to ensure (i) linearity between bacterial concentrations and ATP measurements, and (ii) reproducibility of the assay between the different strains. Excellent linearity between ATP content (i.e., luminescence emission) and OD_600_ measurements was observed (R^2^ = 0.99) for OD_600_ < 0.05 (Fig. S21). Saturation of luminescence signal is observed for higher OD_600_. Therefore, all cultures with OD_600_ higher than 0.05 were diluted prior to ATP quantification to fall within the linearity range. In addition, these calibrations were highly similar between all AMB-1 strains (Fig. S21), showing that the BacTiter-Glo kit is adapted to ATP measurements and give comparable measurements in the bacterial samples. We verified that varying ATP contents cannot be attributed to varying cell volumes, as ATP per OD unit and ATP per cell remained consistent throughout the exponential phase (Fig. S22).

### RNA extraction and sequencing

To assess bacterial metabolic states, *P. magneticum* AMB-1 (wild-type, ΔMIS and ΔMAI) was cultivated under standard conditions (see above) in 50-mL Falcon tubes with (5 µM) or without H_2_O_2_, and harvested during exponential phase (40 h of culture). A total of 10^8^ cells were harvested for each sample. For each condition, three independent biological replicates were used. Total nucleic acids were extracted using the TRIzol Max bacterial RNA isolation kit (TRIzol), followed by removal of genomic DNA using a gDNA elimination column. RNA was further purified using the RNeasy Plus kit (Qiagen). Prior to library preparation, ribosomal RNA (rRNA) was depleted to enrich for messenger RNA (mRNA). Libraries were constructed and sequenced on the Illumina NovaSeq X Plus platform using a paired-end 150 bp strategy (Novogene, UK). Approximately 4 Gb of raw data were generated per sample, yielding an average of 13 to 15 million paired-end reads per library, ensuring sufficient coverage for the transcriptome of *P. magneticum* AMB-1. Raw sequence data processing is detailed in Supporting Information. All data can be accessed on ArrayExpress with accession number E-MTAB-17630.

### Statistics

All statistical analyses were performed with R. Datasets were first analyzed for normality using the Shapiro-Wilk test. For checking the significant differences between conditions, we used ANOVA and Kruskal-Wallis tests for normally and non-normally distributed datasets, respectively. For comparing C_mag_ of wild-type and ΔMIS specifically (Fig. 1H), a Wilcoxon test was used. The coordination between the wild-type response to H_2_O_2_ and the basal expression differences in the mutans was assessed using the Spearman’s rank correlation on variance-stabilizing transformation (VST)-normalized data. Sixteen genes involved in the oxidative stress response were selected: *rpoE, rpoN, rpoD, oxyR*-like, *soxR, gpx, sodA, sodC, ahpC, gor, acrA, acrC, trxA-1, trxA-2, trxB* and *ftsH*. For each mutant, two variables were calculated: x, representing the difference in expression between the wild-type at 5 µM and 0 H_2_O_2_, and y, representing the difference in expression between the mutant at 0 H_2_O_2_ and the wild-type at 0 H_2_O_2_. Spearman’s rank correlation coefficient was calculated for the 16 target genes (ρ_set) and across all genes in the genome (ρ_genome). The statistical significance of the target gene set was assessed using a permutation test. Specifically, 5 000 random gene sets of 16 genes were sampled, and the proportion of sets exhibiting a mean correlation greater than or equal to the observed correlation was used to estimate the *P*-value. *P*-values were adjusted for multiple testing using the Benjamini-Hochberg procedure.

## Acknowledgements

We thank Magali Floriani from *Institut de Radioprotéction et de Sûreté Nucléaire* (IRSN) for assistance with electron microscopy. We are also grateful to Marie Bertrand and Yonghua Li-Beisson (EBMP, BIAM) for their support and help with Flow Cytometry experiments. M.G. and M.A. are supported by a grant through the French National Research Agency (BIOMAG project: ANR-23-CE49-0007) and the CNRS TelluS/INTERRVIE program (grant #MagBio). D.F. is support by the EU through the BioMagnetLink ERA Chair project (grant agreement ID: 101187789). The research leading to these results has received funding from the European Union Horizon 2020 Research and Innovation Program under Grant agreement no. 823717 – ESTEEM3. The authors acknowledge the French National Research Agency under the “Investissement d’Avenir” Program reference no. ANR-10-EQPX-38-01 and the “Conseil Régional Midi-Pyrénées” and the European FEDER for financial support within the CPER program. This work was also supported by the international associated laboratory M2OZART. We acknowledge MPIKG and HZB for the allocation of beamtime under proposal number 212-10564-CR. This work received support from Soleil synchrotron facility under the awarded beamtime proposal #20210218. This work was supported by the Spanish State Research Agency (AEI) / 10.13039/501100011033 and by the Ministry of Science and Innovation under projects MCINN-24-PID2023-150968OA-I00 and CNS2025-166831. Figures 3, 4, 5 and 6 were created with BioRender (https://BioRender.com).

