## Supplementary file for "Magnetic organelle gene clusters enable bacterial growth and ATP homeostasis under oxidizing conditions"

#### This PDF file includes:

Supporting Notes
Supporting Figures
Supporting Tables
Supporting References

### Supporting Notes

#### Coercivity analysis

Coercivity analysis of the central ridge extracted from first-order reversal curves requires a model that is able to reproduce coercivity distributions with finite values at $B_c = 0$. Functions commonly used to fit magnetization curves on a logarithmic field scale, such as modifications of the normal distribution (1), are not well suited for this purpose, since they all converge to zero at $B_c = 0$. Therefore, coercivity analysis has been performed on a linear field scale using a linear combination of two gamma distribution functions modified to reproduce positive distributions with variable location parameter $\mu$, width parameter $\sigma$, and skewness $k$. The function with positive (right) skewness is defined as:

$$g(x,\mu,\sigma,k) = \frac{1}{\Gamma(1/k)\left(\sigma\sqrt{k}\right)^{1/k}}\left(x-\mu+\frac{\sigma}{\sqrt{k}}\right)^{1/k-1}\exp\left[-\frac{x-\mu}{\sigma\sqrt{k}}-\frac{1}{k}\right]$$

for $k > 0$ and $\mu > \sigma/\sqrt{k}$. The counterpart with negative (left) skewness is given by:

$$g(x,\mu,\sigma,k) = \frac{1}{\Gamma(-1/k)\left(\sigma\sqrt{-k}\right)^{1/k}}\left(-x-\mu+\frac{\sigma}{\sqrt{-k}}\right)^{1/k-1}\exp\left[-\frac{x+\mu}{\sigma\sqrt{-k}}+\frac{1}{k}\right]$$

for $k < 0$ and $\mu > -\sigma/\sqrt{-k}$. The limit for $k \to 0$ is a normal distribution with mean $\mu$, and standard deviation $\sigma$. A one-sided exponential function is obtained with $|k| = 1/2$.
The central ridge coercivity distributions $f^{\mathrm{wt}}$ of wild-type cells and $f^{\Delta\mathrm{MIS}}$ of ΔMIS cells have been fitted by least-squared minimization of the joint model residuals:

$$m_{\mathrm{LC}}^{\mathrm{wt}} g(B,\mu_{\mathrm{LC}},\sigma_{\mathrm{LC}},k_{\mathrm{LC}}) + m_{\mathrm{HC}}^{\mathrm{wt}} g(B,\mu_{\mathrm{HC}}^{\mathrm{wt}},\sigma_{\mathrm{HC}}^{\mathrm{wt}},k_{\mathrm{HC}}^{\mathrm{wt}}) - f^{\mathrm{wt}}(B)$$

And:

$$m_{\mathrm{LC}}^{\Delta\mathrm{MIS}} g(B,\mu_{\mathrm{LC}},\sigma_{\mathrm{LC}},k_{\mathrm{LC}}) + m_{\mathrm{HC}}^{\Delta\mathrm{MIS}} g(B,\mu_{\mathrm{HC}}^{\mathrm{wt}},\sigma_{\mathrm{HC}}^{\mathrm{wt}},k_{\mathrm{HC}}^{\mathrm{wt}}) - f^{\Delta\mathrm{MIS}}(B)$$

with $m$-parameters indicating the magnetic contribution of the corresponding component and the subscripts LC and HC for the low-coercivity and high-coercivity components, respectively. Distinct shape parameters have been used to model the HC components, in order to accommodate the evident differences between wild-type and ΔMIS populations, while a single set of shape parameters has been used for the LC component, since this was sufficient to reproduce the coercivity distributions within their respective error ranges. This avoids model instabilities due to overfitting. Model uncertainties have been estimated with a Monte Carlo approach by fitting 100 modified versions of the original coercivity distributions where a random gaussian error with same standard deviation as the error estimate of the central ridge has been added.

#### Interpretation of magnetic measurements

##### *FORC diagrams*

FORC diagrams of wild-type and ΔMIS feature common signatures typical for intact single chains of magnetosomes (2). These signatures include (i) a sharp ridge along $B_u = 0$, called central ridge, and (ii) a doublet of negative and positive amplitudes extending over $B_u < 0$ and almost symmetric about the $B_u = -B_c$ diagonal. The central ridge is produced by switching between two-state magnetic configurations where the magnetic moments of all magnetosomes are consistently parallel or antiparallel to the chain axis (3), while the positive/negative doublet is produced by reversible magnetic moment rotation in the applied field.

FORC diagrams also feature details signaling important departures from the behavior of idealized, gap-free magnetosome chains. For instance, positive FORC contributions over $B_u > 0$, as denoted by the lowermost contour lines, reflect magnetic switching processes occurring in an internal biasing field produced by nearby non-switching magnetic sources. Most importantly, the distribution of coercivities includes non-negligible contributions at and close to $B_c = 0$, especially in the case of wild-type cells (Fig. S4). For comparison, mature, gap-free chains require a finite minimum field $B_c^{min}$ to be switched, owing to the strong magnetic anisotropy imposed by the chain axis and the strong magnetic coupling between the crystals. The corresponding coercivity distribution is zero for all fields $B_c < B_c^{min}$, as seen with numerical FORC simulations (3) and direct measurements (4, 5). The central ridges depicted in Fig. S4, on the other hand, are well explained by two coercivity components, one (HC) starting at $B_c^{min} \approx 10$ mT and extending to ~70 mT, which fulfills the expected properties of ideal chains, and the other (LC) covering the lower coercivity range comprised between 0 and 40 mT. The difference between the median fields of component HC (~35 mT for wild-type cells vs. ~29 mT for ΔMIS AMB-1, Table S2) can be explained by the lower particle elongation in the mutant strain (6) (Fig. S4, Table S1).

##### *Thermal stability considerations*

The vanishing coercivities implied by the LC component have two possible explanations. The first explanation relies on magnetic viscosity, which is the spontaneous magnetization decay of magnetic contributions close to the lower thermal stability threshold. This threshold is given by

$$\beta = \frac{V \mu_s B_a}{2 k_B T} = \ln \frac{t}{\tau_0} \quad (116)$$

where $\mu_s$ is the spontaneous magnetization of the magnetic particles under consideration (e.g., 480 kA/m for magnetite) and $V$ their volume, $B_a \approx 2B_c$ is the so-called anisotropy field, $k_B$ the Boltzmann constant, $T$ the absolute temperature, $t$ the measurement time, and $\tau_0 \approx 0.1$–1 ns the atomic reorganization time. For isolated, nearly equidimensional magnetite particles, the resulting threshold diameter is ~26 nm. The added anisotropy of magnetosome chains reduces this threshold to <15 nm for gap fractions <0.5 (7). Almost all crystal sizes of the wild-type and ΔMIS populations exceed this threshold, meaning that the magnetization of chains and chain fragments with three or more magnetosomes is thermally stable.

Nevertheless, thermal energy can assist magnetic moment reversals, effectively decreasing the thermal-free switching field $B_{sw}$ by an amount $B_f$ called fluctuation field (8), so that $B_c = B_{sw} - B_f$ with

$$\frac{B_\mathrm{f}}{B_\mathrm{sw}} = \frac{1}{\beta} = \frac{2k_\mathrm{B}T}{V\mu_\mathrm{s}B_\mathrm{a}}$$

This equation is valid for any magnetic system if $V$ is interpreted as the effective volume that triggers the switching process, such as for instance the end magnetosomes in a chain (3). The fluctuation field can be understood as a random field that adds to the externally applied field during any type of magnetic measurement, including FORC. In addition to reducing $B_c$ in coercivity distributions, the fluctuation field creates a vertical offset of the central ridge (9) to a mean $B_u$ given by $B_f$ itself. Accordingly, the central ridge offset (Fig. S20) can be used as a direct measure of the fluctuation fields that assist the magnetic switching of magnetosome chains. Since $B_a = 2B_{sw}$ for uniaxial systems of magnetic particles, the fluctuation field equation further simplifies to

$$B_\mathrm{f} = \frac{k_\mathrm{B}T}{V\mu_\mathrm{s}}$$

As seen in Supplementary Fig. S20, $B_f$ never exceeds ~0.37 mT over the entire range of coercivities, which means that switching processes, including those responsible for the LC component, are not governed by magnetic viscosity. This is also confirmed by the ratio $M_{rs}/M_s$ between saturation remanence and saturation magnetization (0.471 for wild-type cells and 0.475 for ΔMIS), which is very close to the theoretical value of 0.5 for randomly oriented, stable chains.

The vertical central ridge offset $\langle B_u \rangle$ for $B_c$ values above the coercivity range of the LC component (i.e., >40 mT) is expected to match exactly the equation for the fluctuation field, since biases that might be introduced by the interaction between chain fragments with opposed polarity (see below) can be excluded. In this case, the effective volume that triggers a magnetic switching of the whole magnetosome chain is given by

$$V\mu_\mathrm{s} = \frac{k_\mathrm{B}T}{\langle B_\mathrm{u} \rangle}$$

Assuming stoichiometric magnetite composition, (i.e., $\mu_s$ = 480 kA/m) for measurements performed at room temperature ($T$ = 293 K), and using $\langle B_u \rangle$ = 0.29 mT for ΔMIS cells at $B_c$ = 40 mT, which is the field closest to the peak of the HC component that does not overlap with the LC range (Fig. S20), one obtains $V$ = 30.7 nm$^3$. This is the volume of a sphere with a diameter of 38.1 nm, close to the real average size of 40 nm for ΔMIS magnetosomes. This result confirms previous findings from numerical simulations (3) that the magnetic moment switching of magnetosome chains is triggered by the end magnetosomes.

*Magnetic model of fragmented chains*

According to the above thermal stability analysis, the LC component must originate from magnetosome chains that cannot be represented by the standard two-state model illustrated above. The much larger relative contribution of LC in wild-type cells (~44%), compared to ΔMIS AMB-1 (~24%), suggests that the origin of this component is related to the presence of chain gaps. Chain gaps decrease the magnetic coupling within the chain. This coupling is responsible for the simultaneous switching of all magnetosomes once a sufficiently large field is applied in the opposite direction (3, 4). Sufficiently large gaps can decouple chain fragments, adding stable magnetic states with mixed fragment polarity, which can exist naturally, as seen by the increase of the cell's total magnetic moment after applying a 200-mT field, or be induced during FORC measurements, as seen by the LC component. The low coercivity component can be explained by the fact that short fragments have a lower anisotropy energy and can be switched by smaller fields. Furthermore, mixed fragment polarity creates opposing stray fields that increase the total magnetic energy, drastically reducing the magnitude of the external field required to switch the system back to a homogeneous polarity with lower energy.

The whole FORC signature of fragmented chains can be understood using the conceptual model of Supplementary Fig. S23. This model considers a chain consisting of two fragments of different lengths separated by a gap. The shorter fragment consists of few magnetosomes (typically 1–4) and is significantly less magnetically stable than the longer fragment. Furthermore, it is assumed that the gap is sufficiently large to reduce the magnetic coupling of the two fragments beyond the point where the chain has only a single pair of magnetic states where all moments are parallel (+S) or antiparallel to (−S) to the chain axis. The weaker coupling enables an additional pair of states (±M) where the two fragments have opposed polarities.

In order to understand the consequence of these additional magnetic states on the FORC signature of the fragmented chain model we start from the definition of the FORC function (9)

$$\rho(B, B_\mathrm{r}) = -\frac{1}{2}\frac{\partial^2 M}{\partial B_\mathrm{r} \partial B}$$

as the mixed derivative of the magnetization curve $M$ with respect to the reversal field $B_\mathrm{r}$ and the applied field $B$. Each magnetization curve starts after resetting the system to positive saturation through the application of a large positive field, followed by a field sweep to the reversal field $B_r$. From this point onwards, the magnetization is measured while the applied field $B$ increases from $B_r$ back to positive saturation. These measurements are repeated for a series of decreasing reversal fields, until the last curve that starts from negative saturation is measured. If this measurement protocol is applied to the fragmented chain model of Fig. S23, we obtain three groups of distinct curves. As long as $B_r$ is larger than the field $-B_1$ required for reversing the short fragment, the chain remains in the +S state yielding identical curves, so that the derivative with respect to $B_r$ is zero and so the FORC function.

At $B_r = -B_1$, the initial state of the corresponding curve, labelled as FORC 1, becomes +M: the long fragment maintains its original positive polarity while the short fragment was switched to negative polarity. The net magnetic moment is still positive, so that this state is denoted with a positive sign. This mixed state is preserved until the applied field reaches the small positive field $B_3$ required to switch the short fragment back to positive saturation. The corresponding +M → +S transition produces a magnetization jump that was not present in the previous curves. Accordingly, $\partial M/\partial B_r$ is a step function of $B$ centered at $B_3$, whose derivative with respect to $B$ yields a positive impulse at $B_3$ (red dot in Fig. S23) with corresponding FORC coordinates $B_c = (B - B_r)/2 = (B_3 + B_1)/2$ and $B_u = (B + B_r)/2 = (B_3 - B_1)/2$. Notice that $B_u$ depends on the difference between the nucleation ($B_1$) and denucleation ($B_3$) fields of the M state, which in turn depends on the strength of the magnetic coupling between the two fragments.

From this point on, subsequent curves are identical to FORC 1 and do not contribute to the FORC diagram (because of the zero derivative with respect to $B_r$), until $B_r$ becomes equal to the field $-B_2$ required to switch the second fragment and bring the chain into the negative saturation state −S. The corresponding curve, labelled as FORC 2, now contains two magnetization jumps corresponding to the successive switching of the shorter fragment and the longer fragment, so that the chain is brought back to positive saturation via the −S → −M and −M → +S transitions at the fields $B_1$ and $B_2$, respectively. The difference between this curve and the previous one contains three magnetization jumps, a negative jump at $B_3$ inherited from the previous curve, and the two positive jumps at $B_1$ and $B_2$, respectively.

The position of the four peaks in the FORC space depends on the relative stability of the two chain fragments and their coupling strength. Decreasing the coupling strength will cause $B_3$ to converge to $B_1$, making the corresponding leftmost peak approach $B_u = 0$ and thus the central ridge. This explains the LC component of the central ridge in terms of contributions from short fragments that are completely or almost completely decoupled from the rest of the chain by sufficiently large gaps.

Magnetic coupling between fragments can be expressed by the stray field $B_i$ of one chain fragment at the center position of the nearest magnetosome of the adjacent fragment. Defining $\gamma > 1$ as the center-to-center distance of consecutive magnetosomes, normalized by their diameter, one has

$$B_{\mathrm{i}} \approx \frac{\mu_{\mathrm{s}}}{12\gamma^3}\sum_{i=1}^{n}\frac{1}{(n_{\mathrm{gap}}+i)^3}$$

where $\mu_s$ is the spontaneous magnetization of the magnetosomes (e.g., 480 kA/m for magnetite), $n$ is the number of magnetosomes in a chain fragment, and $n_{gap}$ is the number of missing magnetosomes constituting the gap. Taking the geometry of the chain in Fig. S1a as an example with $\gamma = 1.3$–$1.5$ and $n_{gap} = 5$–$14$, one obtains $B_i = 0.015$–$0.15$ mT. Since $B_i$ acts as a biasing field for the switching of individual fragments, it coincides with the vertical coordinate $B_u = (B_3 - B_1)/2$ of the LC peak in Fig. S23. Given the small value of $B_i$, it is interpretable as the downward vertical offset of the central ridge over the $B_c$ range of the LC component. This offset adds to the opposite offset created by the fluctuation field, explaining the observed difference between the LC and HC ranges, which is of the order of 0.1 mT.

Smaller gaps, on the other hand, lead to significant FORC contributions with a net positive sign below the central ridge, as seen in Fig. S23. These contributions add to the signature of reversible magnetic moment rotation (2) over $B_u < 0$, breaking its symmetry. The integral of the FORC function over $B_u < 0$ thus yields the magnetization associated with the nucleation and denucleation of mixed states that is not captured with the central ridge. These contributions are much smaller than the LC component of the central ridge and nearly equal for wild-type and ΔMIS populations (Table S2). The difference between the chain geometry of the two populations is thus limited to the more frequent occurrence of large gaps, similar to those depicted in Fig. S1, in the wild-type population.

*Magnetic signature of partially oxidized magnetosome chains*

The equations for thermal stability analysis can be used to verify the mean oxidation state of magnetosomes in wild-type and ΔMIS cells. As discussed above, the activation volume $V$ can be assumed to be proportional to the mean magnetosome volume, so that the ratio between the spontaneous magnetization of wild-type and ΔMIS magnetosomes is given by

$$\frac{\mu_{\mathrm{s}}^{\mathrm{wt}}}{\mu_{\mathrm{s}}^{\Delta\mathrm{MIS}}} = \frac{\langle B_{\mathrm{u}}\rangle^{\Delta\mathrm{MIS}}}{\langle B_{\mathrm{u}}\rangle^{\mathrm{wt}}}$$

Using the $\langle B_u \rangle$ values at $B_c = 40$ mT, closest to the peak of the HC components but still within the coercivity range unaffected by the LC component, the above equation yields a ratio of $0.82 \pm 0.02$. This result can be compared with the spontaneous magnetization of partially oxidized magnetite with equivalent composition obtained from X-ray absorption spectroscopy. The saturation moment for cation-deficient magnetite with structural formula

$$\mathrm{Fe}^{3+}_{2+2z/3}\mathrm{Fe}^{2+}_{1-z}\square_{z/3}\mathrm{O}^{2-}_4$$

decreases linearly(9) from $4\mu_B$ for pure magnetite endmember ($z = 0$) to $3\mu_B$ for pure maghemite endmember ($z = 1$). Accordingly

$$\mu_s = \frac{p_0 + \frac{3}{4}p_1}{p_0 + p_1}\mu_s^0$$

with $p_0$ and $p_1$ being the magnetite and maghemite endmember fractions, and $\mu_s^0 = 480$ kA/m the spontaneous magnetization of magnetite. Ferrihydrite is not magnetic at room temperature and does not contribute to $\mu_s$. Using the values obtained from X-ray absorption spectroscopy, $\mu_s = 405$ kA/m for wild type magnetosomes and 480 kA/m for ΔMIS magnetosomes. The corresponding ratio of 0.84 agrees with the same ratio calculated from the central ridge offsets within the uncertainty of magnetic measurements.

**X-ray absorption spectroscopy (XAS and XMCD)**

We previously identified magnetite, maghemite and ferrihydrite in wild-type bacteria using the same methodology (10). Ferrihydrite reference material was precipitated following published methods (11). Maghemite was obtained from oxidation of chemical magnetite by heating at 120 °C in an oven for 48 h (12). XAS and XMCD spectra of references are given in Fig. S17. Data were obtained as previously described (13) by flipping both the circular polarization (left or right) and the orientation of the magnetic field being either parallel or antiparallel to the photon propagation vector. XAS data were obtained by averaging two spectra corresponding to the left and right polarizations respectively, while XMCD spectra were calculated as the difference between the two when the magnetic field is applied in the direction of the incoming beam (perpendicular to the sample). XAS linear combinations were calculated with the Athena software using the Ifeffit package (Fig. S18). The merit of goodness-to-fit (R-factor) was used to determine the best fits with lowest number of components from magnetite, maghemite and ferrihydrite (i.e., an additional component is only added to the fit if the goodness-to-fit value improves more than 10%). No fitting with maghemite could be acquired for spectra obtained from ΔMIS bacteria, even when adapting the energy range and normalization approach, which consistently yielded a maghemite contribution of zero. Linear-combination analysis of the XMCD spectra using theoretical peak contribution was performed following previously published methodology (14, 15).

**RNA-seq data processing**

Raw sequence data were processed using an RNA-SEQ workflow on Nextflow (16, 17). Reads were aligned to the *P. magneticum* AMB-1 reference genome (NCBI: GCF_000009985.1) using STAR (18). Gene-level quantification was performed with Salmon, utilizing the NCBI RefSeq annotation GCF_000009985.1-RS_2024_12_09 (19). Statistical analyses were conducted using R software (v.4.x) using the DESeq2 package (20). A generalized linear model (GLM) was implemented with the design formula ~ strain + conc + strain:conc to assess: (i) baseline differences between strains, (ii) the global effect of $H_2O_2$ concentration (0 vs 5 μM), and (iii) the interaction between genotype and oxidative stress. Prior to analysis, a filtering step was applied, removing genes with fewer than 10 reads across at least 3 samples. For the global differentially expressed genes (DEG), a two-step filtering approach was employed (Fig. S13). Genes were first selected based on statistical significance with an adjusted p-value (padj) < 0.05 (Benjamini-Hochberg correction), and subsequently filtered for a minimum two-fold change in expression ($|\log_2 FC| > 1$). Conversely, for established pathways of interest (*e.g.*, oxidative stress defense, sigma factors, or respiration), a targeted analysis was performed. For these specific genes, the $\log_2$FC values are reported regardless of the fold-change amplitude, accompanied by their respective adjusted p-values to provide a comprehensive view of their transcriptional modulation. Functional characterization and visualization of these genomic data were supported by the enrichplot (v1.30.5) (21) and clusterProfiler suites (v4.18.4) (22). Exploratory data analysis was performed using Principal Component Analysis (PCA) to evaluate sample clustering and reproducibility. Pathway-specific heatmaps were generated using the ComplexHeatmap package (v2.26.1) (23), with graphical parameters and color scales handled via the circlize framework (v0.4.18) (24). In these visualizations, color intensity represents the $\log_2$(FC), while the specific adjusted *p*-value is explicitly printed within each cell to facilitate direct statistical interpretation.

**Identification of *oxyR*-like in AMB-1**

The amino acid sequence of the OxyR protein from *Escherichia coli* (strain K12, UniProt ID: P0ACQ4) was compared to homologues (OxyR-like) from *Magnetospirillum gryphiswaldense* (WP_106002419.1) (30) and *Paramagnetospirillum magneticum* AMB-1 (AMB_RS01265, NCBI: GCF_000009985.1). Multiple sequence alignment was performed using the T-Coffee algorithm with default parameters (31). The resulting alignment was visualized and processed in Jalview (32). Protein conservation was highlighted using a percentage identity color scheme with a 50% threshold, allowing for the identification of structural and functional residues stabilized across these species (Fig. S24). The alignment reveals a high degree of similarity between the AMB_RS01265 protein from *P. magneticum* AMB-1 and the OxyR-like protein of the closely related magnetotactic strain *M. gryphiswaldense* MSR-1, both of which were initially annotated as LysR-type transcriptional regulators. Furthermore, the conservation of key functional domains in these two sequences relative to the well-characterized OxyR protein from *E. coli* confirms their identity as members of the OxyR regulatory family. Due to its high sequence homology with these known regulators, the AMB_RS01265 protein will hereafter be referred to as OxyR-like throughout this study.

**Detection of reactive oxygen species (ROS) using fluorescent probes**

Fluorescent probes have been widely used in biological systems as indicators of ROS. One of the most commonly used probes is 2',7'-dichlorofluorescin diacetate (DCFH-DA), which is oxidized to the fluorescent compound 2',7'-dichlorofluorescein (DCF) via complex radical reactions. DCFH-DA can be internalized in cells, and is often described as reacting with several reactive oxygen species, including hydrogen peroxide ($H_2O_2$), hydroxyl radicals ($OH^{\bullet}$), and superoxide anions ($O_2^{\bullet-}$). The specificity of DCFH-DA toward individual reactive oxygen species has been extensively investigated, and has led to contradictory conclusions (see for instance LeBel et al., Myhre et al., and references therein) (25, 26). Hydrogen peroxide and hydroxyl radicals alone do not efficiently oxidize DCFH, whereas the presence of iron enhances DCF formation. This observation suggests that iron-mediated Fenton reactions are efficiently detected using this probe rather than individual reactive oxygen species. In addition, DCFH-DA is also very sensitive to reactive nitrogen species, especially peroxynitrite ($ONOO^-$).

Such fluorescent probes have been used in two mutant strains of the magnetotactic bacterium *Magnetospirillum gryphiswaldense* MSR-1 unable to produce either magnetite or the entire magnetosome organelle (27, 28). The fluorescence intensity was higher in both mutant strains, relative to wild-type producing both magnetite and magnetosomes. This has led to the hypothesis of increased intracellular levels of reactive oxygen species in bacteria unable to form magnetite and magnetosomes. However, magnetite is known to catalyze Fenton reactions and the production of ROS (mainly $OH^{\bullet}$):

$$Fe^{2+} + H_2O_2 \rightarrow Fe^{3+} + OH^- + OH^{\bullet}$$

In the case of MTB and oxidation of magnetite by $H_2O_2$, intracellular radical production described would be localized (i.e., in the magnetosomes). A decrease of fluorescence intensity in wild-type bacteria producing magnetosomes is also compatible with an increased oxidative stress in the case of magnetite-catalyzed Fenton reactions. This is due to the much shorter half-life of hydroxyl radicals ($10^{-9}$ s) than of hydrogen peroxide ($> 10^{-3}$ s) (29). Unlike $H_2O_2$, the high and non-specific reactivity of $OH^{\bullet}$ radicals leads to their almost immediate consumption and severely limits diffusion away from magnetosomes (diffusion distance in the cell of > 1 μm and 1 nm for $H_2O_2$ and $OH^{\bullet}$, respectively) (29), thereby preventing their efficient detection by the fluorescent probes in the cytoplasm. $OH^{\bullet}$ produced in magnetosomes is virtually invisible to fluorescent probes, which could lead to a decrease in ROS-associated fluorescence despite increased oxidative stress. Consequently, the previously reported decrease of fluorescence in magnetosome-forming bacteria is compatible with oxidative properties of magnetite.

**Evaluating the reactive surface area of magnetite in wild-type and ΔMIS AMB-1**

To evaluate the effect of impact of nanoparticle elongation on the reactive surface area, two nanoparticle geometries were approximated as prolate spheroids. The surface area can be calculated following:

$$A = 2\pi r^2 \times \left(1 + \frac{R}{re} \arcsin(e)\right)$$

Where r is the minor semi-axis (calculated from the nanoparticle width), R is the major semi-axis (calculated from the nanoparticle length) and e the eccentricity defined as:

$$e = \sqrt{1 - \frac{r^2}{R^2}}$$

The reactive surface area (A) was calculated using Supporting Table 1, yielding a surface area of 18 991 and 17 387 $nm^2$ for wild-type and ΔMIS AMB-1, respectively. The total reactive surface per cell ($A_{cell}$) can be calculated from the mean number of nanoparticles per cell (n) in both strains:

$$A_{cell} = A \times n$$

From data in Supporting Table 1, the total increase of reactive surface area per cell in wild-type bacteria relative to ΔMIS is ~20 %.

#### Supporting Figures

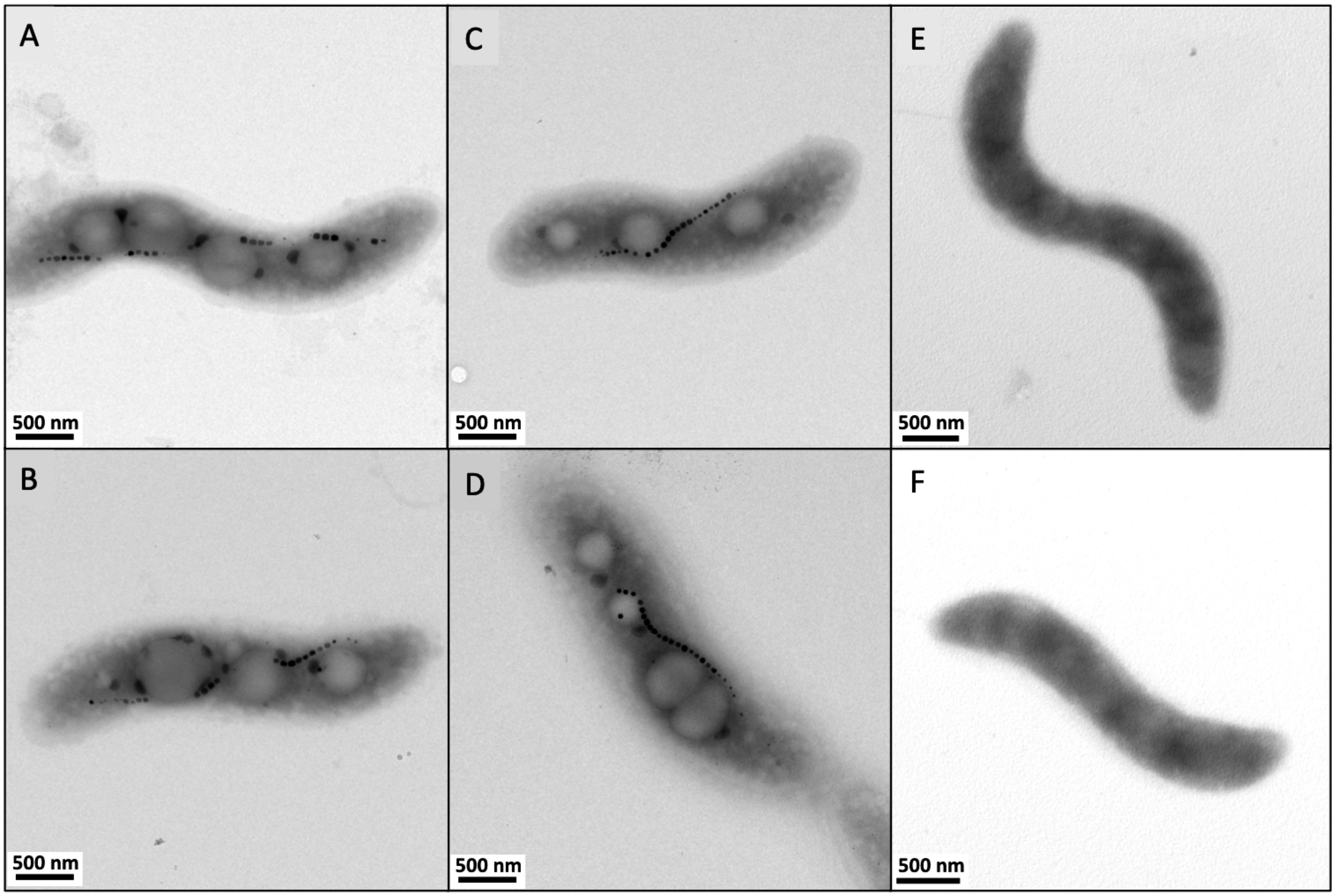


**Fig. S1.** Additional transmission electron microscopy micrographs of (A, B) wild-type, (C, D) ΔMIS and (E, F) ΔMAI AMB-1 bacteria.

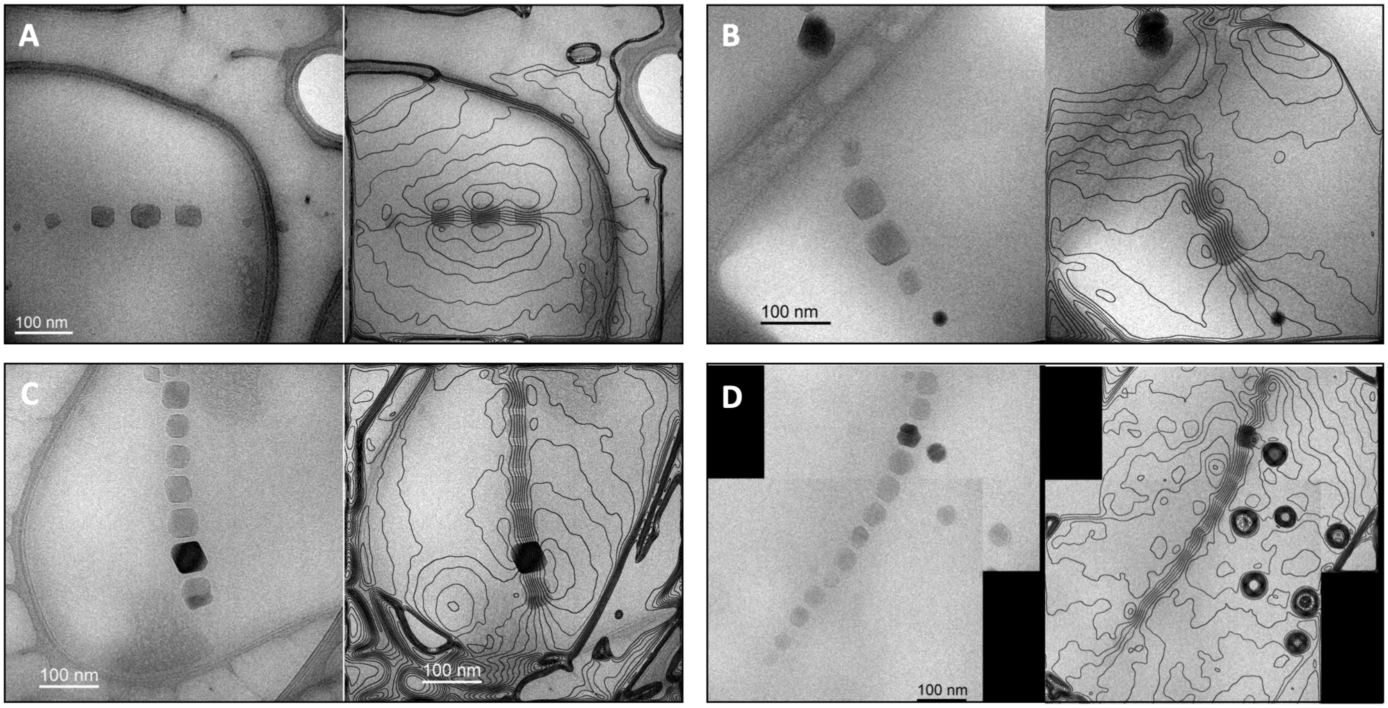


**Fig. S2.** Additional magnetic phase contours determined by off-axis electron holography images of magnetosome chains in (A, B) wild-type and (C, D) ΔMIS AMB-1.

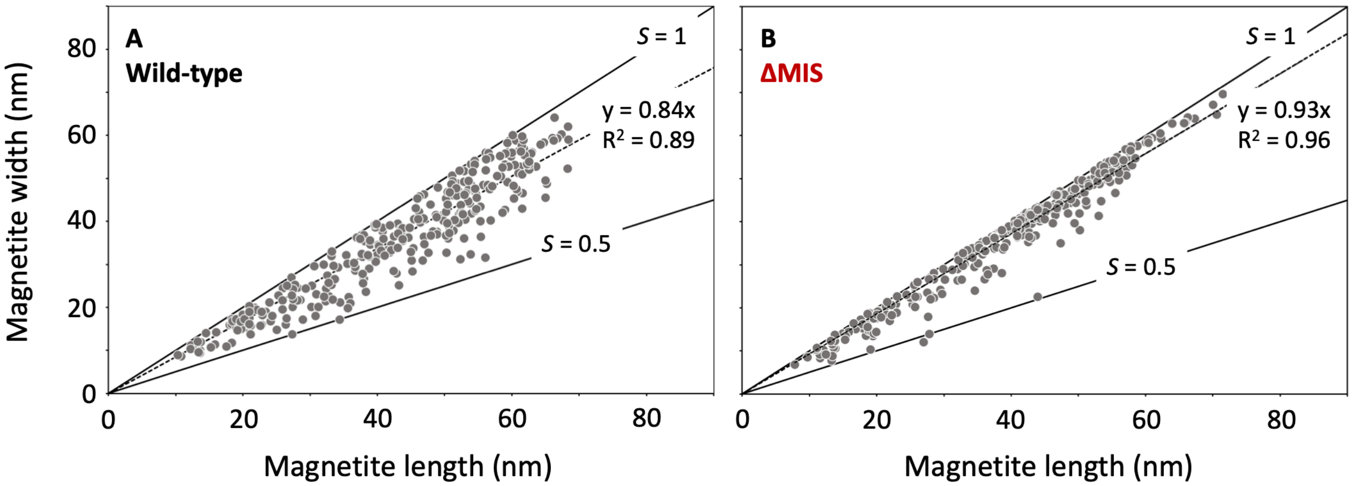


**Fig. S3.** Magnetite particle length, width and shape factor (*S*, defined as the ratio width/length and quantifying the particle elongation) in (A) wild-type and (B) ΔMIS AMB-1 measured from transmission electron microscopy.

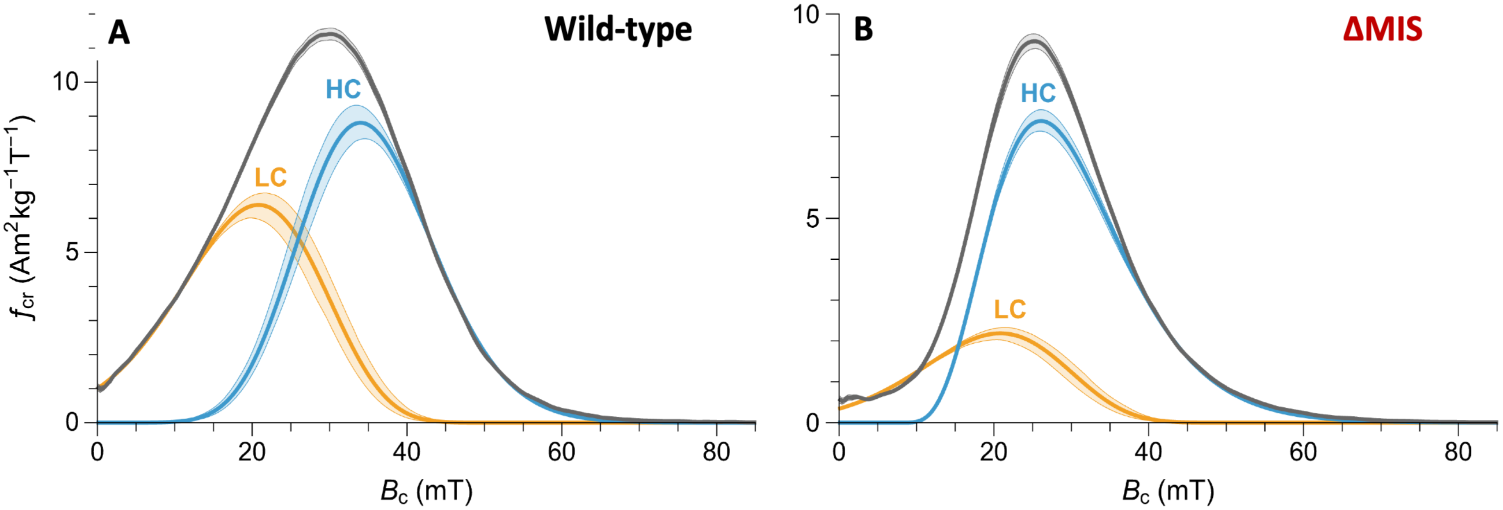


**Fig. S4.** Coercivity analysis of the central ridge of (A) wild-type and (B) ΔMIS AMB-1 populations. The central ridge has been fitted with two modified gamma distributions (LC and HC), with a common LC shape for both populations. Shading around lines represents the 1-σ uncertainty estimate obtained with a Monte Carlo simulation of central ridges with added errors according to the FORC processing output.

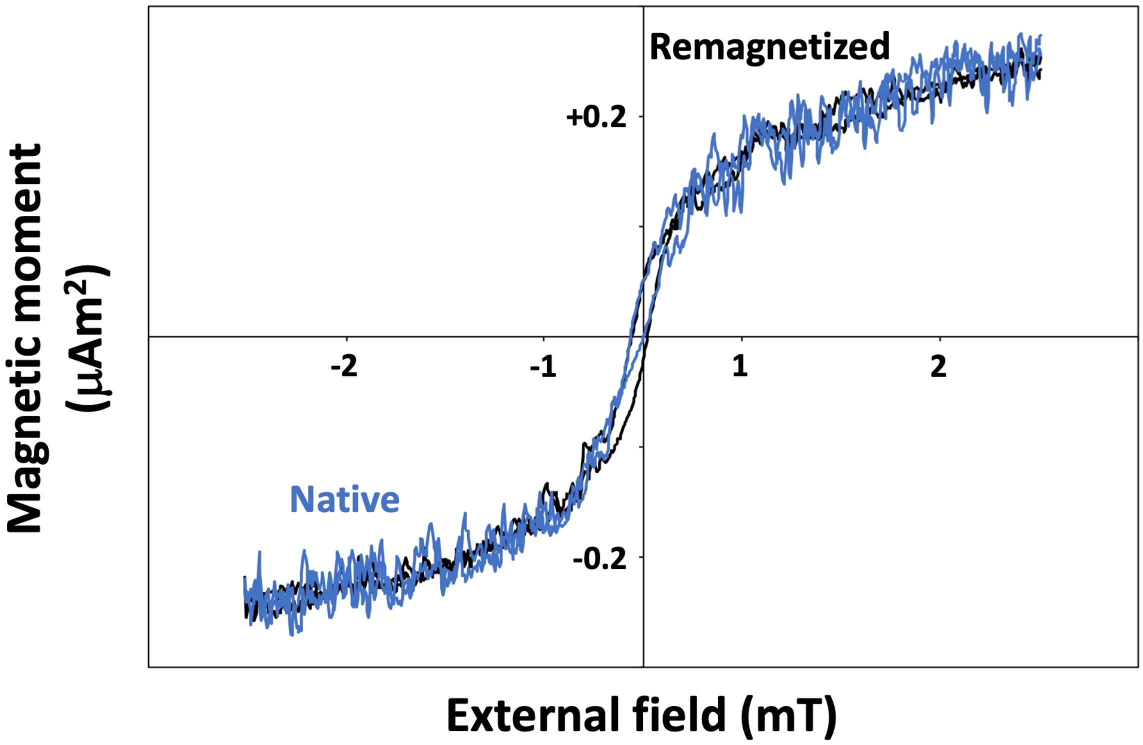


**Fig. S5.** Additional magnetic moment of ΔMIS cultures in their native states (blue curve) and after remagnetization aligning magnetic moment of individual magnetite crystals by exposing the same culture to an external field of 200 mT (i.e., above magnetite coercivity – black curve).

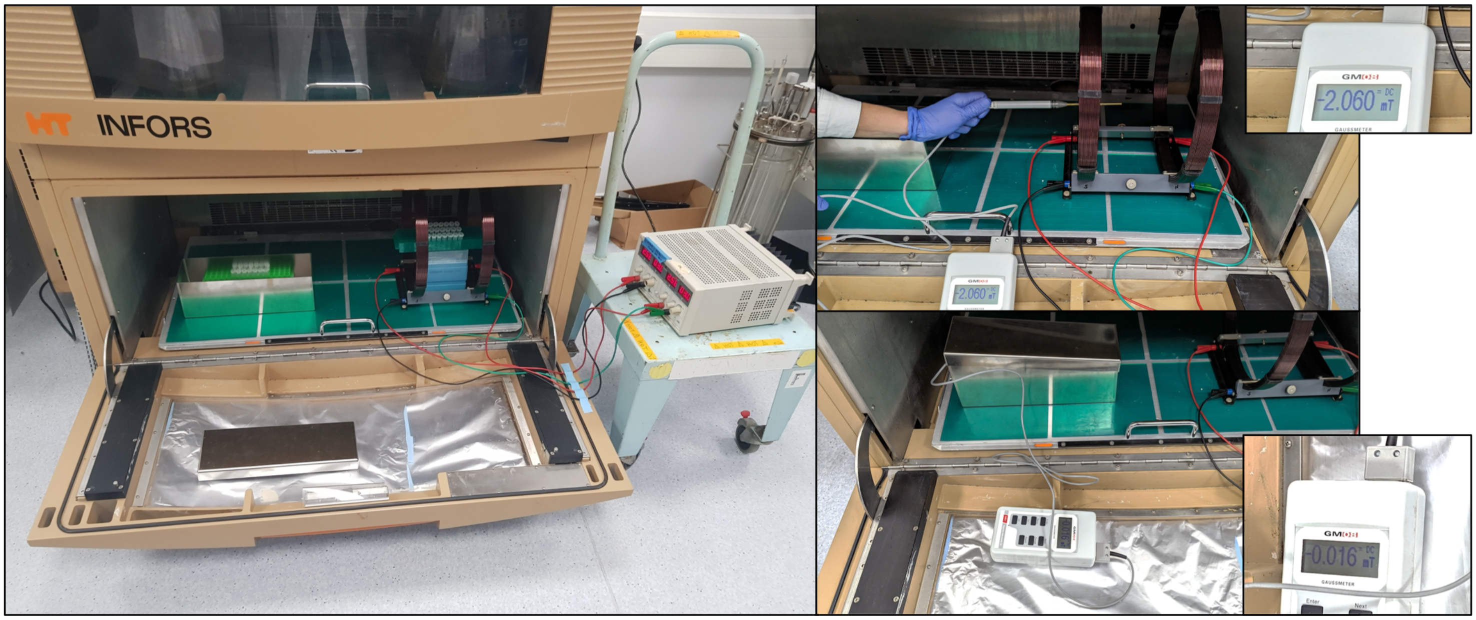


**Fig. S6.** Experimental setup of AMB-1 cultures under controlled magnetic field conditions showing the μ-metal box and Helmholtz coils (left panel) placed into the incubator. The intensity of the external field in the metal box and the coils was checked with a gaussmeter (right panels) before starting bacterial cultures.

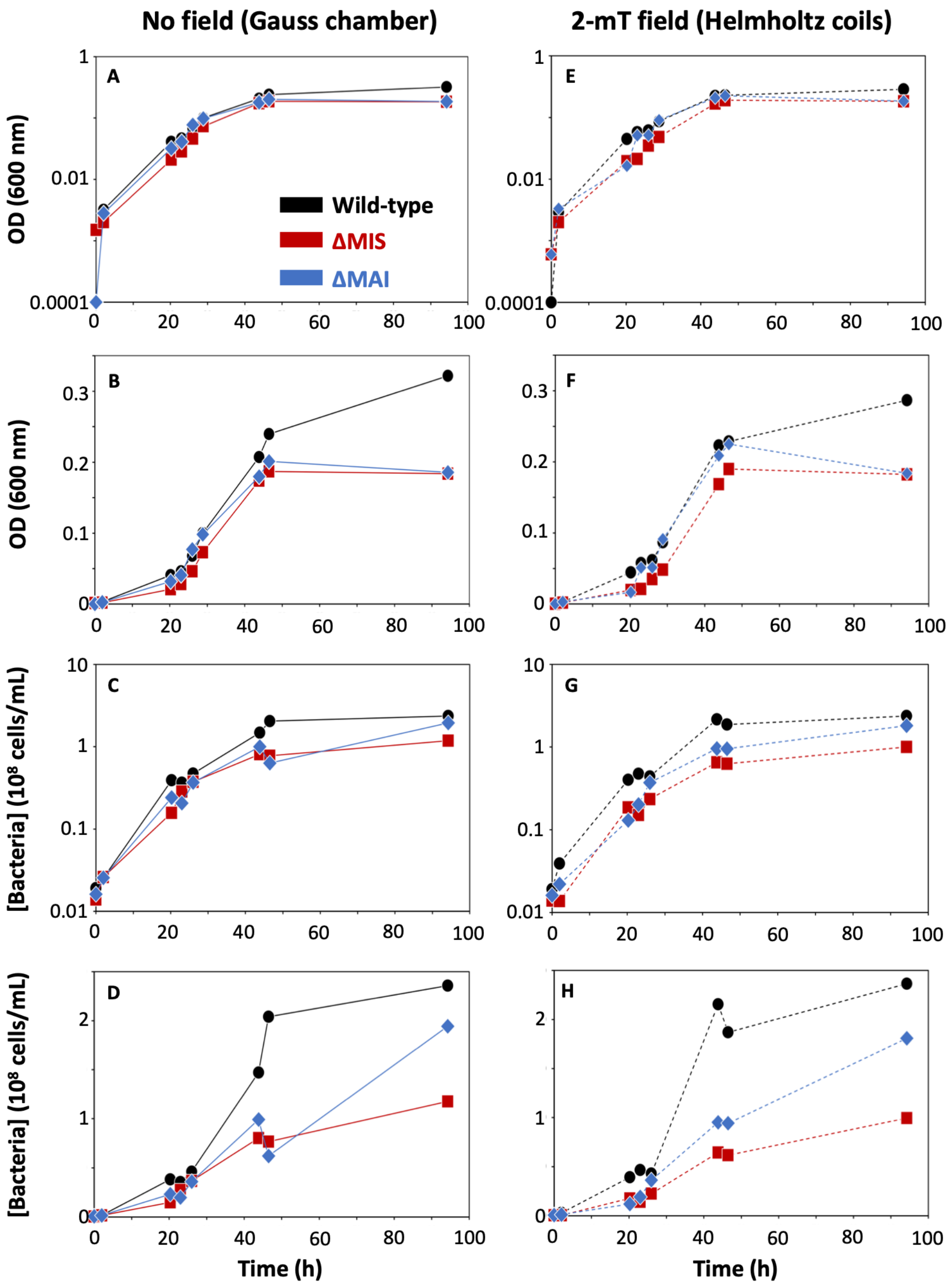


**Fig. S7.** First replicate of wild-type (black circles), ΔMIS (red squares) and ΔMAI (blue diamonds) cultures in (A-D) zero (solid curves) and (E-H) 2-mT (dashed curves) fields. (A, B, E, F) Optical density measured at 600 nm. (C, D, H, G) Bacterial concentrations measured from flow-cytometry.

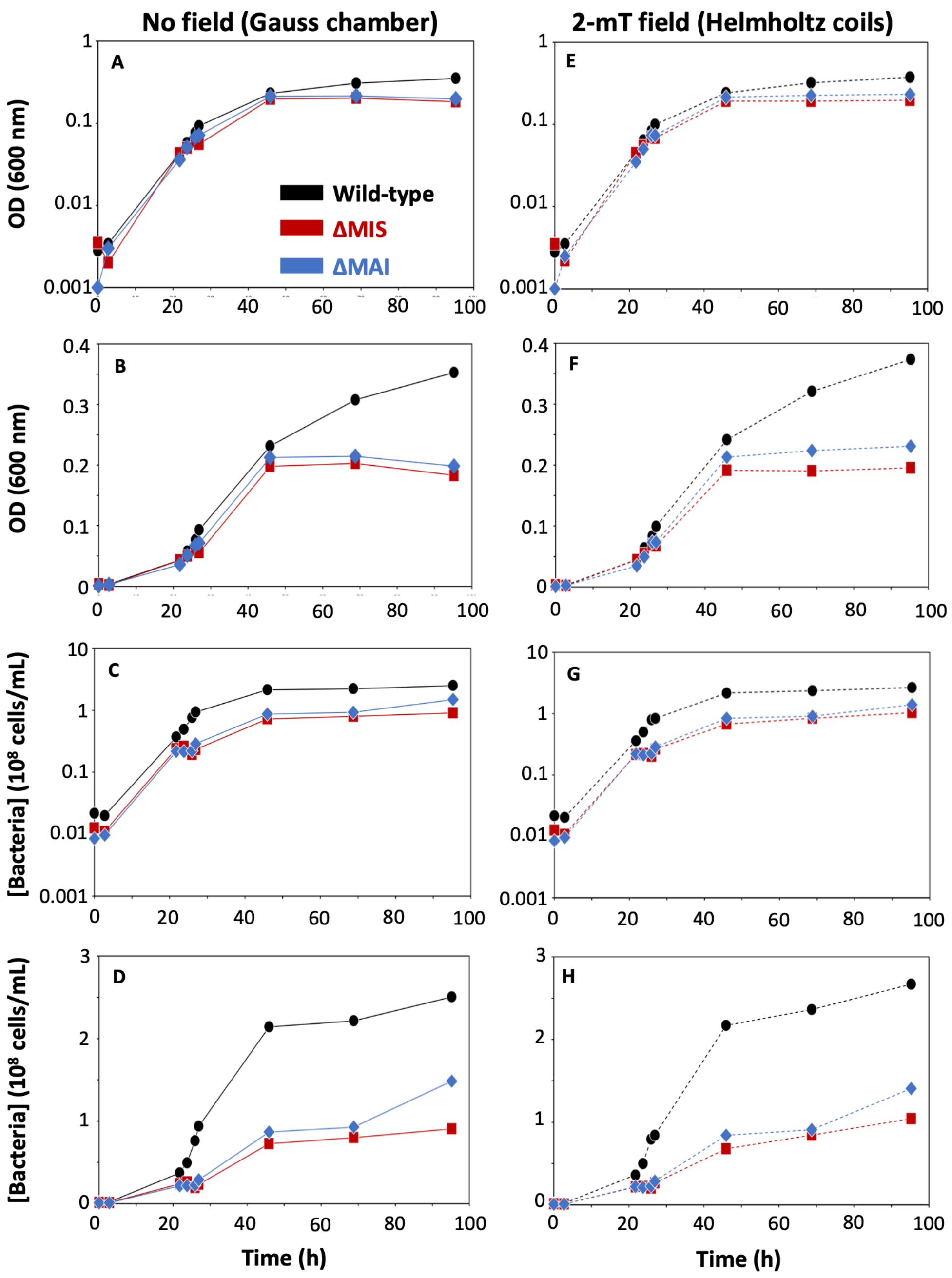


**Fig. S8.** Second replicate of wild-type (black circles), ΔMIS (red squares) and ΔMAI (blue diamonds) cultures in (A-D) zero (solid curves) and (E-H) 2-mT (dashed curves) fields. (A, B, E, F) Optical density measured at 600 nm. (C, D, H, G) Bacterial concentrations measured from flow-cytometry.

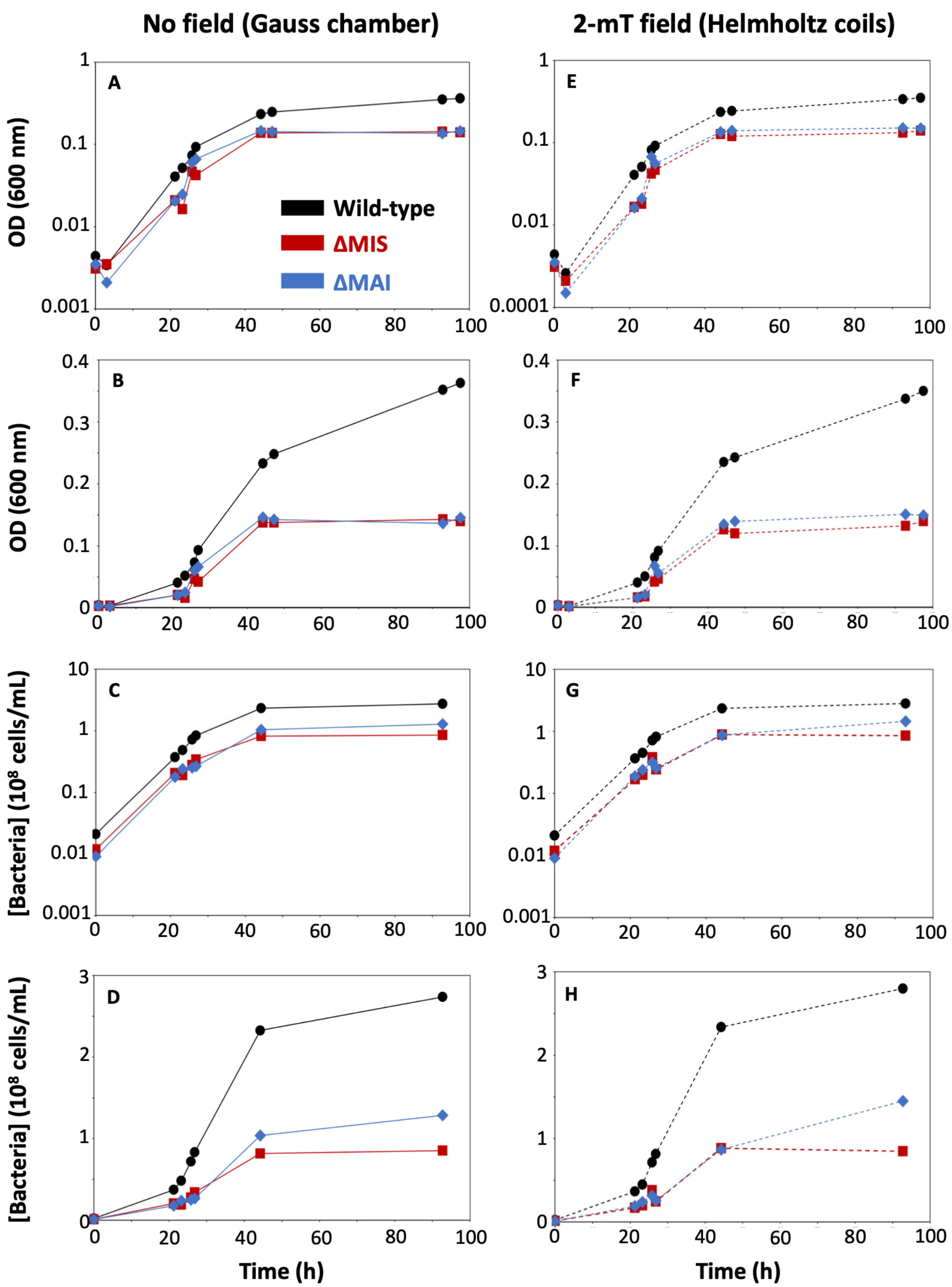


**Fig. S9.** Third replicate of wild-type (black circles), ΔMIS (red squares) and ΔMAI (blue diamonds) cultures in (A-D) zero (solid curves) and (E-H) 2-mT (dashed curves) fields. (A, B, E, F) Optical density measured at 600 nm. (C, D, G, H) Bacterial concentrations measured from flow-cytometry.

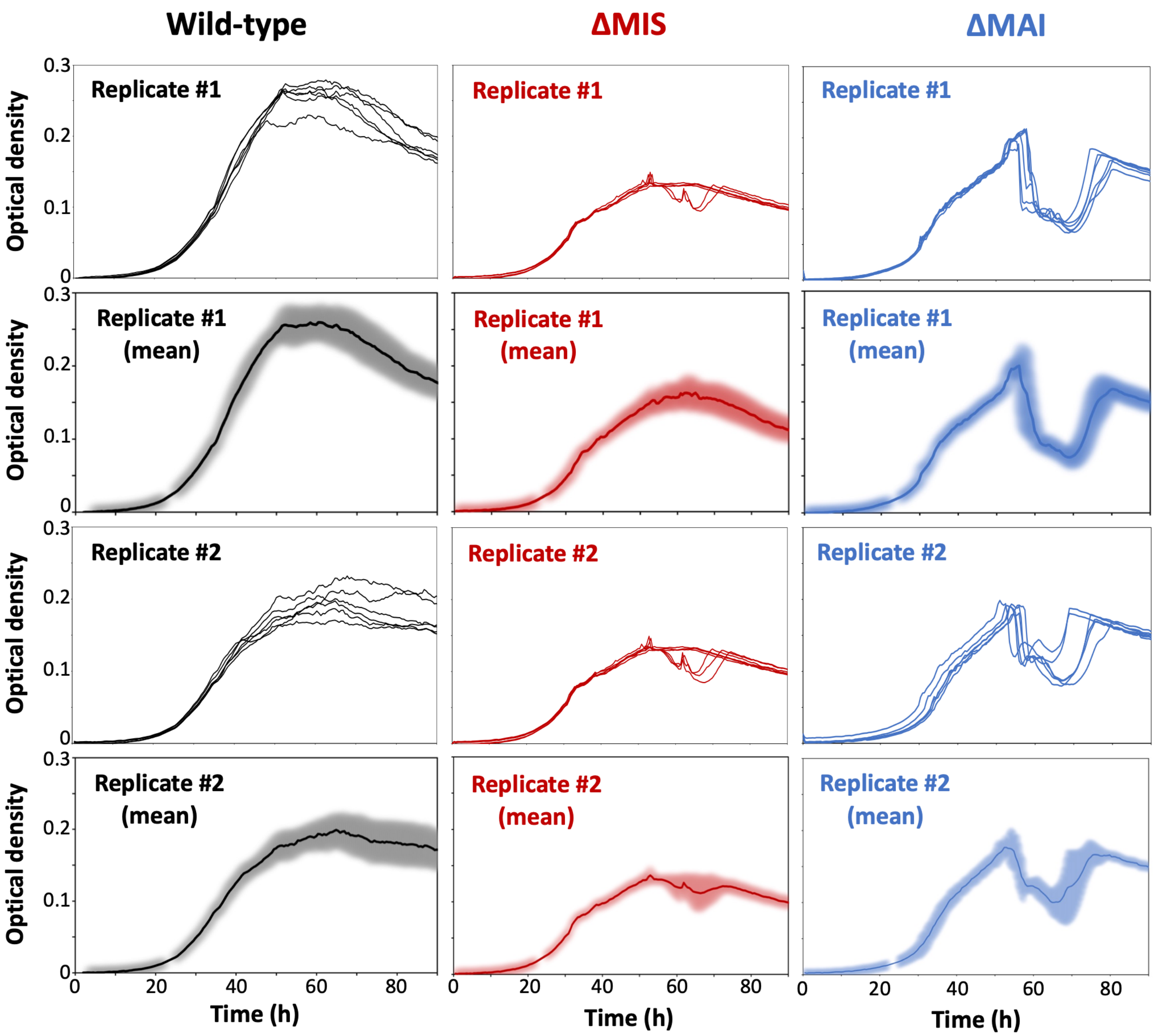


**Fig. S10.** Cultures of the three AMB-1 strain in a microplate reader and monitored from measurement of optical density at 600 nm. Each replicate corresponds to 6 cultures from the same inoculum. Shaded areas in the mean curves represent 1 SD.

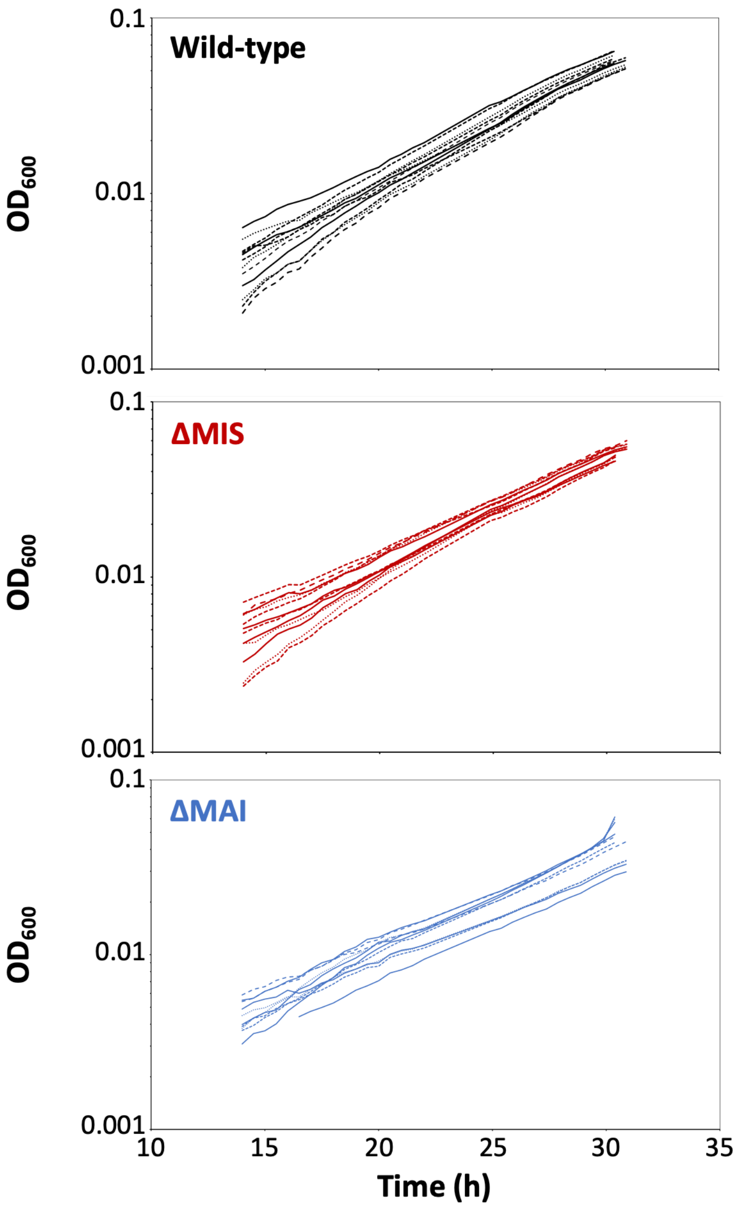


**Fig. S11.** Growth curves of wild-type, ΔMIS and ΔMAI cultures represented as log scale during the exponential phase between 14 and 32h of growth, and used to calculate doubling times (Fig. 2h). All curves were extracted from Extended Fig. 11.

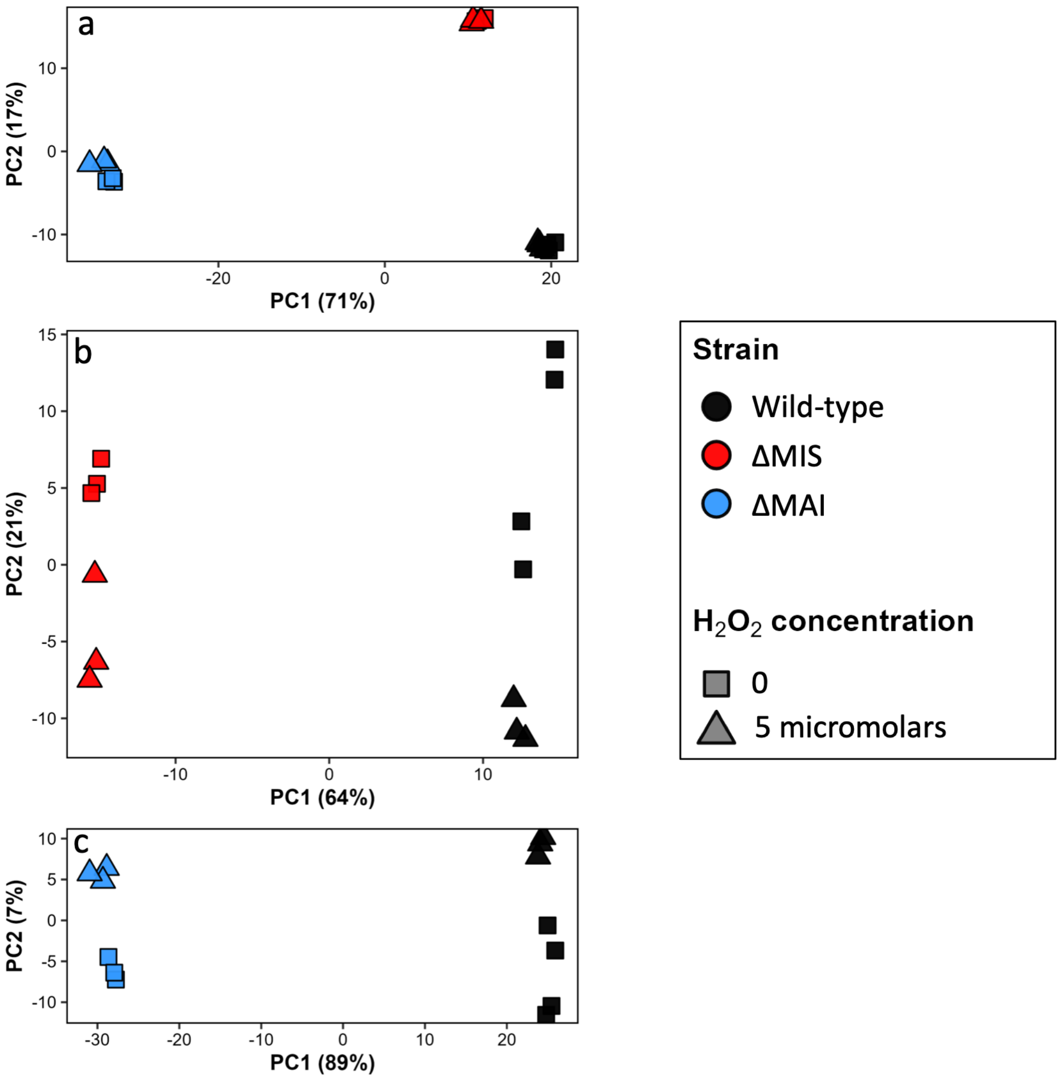


**Fig. S12.** Principal component analyses (PCA) of RNA-seq. (A) Main PCA projection of all samples onto the first two principal components. (B) PCA projection of wild-type and ΔMIS samples. (C) PCA projection of wild-type and ΔMIS samples.

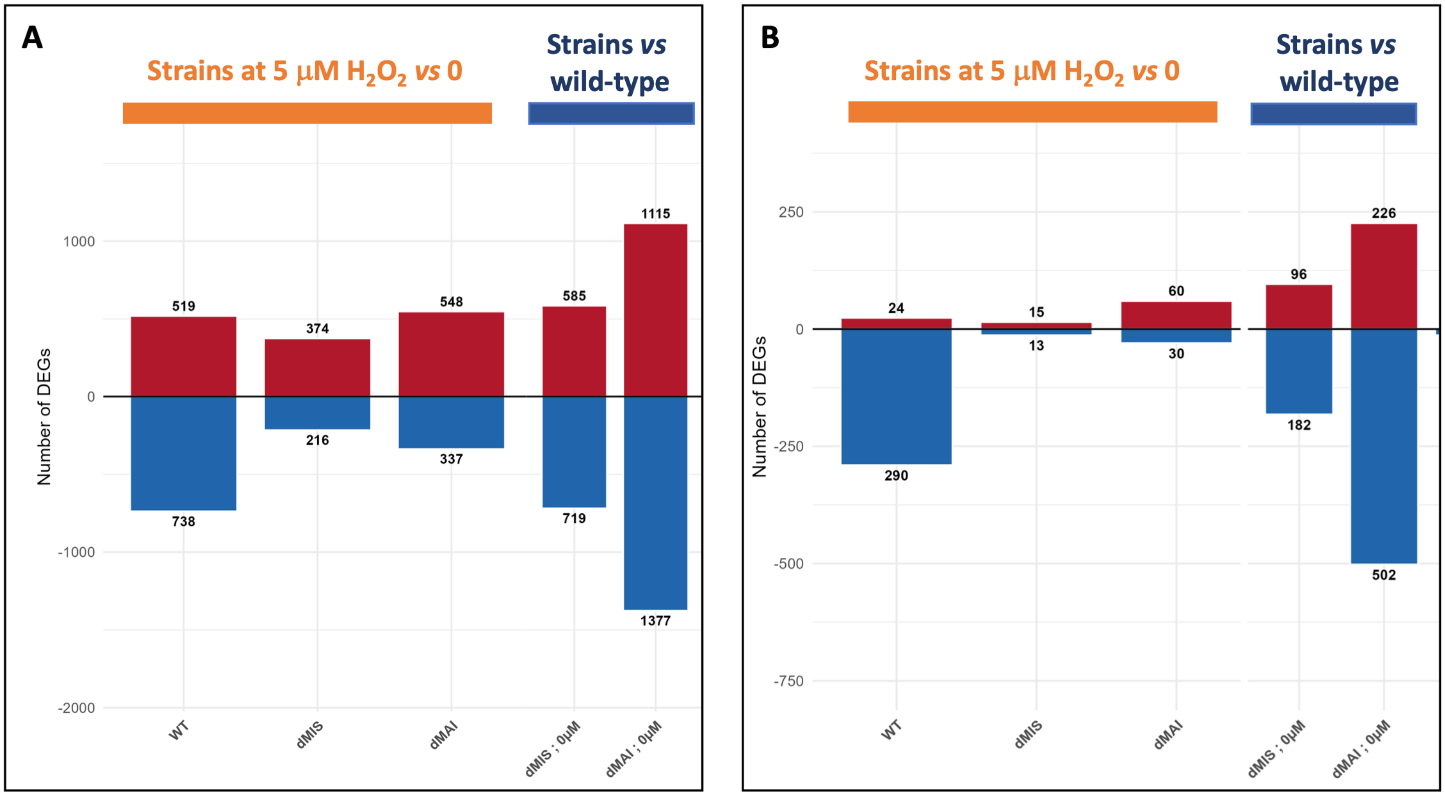


**Fig. S13.** Differentially expressed genes (DEG) in bacteria cultivated with 5 µM of $H_2O_2$ (yellow – normalized to no $H_2O_2$) and in the two mutant strains normalized to wild-type (dark blue – no $H_2O_2$). (A) shows all genes with an adjusted p-value < 0.05, and (B) shows genes with an adjusted p-value < 0.05 and a fold change (FC) of at least $|log_2FC| > 1$.

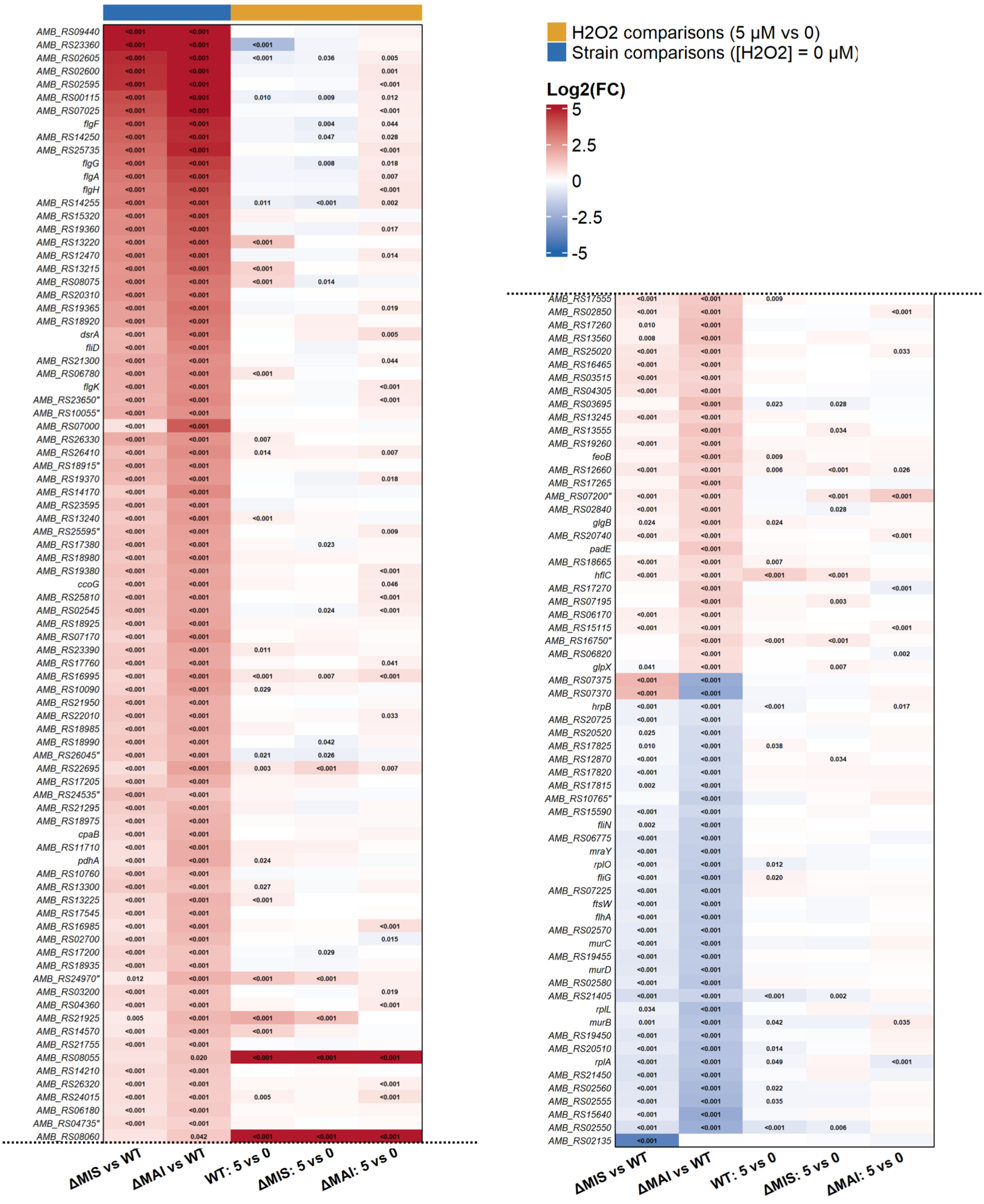


**Fig. S14.** 150 most differentially expressed genes (padj < 0.05, log2(FC) > 1) in RNA-seq analyses. Genes from the MIS and the MAI were excluded. Detailed list with HMM products is given in Extended Table 2.

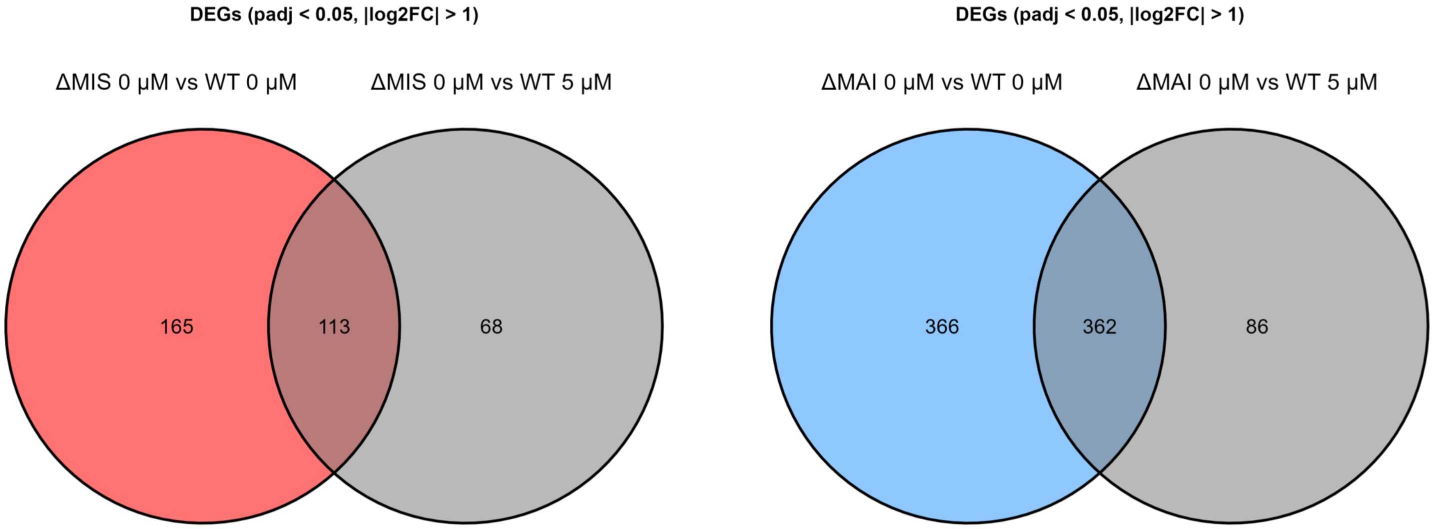


**Fig. S15.** Venn diagrams representing the number of differentially expressed genes (DEG) in ΔMIS (left panel) and ΔMAI (right panel) bacteria cultivated without added $H_2O_2$ relative to wild-type bacteria cultivated with and without added $H_2O_2$.

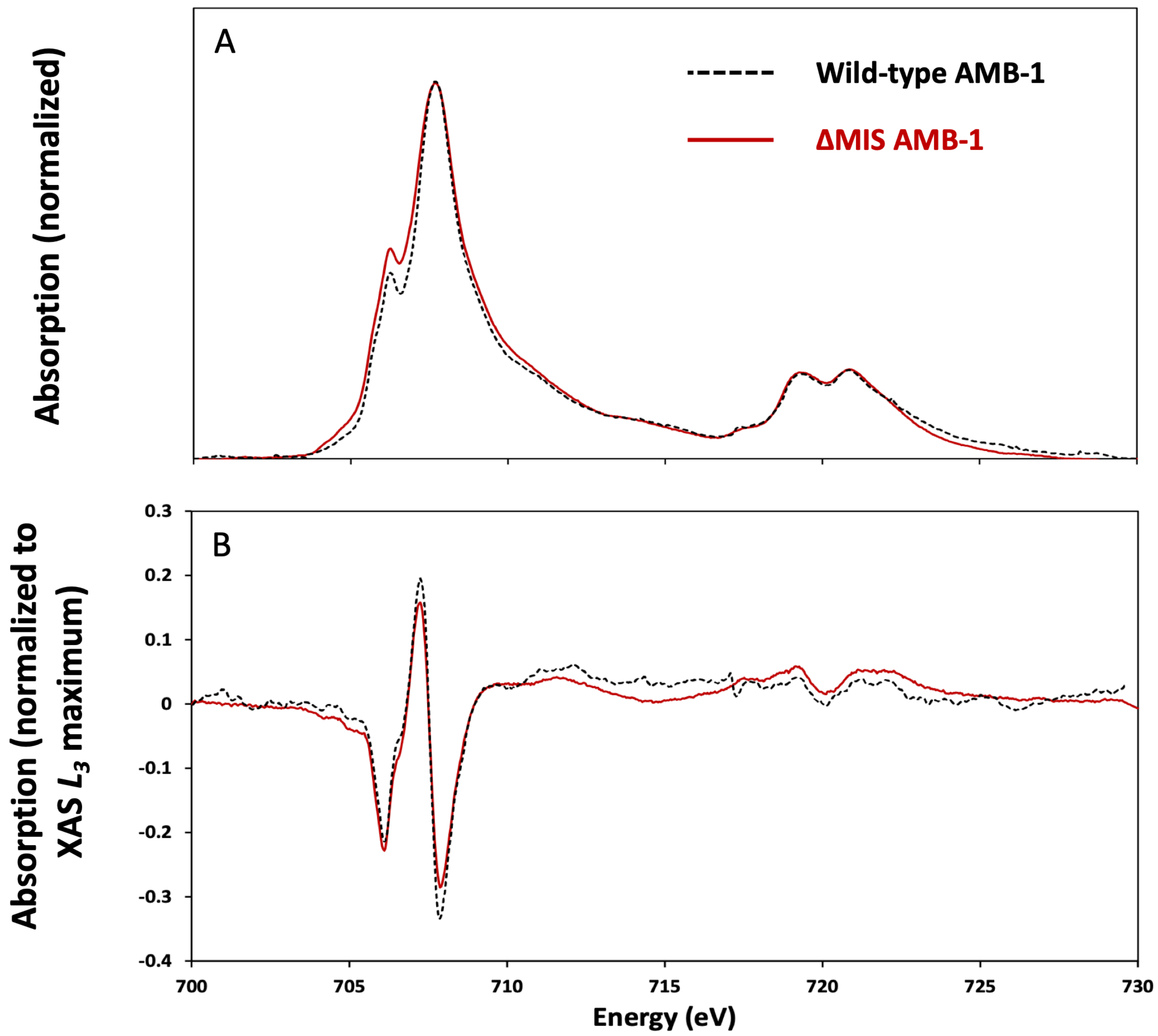


**Fig. S16.** X-ray absorption spectroscopy (XAS) (A) and X-ray circular magnetic dichroism (XMCD) (B) spectra obtained at the Fe $L_{2,3}$-edges from wild-type and ΔMIS bacteria.

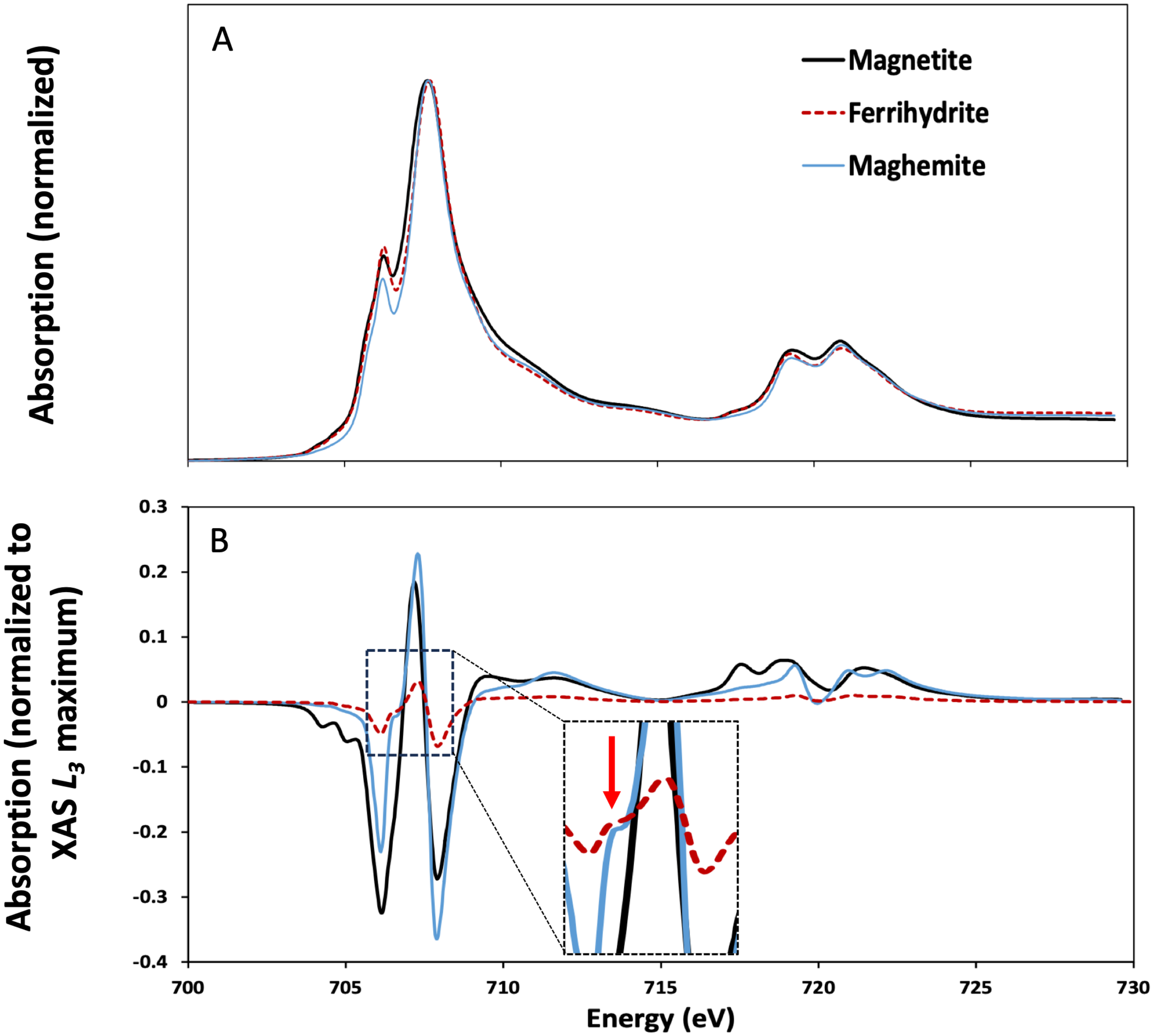


**Fig. S17.** X-ray absorption spectroscopy (XAS) (A) and X-ray circular magnetic dichroism (XMCD) (B) spectra obtained at the Fe $L_{2,3}$-edges from chemical magnetite, maghemite, and ferrihydrite. Inset in **b** magnifies the feature at 706.6 eV indicated by the red arrow.

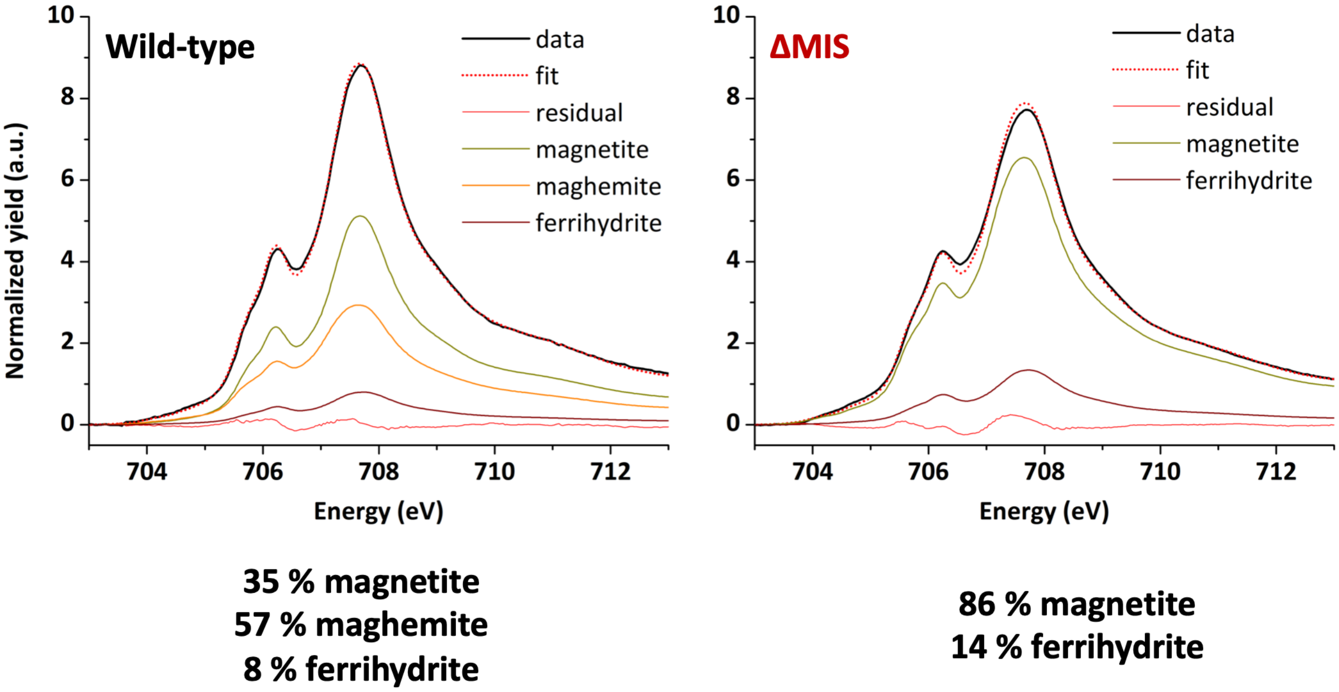


**Fig. S18.** X-ray absorption spectroscopy (XAS) spectra and linear combination fitting obtained at the Fe $L_3$-edge from wild-type and ΔMIS.

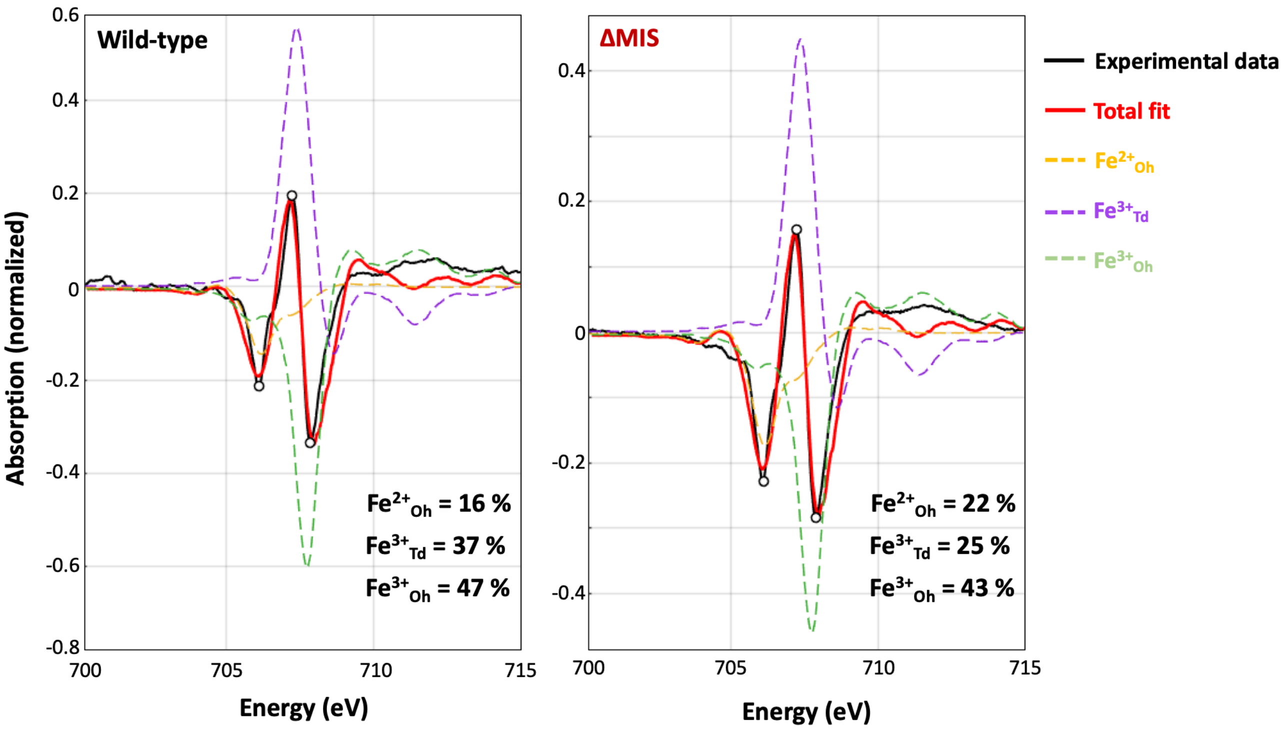


**Fig. S19.** X-ray circular magnetic dichroism (XMCD) spectra and linear combination fitting obtained at the Fe $L_3$-edge from wild-type and ΔMIS.

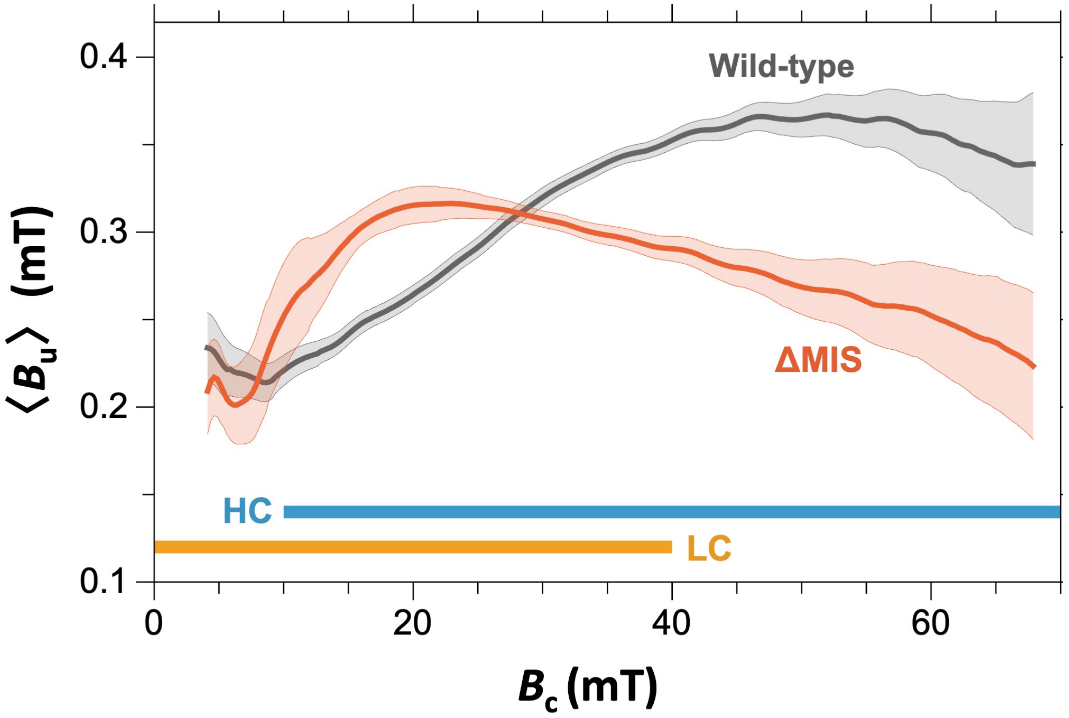


**Fig. S20.** Vertical offset of the central ridge as a function of $B_c$ for wild-type and ΔMIS, as determined with the VARIFORC processing software. The shaded color bands represent the 1-σ uncertainty. Horizontal bars represent the ranges of the LC and HC coercivity components, respectively.

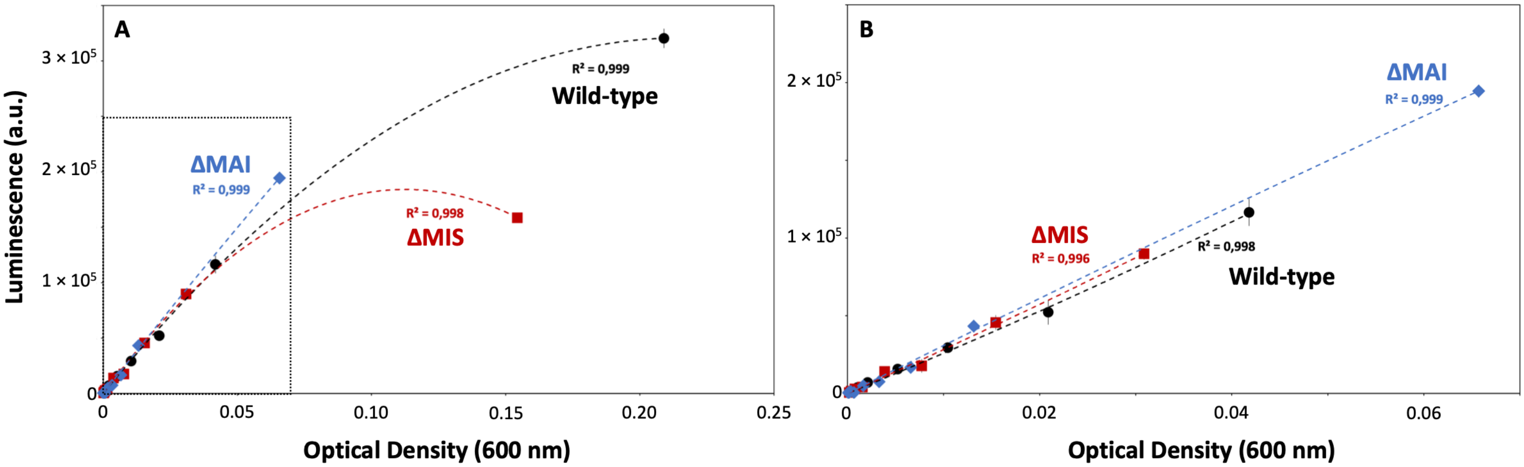


**Fig. S21.** Calibration of ATP measurements in the three AMB-1 strains. Inset in (A) is magnified in (B). Optical density (600 nm) corresponds to the optical density of the tested cell suspensions.

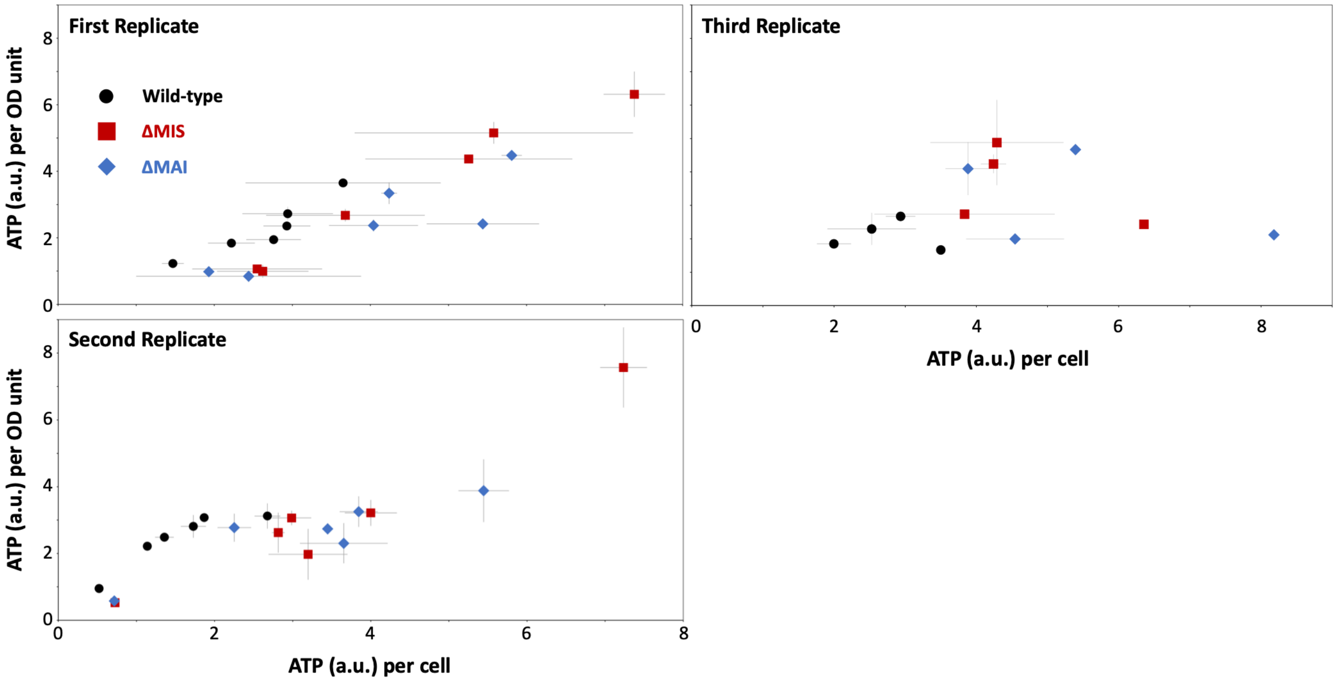


**Fig. S22.** Intracellular ATP content of wild-type (black circles), ΔMIS (red squares) and ΔMAI (blue diamonds) AMB-1 expressed as ATP per cell and ATP per optical density (OD) unit measured at different time points during exponential phase. All data correspond to the mean value of two replicates (one exposed to a 2 mT field and one cultivated in a zero-field environment ±1SD).

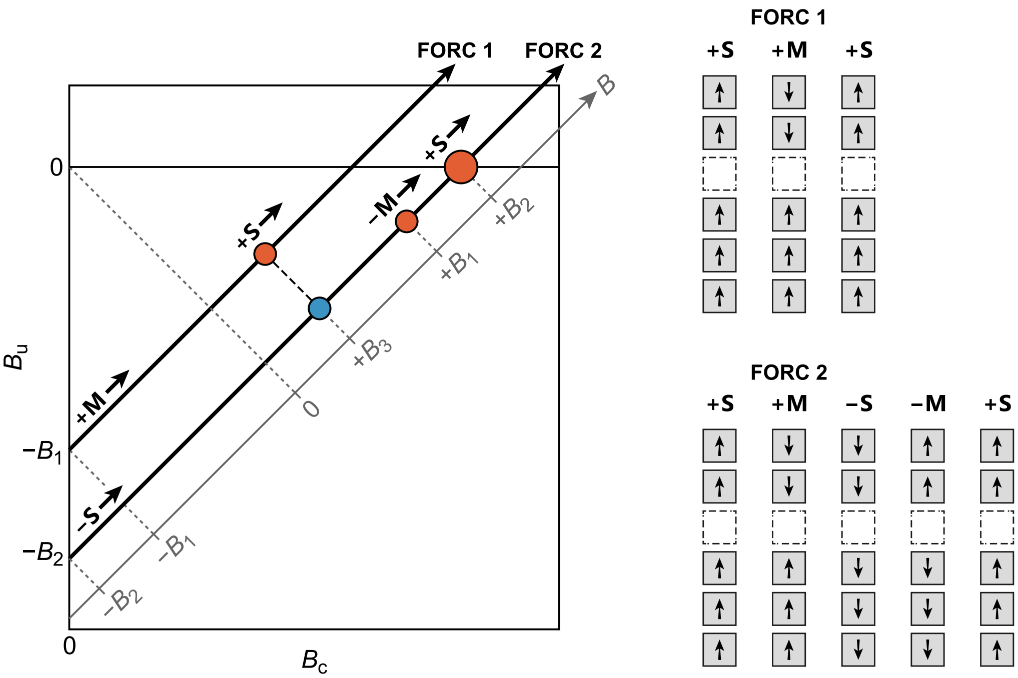


**Fig. S23.** Conceptual model for the FORC signature of a fragmented chain. **Right:** The chain model consists of a short fragment and a long fragment (filled magnetosomes with arrows indicating the direction of the magnetic moment) separated by a gap (dashed square). Based on the stronger uniaxial anisotropy of the longer fragment and the weakened magnetic coupling between the two fragments, the chain has two pairs of magnetic states denoted by ±S for single polarity (both fragments share the same polarity), and ±M for mixed polarity (the fragments have opposite polarity), with the sign indicating the direction of the net magnetic moment with respect to the chain axis. **Left:** Schematic FORC diagram showing the discrete contributions arising from a single fragmented chain with the properties depicted on the right. FORC 1 and FORC 2 are two curves starting just after the +M and the −S states have been nucleated from positive saturation, respectively. These curves form the diagonal traces indicated by the thick black lines in the coordinate system of the FORC diagram. The red dot along FORC 1 indicates the magnetization jump caused by the +M → +S transition, whose derivative produce a positive spike in the FORC space. The first blue dot along FORC 2 depicts the same transition, which appears as a negative spike because of the difference with FORC 1. The other two red dots on FORC 2 arise from the −S → −M transition, followed by −M → +S. The size of the dots is proportional to the spike amplitude.

```
AMB_RS01265_[P.magneticum]_WP_043743099.1/1-295     1 M-NLKQLEYFVRVAELGSFSKAAMVLDIAQPALSRQVRL  38
OxyR-like_[M.gryphiswaldense]_WP_106002419.1/1-240   1 MISFRQLQCLAALAETLHFRRAAERLHVSQPALSAQISQ  39
OXYR_ECOLI_sp|P0ACQ4|/1-305                          1 M-NIRDLEYLVALAEHRHFRRAADSCHVSQPTLSGQIRK  38

AMB_RS01265_[P.magneticum]_WP_043743099.1/1-295    39 LETDLRANLLHRTGRGVALTEAGQRLFDHAVGILQSVER  77
OxyR-like_[M.gryphiswaldense]_WP_106002419.1/1-240  40 LEEHLGVMLVERTRRKVLLTPIGRDMAARAKTVLRDVAE  78
OXYR_ECOLI_sp|P0ACQ4|/1-305                         39 LEDELGVMLLERTSRKVLFTQAGMLLVDQARTVLREVKV  77

AMB_RS01265_[P.magneticum]_WP_043743099.1/1-295    78 ATEDLESARGE-PAGRIVIGLPPSIGRRLIVPLVEGFRQ 115
OxyR-like_[M.gryphiswaldense]_WP_106002419.1/1-240  79 L-EDLARQAQAPLSGTLRVGVLRTLGPYLLPHILPELRQ 116
OXYR_ECOLI_sp|P0ACQ4|/1-305                         78 L-KEMASQQGETMSGPLHIGLIPTVGPYLLPHIIPMLHQ 115

AMB_RS01265_[P.magneticum]_WP_043743099.1/1-295   116 TLPRARLAIVEGLSTHVTEWIATGRVDLGLVH-NPEPNP 153
OxyR-like_[M.gryphiswaldense]_WP_106002419.1/1-240 117 RFPELKLYLREEPRERLLAELAQGDLDILLISAAPEGDD 155
OXYR_ECOLI_sp|P0ACQ4|/1-305                        116 TFPKLEMYLHEAQTHQLLAQLDSGKLDCVILALVKESEA 154

AMB_RS01265_[P.magneticum]_WP_043743099.1/1-295   154 AIEVSRLLDEPLGLVGPAGDVP---ATLRWAELGSYPLI 189
OxyR-like_[M.gryphiswaldense]_WP_106002419.1/1-240 156 HLTVTPLFHEPLWLTMPLGHRLAKRAKLAPADLAGENLI 194
OXYR_ECOLI_sp|P0ACQ4|/1-305                        155 FIEV-PLFDEPMLLAIYEDHPWANRECVPMADLAGEKLL 192

AMB_RS01265_[P.magneticum]_WP_043743099.1/1-295   190 LPERGHAMRRLLEAQAAFAEVRLKVVMEVSSIPSILDLV 228
OxyR-like_[M.gryphiswaldense]_WP_106002419.1/1-240 195 MLEVGDGMRDPALDLCRRSGATEHPDFRATSLDSLRQMV 233
OXYR_ECOLI_sp|P0ACQ4|/1-305                        193 MLEDGHCLRDQAMGFCFEAGADEDTHFRATSLETLRNMV 231

AMB_RS01265_[P.magneticum]_WP_043743099.1/1-295   229 AAGYGHAVLTETALASWRRAGGFSLRRLVEPSVASTLFM 267
OxyR-like_[M.gryphiswaldense]_WP_106002419.1/1-240 234 ATGLGTT-------------------------------- 240
OXYR_ECOLI_sp|P0ACQ4|/1-305                        232 AAGSGITLLPALAVPPERKRDGVVYLPCIKPEPRRTIGL 270

AMB_RS01265_[P.magneticum]_WP_043743099.1/1-295   268 AVSAQKPITPLGRRAMALLQDLVKTEMG----------- 295
OxyR-like_[M.gryphiswaldense]_WP_106002419.1/1-240     ---------------------------------------
OXYR_ECOLI_sp|P0ACQ4|/1-305                        271 V---YRPGSPLRSRYEQL-AEAIRARMDGHFDKVLKQAV 305
```

**Fig. S24.** Multiple sequence alignment of OxyR and OxyR-like proteins from *E. coli* K12, *M. gryphiswaldense*, and *P. magneticum* AMB-1. Blue shading indicates residues with > 50% identity.

#### Supporting Tables

**Table S1.** Magnetite nanoparticle size, nanoparticle shape factor (defined as the ratio width / length), and number of chain fragments in wild-type and ΔMIS AMB-1 (± 1 SD) measured from electron microscopy images using imageJ software.

| | **Wild-type** | **ΔMIS** |
|---|---|---|
| Magnetite length (nm) | 43.01 ± 14.05 | 39.61 ± 14.77 |
| Magnetite shape factor | 0.83 ± 0.11 | 0.91 ± 0.09 |
| Number of magnetite per cell | 20.37 ± 5.75 | 18.80 ± 6.32 |
| Number of chain fragments per cell | 4.33 ± 1.07 | 1.28 ± 0.54 |

**Table S2.** Coercivity distribution parameters of the components LC and HC obtained from the central ridge, with 1-σ uncertainty. $M_{cr}$ — Component magnetization in $mAm^2/kg$, $M_{cr}/M_{\mathrm{tot}}$ — relative contribution of component, $B_{1/2}$ — Median field in mT, $\sigma$ — Standard deviation of field on a $\log_{10}$ scale.

| | **Wild-type LC** | **Wild-type HC** | **ΔMIS LC** | **ΔMIS HC** |
|---|---|---|---|---|
| $M_{cr}$ | 145 ± 8 | 190 ± 8 | 50 ± 3 | 159 ± 3 |
| $M_{cr}/M_{\mathrm{tot}}$ | 0.44 ± 0.02 | 0.56 ± 0.02 | 0.24 ± 0.01 | 0.76 ± 0.01 |
| $B_{1/2}$ | 19 ± 0.5 | 35 ± 0.3 | 19 ± 0.5 | 29 ± 0.06 |
| $\sigma$ | 0.279 ± 0.003 | 0.111 ± 0.002 | 0.279 ± 0.003 | 0.133 ± 0.002 |

**Table S3.** Detailed list of the 150 most differentially expressed genes discussed in Extended Fig. 14.

| Gene id | Gene name | HMM product | padj |
|---|---|---|---|
| AMB_RS00115 | AMB_RS00115 | hypothetical protein | 0.00E+00 |
| AMB_RS01195 | AMB_RS01195 | TNT domain-containing protein | 2.82E-36 |
| AMB_RS02135 | AMB_RS02135 | acyltransferase family protein | 1.52E-224 |
| AMB_RS02545 | AMB_RS02545 | hypothetical protein | 5.72E-215 |
| AMB_RS02550 | AMB_RS02550 | hypothetical protein | 1.74E-79 |
| AMB_RS02555 | AMB_RS02555 | MinD/ParA family protein | 7.93E-75 |
| AMB_RS02560 | AMB_RS02560 | GTP-binding protein | 5.23E-83 |
| AMB_RS02565 | flhA | flagellar biosynthesis protein | 2.46E-53 |
| AMB_RS02570 | AMB_RS02570 | sigma-54-dependent transcriptional regulator FlbD | 2.70E-63 |
| AMB_RS02575 | fliN | flagellar motor switch protein FliN | 1.95E-36 |
| AMB_RS02580 | AMB_RS02580 | FliH/SctL family protein | 2.45E-54 |
| AMB_RS02585 | fliG | flagellar motor switch protein | 3.46E-63 |
| AMB_RS02595 | flgE | flagellar hook-basal body complex protein | 0.00E+00 |
| AMB_RS02600 | flgD | flagellar hook assembly protein | 0.00E+00 |
| AMB_RS02605 | fliK | flagellar hook-length control protein | 0.00E+00 |
| AMB_RS02700 | AMB_RS02700 | transglycosylase domain-containing protein | 1.60E-45 |
| AMB_RS02840 | AMB_RS02840 | response regulator transcription factor | 1.83E-39 |
| AMB_RS02850 | AMB_RS02850 | branched-chain amino acid aminotransferase | 7.72E-131 |
| AMB_RS03200 | AMB_RS03200 | caspase family protein | 1.41E-46 |
| AMB_RS03695 | AMB_RS03695 | hypothetical protein | 4.48E-44 |
| AMB_RS04305 | AMB_RS04305 | HD domain-containing phosphohydrolase | 7.13E-37 |
| AMB_RS04360 | AMB_RS04360 | thiosulfate oxidation carrier protein SoxY | 1.71E-54 |
| AMB_RS04520 | AMB_RS04520 | RT0821/Lpp0805 family surface protein | 8.27E-38 |
| AMB_RS05320 | AMB_RS05320 | IS3 family transposase | 1.07E-281 |
| AMB_RS06180 | AMB_RS06180 | methyl-accepting chemotaxis protein | 5.07E-53 |
| AMB_RS06490 | AMB_RS06490 | hypothetical protein | 3.85E-36 |
| AMB_RS06700 | nsrR | nitric oxide-sensing transcriptional repressor NsrR | 1.21E-37 |
| AMB_RS06775 | AMB_RS06775 | phosphorelay signal transduction system | 1.02E-54 |
| AMB_RS06780 | AMB_RS06780 | hypothetical protein | 8.31E-147 |
| AMB_RS06795 | AMB_RS06795 | hypothetical protein | 2.02E-45 |
| AMB_RS07000 | AMB_RS07000 | acetyl-CoA hydrolase/transferase family protein | 0.00E+00 |
| AMB_RS07025 | flgE | flagellar hook-basal body complex protein | 0.00E+00 |
| AMB_RS07170 | AMB_RS07170 | bacteriohemerythrin | 7.53E-85 |
| AMB_RS07195 | AMB_RS07195 | AMP-binding protein | 9.15E-34 |
| AMB_RS07200 | AMB_RS07200 | (2Fe-2S)-binding protein | 1.08E-173 |
| AMB_RS07370 | AMB_RS07370 | phage tail protein | 2.68E-97 |
| AMB_RS07375 | AMB_RS07375 | phage tail protein | 2.61E-92 |
| AMB_RS07415 | AMB_RS07415 | hypothetical protein | 1.55E-45 |
| AMB_RS07420 | AMB_RS07420 | nitrous oxide reductase accessory protein NosL | 1.06E-43 |

| | | | |
|---|---|---|---|
| AMB_RS08075 | AMB_RS08075 | hypothetical protein | 8.11E-169 |
| AMB_RS08475 | padE | enylglyoxylate dehydrogenase (acylating) activity | 3.17E-40 |
| AMB_RS09185 | AMB_RS09185 | hypothetical protein | 4.33E-48 |
| AMB_RS09440 | AMB_RS09440 | hypothetical protein | 0.00E+00 |
| AMB_RS10055 | AMB_RS10055 | VCBS domain-containing protein | 1.79E-44 |
| AMB_RS10090 | AMB_RS10090 | ATP binding | 2.56E-127 |
| AMB_RS10760 | AMB_RS10760 | hypothetical protein | 3.76E-43 |
| AMB_RS11130 | AMB_RS11130 | transglycosylase SLT domain-containing protein | 7.10E-49 |
| AMB_RS11380 | AMB_RS11380 | bacteriohemerythrin | 9.55E-41 |
| AMB_RS11705 | pdhA | pyruvate dehydrogenase | 1.17E-55 |
| AMB_RS11710 | AMB_RS11710 | pyruvate dehydrogenase (acetyl-transferring) activity | 1.41E-38 |
| AMB_RS12470 | AMB_RS12470 | hypothetical protein | 5.60E-162 |
| AMB_RS12870 | AMB_RS12870 | bifunctional acetate--CoA ligase family protein | 1.21E-118 |
| AMB_RS13215 | AMB_RS13215 | cation acetate symporter | 5.40E-209 |
| AMB_RS13220 | AMB_RS13220 | DUF485 domain-containing protein | 2.40E-179 |
| AMB_RS13225 | AMB_RS13225 | hypothetical protein | 2.24E-80 |
| AMB_RS13240 | AMB_RS13240 | acyl-CoA synthetase | 6.52E-193 |
| AMB_RS13300 | AMB_RS13300 | phosphorelay signal transduction system | 1.60E-52 |
| AMB_RS13555 | AMB_RS13555 | phosphoribulokinase | 9.75E-36 |
| AMB_RS13560 | AMB_RS13560 | ribulose-bisphosphate carboxylase | 2.57E-47 |
| AMB_RS13735 | feoB | ferrous iron transmembrane transporter activity | 4.24E-43 |
| AMB_RS14170 | AMB_RS14170 | hypothetical protein | 1.20E-130 |
| AMB_RS14210 | AMB_RS14210 | methyl-accepting chemotaxis protein | 1.02E-35 |
| AMB_RS14250 | AMB_RS14250 | hypothetical protein | 5.50E-257 |
| AMB_RS14255 | AMB_RS14255 | hypothetical protein | 7.93E-221 |
| AMB_RS14375 | fliD | flagellar filament capping protein | 4.05E-63 |
| AMB_RS14570 | AMB_RS14570 | response regulator transcription factor | 1.38E-41 |
| AMB_RS14595 | AMB_RS14595 | histone deacetylase family protein | 1.22E-41 |
| AMB_RS15320 | AMB_RS15320 | hypothetical protein | 8.29E-87 |
| AMB_RS15590 | AMB_RS15590 | YgiQ family radical SAM protein | 3.04E-50 |
| AMB_RS15640 | AMB_RS15640 | hypothetical protein | 1.09E-103 |
| AMB_RS15665 | AMB_RS15665 | adenylate kinase | 3.42E-38 |
| AMB_RS15670 | secY | preprotein translocase subunit SecY | 1.76E-37 |
| AMB_RS15675 | rplO | 50S ribosomal protein L15 | 2.71E-43 |
| AMB_RS15820 | rplL | 50S ribosomal protein L7/L12 | 6.30E-48 |
| AMB_RS15825 | rplJ | 50S ribosomal protein L10 | 3.85E-38 |
| AMB_RS15830 | rplA | 50S ribosomal protein L1 | 2.92E-57 |
| AMB_RS15835 | rplK | 50S ribosomal protein L11 | 1.92E-42 |
| AMB_RS16775 | tnpC | transposase activity | 5.61E-34 |
| AMB_RS16975 | ccoG | cytochrome c oxidase accessory protein CcoG | 1.22E-42 |
| AMB_RS16985 | AMB_RS16985 | DUF6691 family protein | 4.82E-56 |
| AMB_RS16990 | AMB_RS16990 | YeeE/YedE family protein | 6.52E-36 |
| AMB_RS16995 | AMB_RS16995 | peroxiredoxin activity | 3.70E-88 |

| | | | |
|---|---|---|---|
| AMB_RS17015 | cobA | uroporphyrinogen-III C-methyltransferase | 4.94E-53 |
| AMB_RS17035 | dsrA | dissimilatory-type sulfite reductase subunit alpha | 5.98E-64 |
| AMB_RS17040 | dsrB | dissimilatory-type sulfite reductase subunit beta" | 2.60E-41 |
| AMB_RS17045 | tusD | sulfurtransferase complex subunit | 4.28E-42 |
| AMB_RS17200 | AMB_RS17200 | two-component system response regulator TorR | 9.24E-64 |
| AMB_RS17205 | AMB_RS17205 | hypothetical protein | 9.66E-41 |
| AMB_RS17260 | AMB_RS17260 | efflux RND transporter permease subunit | 1.34E-42 |
| AMB_RS17265 | AMB_RS17265 | efflux RND transporter periplasmic adaptor subunit | 3.54E-40 |
| AMB_RS17270 | AMB_RS17270 | TetR/AcrR family transcriptional regulato | 4.23E-40 |
| AMB_RS17380 | AMB_RS17380 | FIST signal transduction protein | 2.79E-228 |
| AMB_RS17545 | AMB_RS17545 | methyl-accepting chemotaxis protein | 3.60E-90 |
| AMB_RS17555 | AMB_RS17555 | hypothetical protein | 1.82E-105 |
| AMB_RS17655 | hflC | protease modulator HflC | 1.19E-36 |
| AMB_RS17670 | flgH | flagellar basal body L-ring protein | 3.47E-267 |
| AMB_RS17675 | flgA | flagellar basal body P-ring formation chaperone | 5.56E-234 |
| AMB_RS17680 | flgG | flagellar basal-body rod protein | 1.90E-247 |
| AMB_RS17685 | flgF | flagellar basal-body rod protein | 0.00E+00 |
| AMB_RS17760 | AMB_RS17760 | hypothetical protein | 1.55E-64 |
| AMB_RS17820 | AMB_RS17820 | MFS transporter | 6.96E-38 |
| AMB_RS17825 | AMB_RS17825 | hypothetical protein | 6.96E-38 |
| AMB_RS17925 | AMB_RS17925 | molybdopterin oxidoreductase family protein | 9.74E-48 |
| AMB_RS18915 | AMB_RS18915 | lp family type IVb pilin | 4.80E-44 |
| AMB_RS18920 | AMB_RS18920 | hypothetical protein | 2.19E-107 |
| AMB_RS18925 | AMB_RS18925 | A24 family peptidase | 1.78E-77 |
| AMB_RS18930 | cpaB | Flp pilus assembly protein | 1.34E-69 |
| AMB_RS18935 | AMB_RS18935 | type II and III secretion system protein family | 2.03E-42 |
| AMB_RS18975 | AMB_RS18975 | pilus assembly protein N-terminal domain-containing protein | 3.41E-61 |
| AMB_RS18980 | AMB_RS18980 | TadE/TadG family type IV pilus assembly protein | 3.94E-75 |
| AMB_RS18985 | AMB_RS18985 | TadE/TadG family type IV pilus assembly protein | 3.58E-74 |
| AMB_RS18990 | AMB_RS18990 | TadE/TadG family type IV pilus assembly protein | 2.86E-49 |
| AMB_RS19260 | AMB_RS19260 | methyl accepting chemotaxis protein complex | 4.06E-42 |
| AMB_RS19360 | flgI | flagellar basal body P-ring protein | 3.22E-198 |
| AMB_RS19365 | flgJ | flagellar rod assembly protein/muramidase | 8.50E-147 |
| AMB_RS19370 | AMB_RS19370 | flagellar protein FlgN | 5.58E-78 |
| AMB_RS19375 | flgK | flagellar hook-associated protein | 1.78E-156 |
| AMB_RS19380 | AMB_RS19380 | flagellin | 1.83E-87 |
| AMB_RS19530 | AMB_RS19530 | EAL domain-containing protein | 6.66E-40 |
| AMB_RS19535 | AMB_RS19535 | phosphorelay signal transduction system | 1.76E-35 |
| AMB_RS19735 | AMB_RS19735 | methyl-accepting chemotaxis protein | 1.03E-47 |
| AMB_RS20310 | AMB_RS20310 | hypothetical protein | 0.00E+00 |
| AMB_RS20510 | AMB_RS20510 | HlyD family type I secretion periplasmic adaptor subunit | 4.55E-45 |
| AMB_RS20740 | AMB_RS20740 | hypothetical protein | 3.62E-46 |

| | | | |
|---|---|---|---|
| AMB_RS21300 | AMB_RS21300 | hypothetical protein | 4.59E-119 |
| AMB_RS21405 | AMB_RS21405 | portal protein | 4.06E-54 |
| AMB_RS21450 | AMB_RS21450 | acyl-CoA carboxylase subunit beta | 1.10E-44 |
| AMB_RS21755 | AMB_RS21755 | hypothetical protein | 6.41E-34 |
| AMB_RS21925 | AMB_RS21925 | DUF4198 domain-containing protein | 3.01E-38 |
| AMB_RS21950 | AMB_RS21950 | hypothetical protein | 2.06E-105 |
| AMB_RS22010 | AMB_RS22010 | hypothetical protein | 3.37E-61 |
| AMB_RS22060 | AMB_RS22060 | helix-turn-helix transcriptional regulator | 1.65E-51 |
| AMB_RS22265 | AMB_RS22265 | glucosaminidase domain-containing protein | 4.85E-34 |
| AMB_RS22695 | AMB_RS22695 | hypothetical protein | 1.63E-49 |
| AMB_RS23255 | AMB_RS23255 | AcyC Adenylate cyclase | 8.28E-131 |
| AMB_RS23360 | AMB_RS23360 | methyl-accepting chemotaxis protein | 3.40E-81 |
| AMB_RS23390 | AMB_RS23390 | VCBS domain-containing protein | 2.40E-54 |
| AMB_RS23595 | AMB_RS23595 | phosphorelay signal transduction system | 1.03E-44 |
| AMB_RS23650 | AMB_RS23650 | hypothetical protein | 7.67E-293 |
| AMB_RS24015 | AMB_RS24015 | tetrahydromethanopterin S-methyltransferase subunit A | 1.28E-52 |
| AMB_RS24485 | AMB_RS24485 | DUF1465 family protein | 3.68E-34 |
| AMB_RS24535 | AMB_RS24535 | hemerythrin domain-containing protein | 1.36E-34 |
| AMB_RS25020 | AMB_RS25020 | GGDEF domain-containing protein | 9.37E-51 |
| AMB_RS25595 | AMB_RS25595 | VCBS domain-containing protein | 1.35E-92 |
| AMB_RS25735 | AMB_RS25735 | hypothetical protein | 0.00E+00 |
| AMB_RS25810 | AMB_RS25810 | VCBS domain-containing protein | 9.64E-125 |
| AMB_RS26320 | AMB_RS26320 | VCBS domain-containing protein | 2.13E-64 |
| AMB_RS26330 | AMB_RS26330 | VCBS domain-containing protein | 5.75E-121 |
| AMB_RS26410 | AMB_RS26410 | VCBS domain-containing protein | 3.39E-137 |
| AMB_RS26745 | AMB_RS26745 | hypothetical protein | 1.44E-93 |

**Table S4.** Detailed list of genes discussed in Figure 3.

| Category | Sub-group | Locus tag | Gene name | Known function |
|---|---|---|---|---|
| **Oxidative stress response & regulation** | Sigma factors | AMB_RS17600 | *rpoE* | RNA polymerase extracytoplasmic-stress sigma factor (sigma-E) |
| | | AMB_RS03585 | *rpoD* | RNA polymerase primary sigma factor 70 |
| | Regulators | AMB_RS01265 | *oxyR-like* | Peroxide-stress-responsive transcriptional regulator (OxyR family, LysR-type) |
| | ROS defense mechanisms | AMB_RS18360 | *gpx* | Glutathione peroxidase |
| | | AMB_RS14995 | *sodC* | Superoxide dismutase |
| | | AMB_RS08430 | *gor* | Glutathione reductase |
| | | AMB_RS00045 | *trxA* | Thioredoxin |
| | | AMB_RS16195 | *ftsH* | ATP-dependent zinc metalloprotease FtsH |
| **Energy production** | ATP synthase | AMB_RS20210 | *atpB* | F1F0 ATP synthase, subunit a (membrane sector) |
| | | AMB_RS20215 | *atpH* | F1F0 ATP synthase, subunit delta/epsilon paralog |
| | | AMB_RS20220 | *atpG* | F1F0 ATP synthase, subunit gamma paralog |
| | | AMB_RS20225 | *atpG (copy)* | F1F0 ATP synthase, subunit gamma paralog |
| | | AMB_RS20930 | *atpC* | F1F0 ATP synthase, subunit epsilon |
| | | AMB_RS20935 | *atpD* | F1F0 ATP synthase, subunit beta |
| | | AMB_RS20945 | *atpA* | F1F0 ATP synthase, subunit alpha |
| | | AMB_RS20950 | *atpH (copy)* | F1F0 ATP synthase, subunit delta/epsilon paralog |
| **Metabolism & cell biology (anabolism)** | Pentose phosphate pathway | AMB_RS11810 | *rpiB* | Ribose-5-phosphate isomerase B |
| | | AMB_RS06980 | *rpe* | Ribulose-phosphate 3-epimerase |
| | | AMB_RS06990 | *tktA* | Transketolase |
| | Cell cycle & genetic informations | AMB_RS15795 | *rpsL* | 30S ribosomal protein S12 |
| | | AMB_RS15820 | *rplL* | 50S ribosomal protein L7/L12 |
| | | AMB_RS22165 | *pheS* | Phenylalanine-tRNA ligase, alpha subunit |
| | | AMB_RS19505 | *ftsZ* | Cell division protein FtsZ (tubulin homolog) |
| | | AMB_RS19500 | *ftsA* | Cell division protein FtsA (actin homolog) |
| | | AMB_RS15355 | *parA* | Chromosome/plasmid partitioning ATPase ParA |

| | | | | |
|---|---|---|---|---|
| | | AMB_RS15815 | *rpoB* | DNA-directed RNA polymerase, beta subunit |
| | | AMB_RS03275 | *dnaA* | Chromosomal replication initiator protein DnaA |
| | | AMB_RS12020 | *gyrA* | DNA gyrase, subunit A |
| | Peptidoglycan synthesis | AMB_RS19480 | *murC* | UDP-N-acetylmuramate--L-alanine ligase |
| | | AMB_RS19465 | *murD* | UDP-N-acetylmuramoylalanine--D-glutamate ligase |
| | | AMB_RS19455 | *murF* | UDP-N-acetylmuramoyl-tripeptide--D-alanyl-D-alanine ligase |
| | Fatty acid synthesis | AMB_RS20305 | *accD* | Acetyl-CoA carboxylase, carboxyltransferase beta subunit |
| | | AMB_RS11890 | *fabH* | 3-oxoacyl-ACP synthase III (initiates fatty acid synthesis) |
| | | AMB_RS21370 | *fabI* | Enoyl-ACP reductase |
| | | AMB_RS21270 | *accA* | Acetyl-CoA carboxylase, carboxyltransferase alpha subunit |
| | Nucleotide synthesis | AMB_RS12545 | *pyrH* | Uridylate kinase |
| | | AMB_RS09215 | *pyrG* | CTP synthase |
| | | AMB_RS18020 | *ndk* | Nucleoside diphosphate kinase |
| | | AMB_RS03600 | *carA* | Carbamoyl-phosphate synthase, small subunit |
| | | AMB_RS03605 | *carB* | Carbamoyl-phosphate synthase, large subunit |
| | | AMB_RS10665 | *purQ* | Phosphoribosylformylglycinamidine synthase, glutamine amidotransferase subunit |
| | | AMB_RS10670 | *purL* | Phosphoribosylformylglycinamidine synthase, PurL subunit |
| | | AMB_RS11140 | *purK* | N5-carboxyaminoimidazole ribonucleotide synthase |
| | | AMB_RS11145 | *purE* | N5-carboxyaminoimidazole ribonucleotide mutase |
| | | AMB_RS14330 | *purM* | Phosphoribosylformylglycinamidine cyclo-ligase |
| | | AMB_RS14335 | *purN* | Phosphoribosylglycinamide formyltransferase |
| | | AMB_RS15365 | *purB* | Adenylosuccinate lyase |
| | | AMB_RS16115 | *purA* | Adenylosuccinate synthetase |
| | | AMB_RS18080 | *purU* | Formyltetrahydrofolate deformylase |
| | Translation factors | AMB_RS16890 | *infA* | Translation initiation factor IF-1 |
| | | AMB_RS20580 | *infB* | Translation initiation factor IF-2 |
| | | AMB_RS01725 | *infC* | Translation initiation factor IF-3 |

| | | | | |
|---|---|---|---|---|
| | | AMB_RS15780 | *tuf* | Elongation factor Tu (EF-Tu) |
| | | AMB_RS15855 | *tuf (copy)* | Elongation factor Tu (EF-Tu) paralog |
| | | AMB_RS12550 | *tsf* | Elongation factor Ts (EF-Ts) |
| | | AMB_RS15785 | *fusA* | Elongation factor G (EF-G) |
| | | AMB_RS08175 | *prfA* | Peptide chain release factor 1 (RF-1) |
| | | AMB_RS14145 | *prfB* | Peptide chain release factor 2 (RF-2) |
| | | AMB_RS01375 | *prfC* | Peptide chain release factor 3 (RF-3) |
| | Ribosomal proteins (30S) | AMB_RS03270 | *rpsT* | 30S ribosomal protein S20 |
| | | AMB_RS10605 | *rpsR* | 30S ribosomal protein S18 |
| | | AMB_RS10610 | *rpsF* | 30S ribosomal protein S6 |
| | | AMB_RS10745 | *rpsD* | 30S ribosomal protein S4 |
| | | AMB_RS12555 | *rpsB* | 30S ribosomal protein S2 |
| | | AMB_RS14180 | *rpsU* | 30S ribosomal protein S21 |
| | | AMB_RS15655 | *rpsK* | 30S ribosomal protein S11 |
| | | AMB_RS15660 | *rpsM* | 30S ribosomal protein S13 |
| | | AMB_RS15685 | *rpsE* | 30S ribosomal protein S5 |
| | | AMB_RS15700 | *rpsH* | 30S ribosomal protein S8 |
| | | AMB_RS15705 | *rpsN* | 30S ribosomal protein S14 |
| | | AMB_RS15725 | *rpsQ* | 30S ribosomal protein S17 |
| | | AMB_RS15740 | *rpsC* | 30S ribosomal protein S3 |
| | | AMB_RS15750 | *rpsS* | 30S ribosomal protein S19 |
| | | AMB_RS15775 | *rpsJ* | 30S ribosomal protein S10 |
| | | AMB_RS15790 | *rpsG* | 30S ribosomal protein S7 |
| | | AMB_RS18000 | *rpsA* | 30S ribosomal protein S1 |
| | | AMB_RS20540 | *rpsP* | 30S ribosomal protein S16 |
| | | AMB_RS20815 | *rpsO* | 30S ribosomal protein S15 |
| | | AMB_RS21545 | *rpsA (copy)* | 30S ribosomal protein S1 (paralog) |

| | | | | |
|---|---|---|---|---|
| | | AMB_RS10590 | *rplI* | 50S ribosomal protein L9 |
| | | AMB_RS15645 | *rplQ* | 50S ribosomal protein L17 |
| | | AMB_RS15675 | *rplO* | 50S ribosomal protein L15 |
| | | AMB_RS15690 | *rplR* | 50S ribosomal protein L18 |
| | | AMB_RS15695 | *rplF* | 50S ribosomal protein L6 |
| | | AMB_RS15710 | *rplE* | 50S ribosomal protein L5 |
| | | AMB_RS15715 | *rplX* | 50S ribosomal protein L24 |
| | | AMB_RS15720 | *rplN* | 50S ribosomal protein L14 |
| | | AMB_RS15735 | *rplP* | 50S ribosomal protein L16 |
| | | AMB_RS15745 | *rplV* | 50S ribosomal protein L22 |
| | | AMB_RS15755 | *rplB* | 50S ribosomal protein L2 |
| | | AMB_RS15765 | *rplD* | 50S ribosomal protein L4 |
| | | AMB_RS15770 | *rplC* | 50S ribosomal protein L3 |
| | Ribosomal proteins (50S) | AMB_RS15825 | *rplJ* | 50S ribosomal protein L10 |
| | | AMB_RS15830 | *rplA* | 50S ribosomal protein L1 |
| | | AMB_RS15835 | *rplK* | 50S ribosomal protein L11 |
| | | AMB_RS18005 | *rplM* | 50S ribosomal protein L13 |
| | | AMB_RS20555 | *rplM (copy)* | 50S ribosomal protein L13 (paralog) |
| | | AMB_RS20615 | *rplU* | 50S ribosomal protein L21 |
| | | AMB_RS22170 | *rplT* | 50S ribosomal protein L20 |
| | | AMB_RS01625 | *rpmB* | 50S ribosomal protein L28 |
| | | AMB_RS11880 | *rpmF* | 50S ribosomal protein L32 |
| | | AMB_RS15300 | *rpmE* | 50S ribosomal protein L31 |
| | | AMB_RS15680 | *rpmD* | 50S ribosomal protein L30 |
| | | AMB_RS15730 | *rpmC* | 50S ribosomal protein L29 |
| | | AMB_RS18060 | *rpmG* | 50S ribosomal protein L33 |
| | | AMB_RS20620 | *rpmA* | 50S ribosomal protein L27 |

| | | | | |
|---|---|---|---|---|
| | | AMB_RS22020 | *rpmH* | 50S ribosomal protein L34 |
| | | AMB_RS22175 | *rpmI* | 50S ribosomal protein L35 |
| | Amino acid activation | AMB_RS17865 | *alaS* | Alanine-tRNA ligase |
| | | AMB_RS00145 | *leuS* | Leucine-tRNA ligase |

**Table S5.** List of MIS genes with HMM products.

| Gene id | Product (HMM) | Number of paralog genes |
|---|---|---|
| AMB_RS02010 | hypothetical protein | - |
| AMB_RS02015 | hypothetical protein | - |
| AMB_RS23830 | mamK-like | 2 |
| AMB_RS23835 | McaA | - |
| AMB_RS24750 | McaB | - |
| AMB_RS23840 | transposase | - |
| AMB_RS26010 | hypothetical protein | - |
| AMB_RS02045 | mamD-like | 2 |
| AMB_RS24755 | hypothetical protein | - |
| AMB_RS24760 | hypothetical protein | - |
| AMB_RS26940 | transposase | - |
| AMB_RS26690 | tnpA | 4 |
| AMB_RS02055 | tnpB | 4 |
| AMB_RS02060 | tnpC | 3 |
| AMB_RS26695 | transposase | - |
| AMB_RS23850 | transposase | - |
| AMB_RS02075 | tnpA | 4 |
| AMB_RS02080 | tpnB | 4 |
| AMB_RS02085 | mamL-like | 2 |
| AMB_RS02090 | transposase | - |
| AMB_RS02095 | transposase | - |
| AMB_RS02100 | transposase | - |
| AMB_RS24775 | hypothetical protein | - |
| AMB_RS23855 | mamE-like | 3 |
| AMB_RS02110 | transposase | - |
| AMB_RS23860 | transposase | - |
| AMB_RS02120 | hypothetical protein | - |
| AMB_RS02125 | mamQ-like | 3 |
| AMB_RS23865 | hypothetical protein | - |
| AMB_RS23870 | parA | 9 |
| AMB_RS26700 | hypothetical protein | - |
| AMB_RS02130 | hypothetical protein | - |

**Table S6.** List of MAI genes with HMM products.

| Gene id | Product (HMM) | Number of paralog genes |
|---|---|---|
| AMB_RS24875 | transposase | - |
| AMB_RS26290 | IS110 family transposase | 1 |
| AMB_RS04800 | MbcA/ParS/Xre antitoxin family protein | 0 |
| AMB_RS24880 | hypothetical protein | - |
| AMB_RS04805 | bacteriohemerythrin | 40 |
| AMB_RS24885 | hypothetical protein | - |
| AMB_RS04810 | hypothetical protein | - |
| AMB_RS04815 | FTR1 family iron permease | 2 |
| AMB_RS04820 | 4Fe-4S binding protein | 4 |
| AMB_RS04825 | iron transporter | 4 |
| AMB_RS04830 | iron transporter | 4 |
| AMB_RS04835 | OprB Carbohydrate-selective porin, OprB family | 6 |
| AMB_RS04840 | efflux RND transporter periplasmic adaptor subunit | 18 |
| AMB_RS23935 | hypothetical protein | - |
| AMB_RS04845 | hypothetical protein | - |
| AMB_RS04850 | hypothetical protein | - |
| AMB_RS04855 | cheY,chemotaxis response regulator CheY | 2 |
| AMB_RS04860 | IS110 family transposase | - |
| AMB_RS04865 | hypothetical protein | - |
| AMB_RS04870 | mamC | 0 |
| AMB_RS26015 | mamD | 1 |
| AMB_RS04880 | mamF | 0 |
| AMB_RS04885 | mamG | 1 |
| AMB_RS04895 | mms6 | 0 |
| AMB_RS04900 | mmsF | 0 |
| AMB_RS04905 | hypothetical protein | - |
| AMB_RS04910 | heme biosynthesis protein HemY | 2 |
| AMB_RS04915 | hypothetical protein | - |
| AMB_RS04920 | mamH | 0 |
| AMB_RS04925 | mamI | 0 |
| AMB_RS04930 | mamE | 2 |
| AMB_RS23165 | mamJ | 0 |
| AMB_RS04945 | mamK | 1 |
| AMB_RS04950 | mamL | 1 |
| AMB_RS04955 | mamM | 0 |
| AMB_RS04960 | mamN | 0 |

| | | |
|---|---|---|
| AMB_RS04965 | mamO | 0 |
| AMB_RS04970 | mamP | 0 |
| AMB_RS04975 | mamA | 0 |
| AMB_RS04980 | mamQ | 2 |
| AMB_RS04985 | mamR | 1 |
| AMB_RS04990 | mamB | 1 |
| AMB_RS04995 | mamS | 1 |
| AMB_RS05000 | mamT | 0 |
| AMB_RS05005 | mamU | 1 |
| AMB_RS05010 | mamV | 0 |
| AMB_RS05015 | hypothetical protein | - |
| AMB_RS25790 | hypothetical protein | - |
| AMB_RS05020 | chemotaxis protein CheY | 2 |
| AMB_RS05025 | mamW | 0 |
| AMB_RS05030 | YdaS antitoxin is a family of putative bacterial antitoxins, neutralising the toxin YdaT, family pfam06254. | - |
| AMB_RS05035 | hypothetical protein | - |
| AMB_RS05040 | hypothetical protein | - |
| AMB_RS05045 | bacteriohemerythrin | 40 |
| AMB_RS05050 | hypothetical protein | - |
| AMB_RS23945 | hypothetical protein | - |
| AMB_RS05070 | HU family DNA-binding protein" | 8 |
| AMB_RS05075 | MucR family transcriptional regulator | 9 |
| AMB_RS05080 | hypothetical protein | - |
| AMB_RS05085 | methyl-accepting chemotaxis protein | 60 |
| AMB_RS05090 | bacteriohemerythrin | 40 |
| AMB_RS05095 | bacteriohemerythrin | 40 |
| AMB_RS23950 | hypothetical protein | - |
| AMB_RS05100 | methyl-accepting chemotaxis protein | 60 |
| AMB_RS05105 | PAS domain-containing protein | 8 |
| AMB_RS05110 | Hpt domain-containing protein | 1 |
| AMB_RS23955 | bacteriohemerythrin | 40 |
| AMB_RS05120 | YjbI Uncharacterized protein YjbI, contains pentapeptide repeats | - |
| AMB_RS05125 | hypothetical protein | - |
| AMB_RS05130 | hypothetical protein | - |
| AMB_RS05135 | hypothetical protein | - |
| AMB_RS05140 | mamE | 0 |
| AMB_RS05145 | FraH protein | - |
| AMB_RS05150 | DegQ family serine endoprotease | - |

| | | |
|---|---|---|
| AMB_RS05155 | mamQ | 2 |
| AMB_RS05160 | mamR | 1 |
| AMB_RS05165 | mamB | 1 |
| AMB_RS05170 | hypothetical protein | - |
| AMB_RS05175 | Fur family transcriptional regulato | 5 |
| AMB_RS05180 | MerR family transcriptional regulator | 2 |
| AMB_RS05185 | heavy metal translocating P-type ATPase | 3 |
| AMB_RS05190 | CDF family Co(II)/Ni(II) efflux transporter DmeF | 1 |
| AMB_RS05195 | hypothetical protein | |
| AMB_RS05200 | ftsZ | 1 |
| AMB_RS05205 | mamZ | 0 |
| AMB_RS05210 | mamX | 0 |
| AMB_RS05215 | mamY | 0 |
| AMB_RS26295 | hypothetical protein | - |
| AMB_RS05220 | hypothetical protein | - |
| AMB_RS05225 | hypothetical protein | - |
| AMB_RS05230 | hypothetical protein | - |
| AMB_RS26945 | transposase | - |
| AMB_RS26020 | carbohydrate porin | 3 |
| AMB_RS05240 | feoA | 1 |
| AMB_RS05245 | feoB | 1 |
| AMB_RS25570 | hypothetical protein | - |
| AMB_RS05250 | hypothetical protein | - |
| AMB_RS24925 | mamG | 2 |
| AMB_RS05260 | hypothetical protein | - |
| AMB_RS05265 | PS1 Serine/threonine protein kinase | 0 |
| AMB_RS24930 | hypothetical protein | - |
| AMB_RS05270 | HEAT repeat domain-containing protein | 2 |
| AMB_RS25575 | HEAT repeat domain-containing protein | 2 |
| AMB_RS23975 | IS3 family transposase | 12 |
| AMB_RS23980 | helix-turn-helix domain-containing protein | 42 |
| AMB_RS05275 | IS3 family transposase | 12 |

**Table S7.** XAS parameters ($L_3$-a and $L_3$-b peak positions and splitting, and $L_3$-b/$L_3$-a intensity ratios) of magnetite, bacterial and reference samples.

| Sample | $L_3$-a (eV) | $L_3$-b (eV) | $L_3$-b/$L_3$-a splitting (eV) | $L_3$-b/$L_3$-a intensity ratio |
|---|---|---|---|---|
| Wild-type AMB-1 | 706.261 | 707.698 | 1.437 | 2.050 |
| ΔMIS AMB-1 | 706.256 | 707.690 | 1.434 | 1.780 |
| Magnetite | 706.251 | 707.641 | 1.390 | 1.862 |
| Maghemite | 706.225 | 707.675 | 1.450 | 2.101 |
| Ferrihydrite | 706.253 | 707.723 | 1.470 | 1.776 |

**Table S8.** OD, $OD_{max}$ and $OD_{min}$ values measured for calculation of $C_{mag}$ coefficients in the wild-type and ΔMIS AMB-1 strains.

| Sample | Time (h) | OD | $OD_{max}$ | $OD_{min}$ |
|---|---|---|---|---|
| **Wild-type (1st replicate)** | 0 | 0.060 | 0.060 | 0.060 |
| | 2.25 | 0.103 | 0.107 | 0.102 |
| | 4.25 | 0.127 | 0.135 | 0.12 |
| | 8 | 0.236 | 0.324 | 0.194 |
| | 9 | 0.287 | 0.416 | 0.238 |
| **Wild-type (2nd replicate)** | 0 | 0.072 | 0.072 | 0.072 |
| | 2.25 | 0.110 | 0.113 | 0.108 |
| | 4.25 | 0.136 | 0.145 | 0.128 |
| | 8 | 0.261 | 0.356 | 0.209 |
| | 9 | 0.298 | 0.436 | 0.245 |
| **Wild-type (3rd replicate)** | 0 | 0.026 | 0.026 | 0.026 |
| | 2.25 | 0.068 | 0.072 | 0.065 |
| | 4.25 | 0.096 | 0.110 | 0.087 |
| | 8 | 0.213 | 0.310 | 0.154 |
| | 9 | 0.259 | 0.390 | 0.203 |
| **ΔMIS (1st replicate)** | 0 | 0.110 | 0.011 | 0.011 |
| | 2.25 | 0.036 | 0.038 | 0.033 |
| | 4.25 | 0.055 | 0.065 | 0.048 |
| | 8 | 0.111 | 0.164 | 0.079 |
| | 9 | 0.133 | 0.222 | 0.095 |
| **ΔMIS (2nd replicate)** | 0 | 0.016 | 0.016 | 0.016 |
| | 2.25 | 0.040 | 0.043 | 0.036 |
| | 4.25 | 0.057 | 0.073 | 0.045 |
| | 8 | 0.125 | 0.200 | 0.083 |
| | 9 | 0.142 | 0.254 | 0.107 |
| **ΔMIS (3rd replicate)** | 0 | 0.036 | 0.036 | 0.036 |
| | 2.25 | 0.054 | 0.060 | 0.051 |
| | 4.25 | 0.078 | 0.101 | 0.064 |
| | 8 | 0.143 | 0.223 | 0.102 |
| | 9 | 0.152 | 0.266 | 0.112 |
